# Computational design of orthogonal TCR α/β interfaces for dual-TCR therapeutics

**DOI:** 10.64898/2026.01.02.697397

**Authors:** Tomoaki Kinjo, Shawn Yu, Nathan Nicely, Andrew Leaver-Fay, William Y. Kim, Brian Kuhlman

## Abstract

T-cell receptors (TCRs) recognize peptides presented by MHC, enabling access to intracellular targets that are largely inaccessible to antibodies and difficult to target with small molecules.

Despite this potential, their inherent cross-reactivity limits tumor specificity, while single-antigen targeting provides limited coverage of intratumoral heterogeneity. Dual-TCR therapeutics comprising two distinct TCRs could enhance tumor specificity via combinatorial recognition while broadening coverage across heterogeneous antigens. However, practical development of dual-TCR therapeutics has been limited by α/β subunit mispairing that prevents efficient production and creates undesired binding properties. Here, we develop orthogonal TCR α/β interfaces that prevent subunit mispairing. Using computational multistate design and second-site suppressor strategies implemented in Rosetta, we identified over 250 TCR variants for experimental screening to assess protein stability and pairing fidelity. The top-performing designs achieved approximately 95% correct pairing, as validated by mass spectrometry and X-ray crystallography. Focusing mutations on constant domains and conserved framework regions of variable domains enabled broad applicability across diverse TCRs while preserving antigen recognition. Using these orthogonal interfaces, we developed trispecific T-cell engagers (TriTEs) that target two cancer-testis antigens and CD3 on T cells, demonstrating enhanced potency under dual-antigen engagement (EC50 of 380 fM) while maintaining high activity when targeting cells displaying a single antigen (EC50s of 48 pM and 20 pM). This orthogonal TCR interface technology establishes a generalizable platform for engineering multi-specific immune therapeutics targeting diverse cancer antigens.

## Introduction

T-cell receptors (TCRs) enable recognition of peptides derived from intracellular proteins presented by major histocompatibility complex (MHC) molecules on the cell surface^1,2^, providing therapeutic access to targets inaccessible to antibodies and small molecules^3–5^. Dual-TCR therapeutics, which incorporate two different TCRs in a single molecule, could enhance tumor specificity through combinatorial recognition, while providing broader coverage of tumors with heterogeneous antigen expression. However, practical development of dual-TCR therapeutics has been hindered by a long-standing challenge of subunit mispairing. Each TCR consists of an α and β chain, and co-expression of two distinct TCRs produces two α and two β chains that can pair indiscriminately, generating misassembled byproducts that reduce production yield and pose risks of undesired binding specificities causing off-target toxicities such as graft-versus-host disease^6–8^.

Previous efforts to address TCR mispairing, including introduction of interchain disulfide bonds^9^, knob-into-hole modifications^10^, and murinization of constant domains^11,12^, have been insufficient to practically prevent mispairing^13,14^. As an alternative strategy, domain-swapped TCRs prevent signaling by mispaired TCRs by gating CD3 assembly through swapping the connecting peptide (CP) and transmembrane (TM) segments between the α and β chains, but they do not directly prevent mispairing at the extracellular α/β interface of the variable and constant domains^15^. Knockout or knockdown of endogenous TCR genes^16,17^, while effective for single-TCR cell-based therapies, cannot prevent mispairing between two exogenous TCRs.

Moreover, previous approaches have largely relied on assessing TCR expression on the cell surface or T-cell functional readouts, rather than quantitative biophysical characterization of α/β interface stability and pairing fidelity. In cellular assays, apparent reductions in mispairing can be influenced by differences in expression between endogenous and transgenic TCRs, as well as by CD3 assembly and signaling, making it difficult to quantitatively measure pairing fidelity attributable to α/β interface orthogonality. This distinction is critical, because expression-based predominance does not ensure correct assembly between two engineered TCRs expressed at comparable levels, where pairing specificity is instead determined by interface orthogonality.

Computational protein design has enabled rational engineering of protein-protein interfaces with quantifiable properties. The Rosetta macromolecular modeling suite employs an energy function based on physical and statistical potentials^18–20^, and has been used in structure prediction, antibody engineering, and de novo protein design^21,22^. Previous work has applied Rosetta to engineer orthogonal interfaces in bispecific antibodies by introducing complementary mutations that favor desired pairings while disfavoring undesired combinations^23–25^. Thus, we hypothesized that computational design combined with quantitative biophysical characterization could address the TCR mispairing problem by engineering orthogonal α/β interfaces.

Here, we present the computational design and experimental validation of orthogonal TCR α/β interfaces that prevent subunit mispairing (**Fig. 1**). We systematically explored mutations across the TCR α/β interface using multistate design (MSD)^26^ and second-site suppressor (SSS) strategies^27^ implemented in Rosetta, generating over one million design models. Focusing mutations on constant domains and conserved framework regions of variable domains^28^ enabled broad applicability across diverse TCR sequences. We experimentally characterized more than 250 designs encompassing over 750 pairing states using quantitative biophysical analysis of interface stability and pairing fidelity, identifying highly orthogonal pairs validated by mass spectrometry and X-ray crystallography. Using these orthogonal interfaces, we developed trispecific T-cell engagers (TriTEs) comprising two TCRs and an anti-CD3ε domain. TriTEs targeting two cancer-testis antigens demonstrated enhanced cytotoxicity upon dual-antigen engagement, while retaining activity against cells with either antigen alone. Our integrated design approach enables correctly assembled dual-TCR therapeutics that enhance tumor specificity and improve antigen coverage.

**Figure 1.**
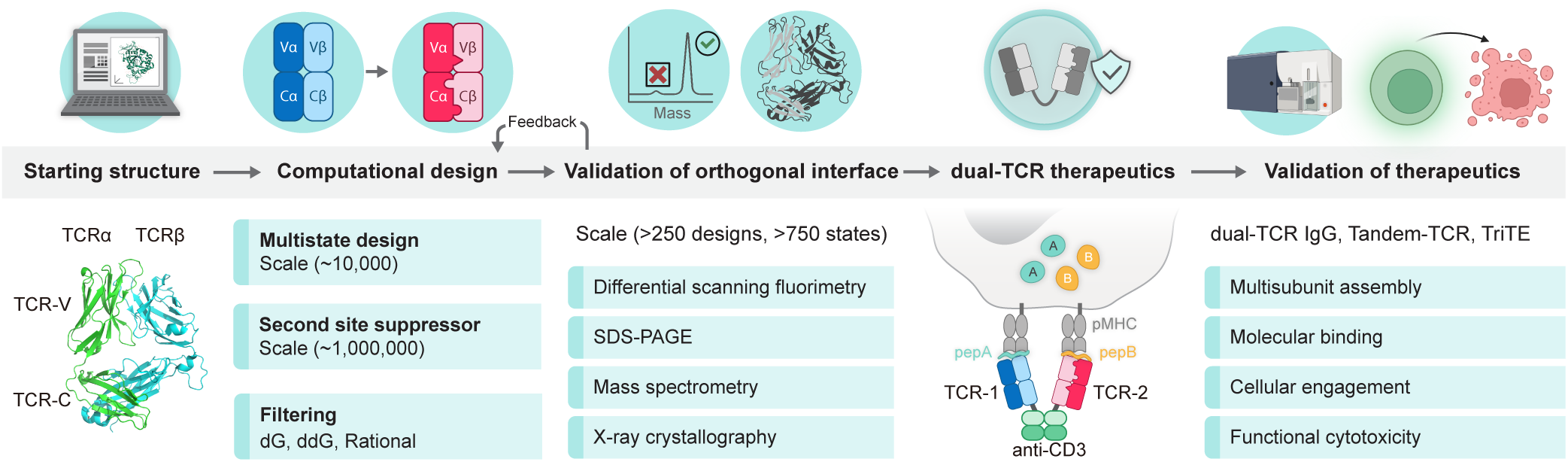
Overall design and validation pipeline for engineering orthogonal TCR interfaces and dual-TCR therapeutics. Schematic overview of the integrated computational and experimental workflow used to engineer orthogonal TCR α/β interfaces and assemble dual-TCR therapeutics. The design concept is to favor correct α/β pairing while actively destabilizing mispaired subunit combinations at the shared interface, thereby preventing assembly of off-target TCRs. Starting from TCR variable (TCR-V) and constant (TCR-C) domain structures, orthogonal interfaces were designed using multistate design (MSD) and second-site suppressor strategies. Candidate designs were filtered based on interface energetics (ΔG and ΔΔG), followed by systematic experimental validation of more than 250 designs encompassing over 750 pairing states using nano-differential scanning fluorimetry (nanoDSF), SDS-PAGE, intact mass spectrometry, and X-ray crystallography. Validated orthogonal interfaces were subsequently incorporated into dual-TCR therapeutic formats, which were evaluated by SDS-PAGE, mass spectrometry, surface plasmon resonance (SPR), flow cytometry, and T-cell-mediated tumor cell cytotoxicity assays.

## Results

### Computational design of orthogonal TCR α/β interfaces

The design goal for orthogonal TCRs is to make mutations at the interface of the TCR α and β chains so that the mutants interact with each other but not with their wild-type partners (**Fig. 2a**). To identify such mutations, we used a multistate design (MSD) protocol in Rosetta^25,26,29^. MSD enables a single protein sequence to satisfy multiple design objectives by simultaneously optimizing across different structural states^24,25,29^. We applied MSD to engineer orthogonal TCR α/β interfaces by optimizing sequences to favor MT/MT pairing while penalizing MT/WT and WT/MT pairing through differential interface scoring. High-scoring designs were subjected to energy-based conformational sampling (Rosetta’s FastRelax protocol) to remove false-positive clashes, and for selected clusters, the resulting ensemble of structures was used for additional rounds of MSD to sample diverse backbone conformations (**Fig. 2b**). We used starting structures comprising a previously-described stabilized TCR constant (stTCR-C) domain (PDB: 6U07)^30^ and a TCR variable (TCR-V) domain (PDB: 2F53)^31^. As MSD is computationally intensive, we selected small clusters of interface residues for each simulation by structure-guided visual inspection. We restricted designs to the TCR-C and the framework region of the TCR-V that are highly conserved in diverse TCRs^28^. Selected clusters comprised residues forming various types of interactions, including hydrogen bonds, salt bridges, and cation-π interactions in TCR-C, and a conserved Gln pair in the TCR-V framework region (**Fig. 2c**). The complete set of residue clusters used for MSD is provided in **Supplementary Table S1**. MSD was applied to each cluster independently as described above. Top-scoring designs showed strong destabilization of MT/WT and WT/MT off-target pairs relative to the MT/MT configuration (**Fig. 2d**). The corresponding MSD scores across all interface clusters are provided in **Supplementary Fig. S1**. Analysis of mean adjusted interface energy differences between on-target and off-target pairing states across inter-chain contacting residue pairs identified key positions in the interface that contribute to orthogonal pairing (**Fig. 2e and Supplementary Fig. S2**).

**Figure 2.**
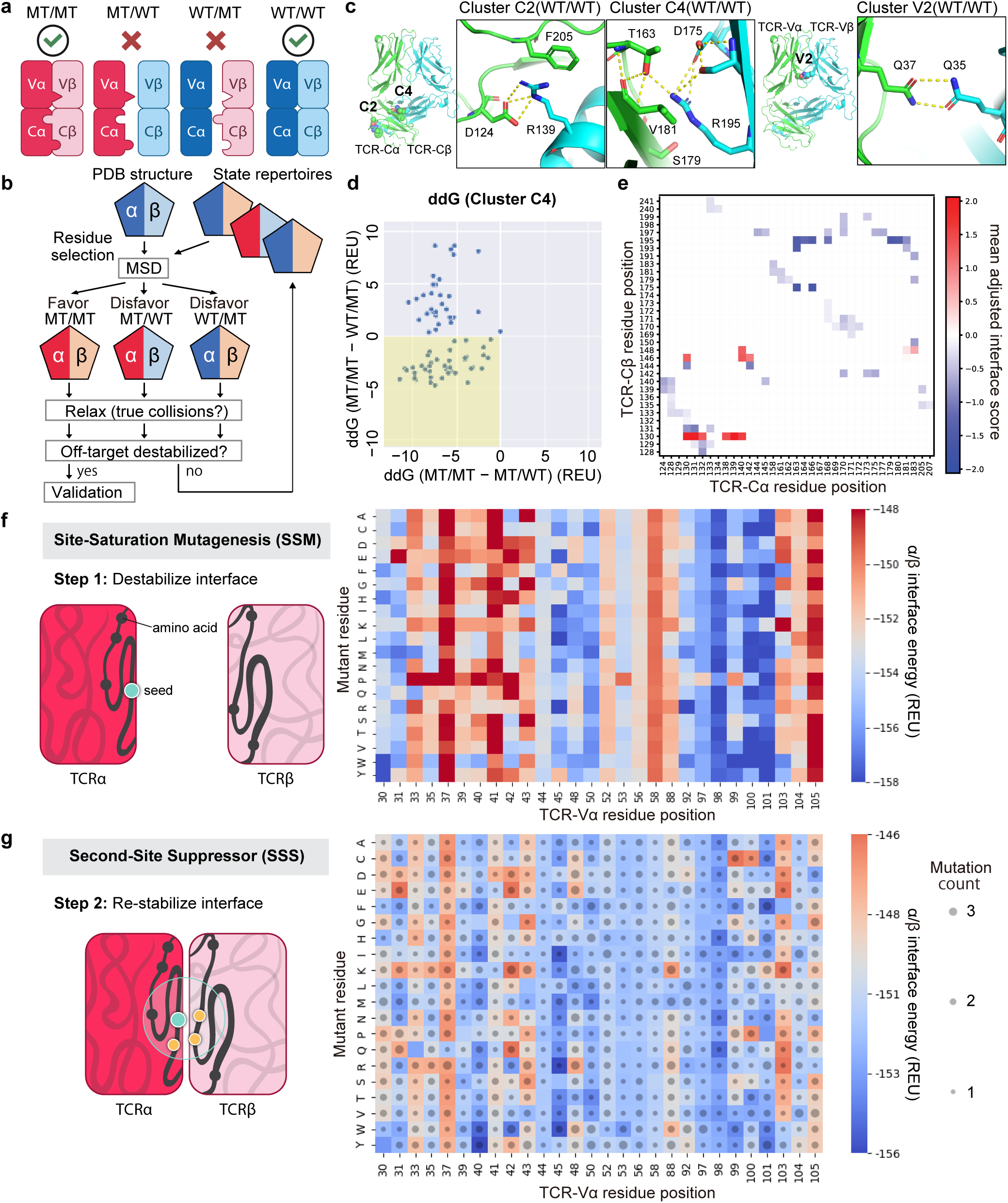
Computational design of orthogonal TCR α/β interfaces. **a**, Design goal for orthogonal TCR α/β interfaces and the four possible pairing states formed upon co-expression of subunits with wild-type (WT) or mutant (MT) interfaces. Pairing states are defined as MT/MT (mutant α + mutant β), MT/WT (mutant α + wild-type β), WT/MT (wild-type α + mutant β), and WT/WT (wild-type α + wild-type β). MT/MT represents the on-target design state, MT/WT and WT/MT represent off-target mixed pairing states, and WT/WT serves as the wild-type reference state. **b**, Multistate design (MSD) framework used to engineer orthogonal TCR α/β interfaces. A single TCR α/β sequence was optimized across multiple structural states, with the desired MT/MT pairing treated as a positive design state and the mixed MT/WT and WT/MT pairings treated as negative states. High-scoring designs were subsequently subjected to structural relaxation to remove false-positive steric clashes. For selected residue clusters, the resulting relaxed structures were used as state repertoires in additional rounds of MSD to sample diverse backbone conformations prior to experimental validation. **c**, Structure-guided selection of representative residue clusters used for MSD calculations at the TCR constant-domain interface and the conserved framework region of the variable-domain interface, focusing on inter-chain contacts while excluding CDR loops to preserve antigen recognition. **d**, Representative MSD interface energy scores (ΔΔG) for a constant-domain residue cluster, showing interface energy differences of off-target pairings (MT/WT and WT/MT) relative to the on-target MT/MT state. **e**, Residue-residue map of mean adjusted interface energy differences between on-target and off-target pairing states projected onto inter-chain contacting TCR α/β residue pairs, with colder colors indicating residue pairs for which off-target pairings are energetically less favorable relative to the on-target state. **f**, Schematic overview of in silico site-saturation mutagenesis (SSM) scanning of the TCR α/β interface. The accompanying map displays α/β interface energies (dG_separated in Rosetta energy units (REU)) for single-point mutations across inter-chain contacting residues for a representative Vα interface. **g**, Schematic overview of second-site suppressor (SSS) screening in which destabilizing single-site mutations are used as seeds to identify compensatory mutations on the partner chain. The accompanying map displays α/β interface energies (dG_separated in REU) for a representative Vα interface, with dot size indicating the number of mutations in the redesign.

Although MSD delivered highly orthogonal solutions, it is computationally expensive and sequence exploration is focused on pre-selected residue clusters, leaving much of the interface unsampled. To achieve more comprehensive coverage across all interface positions, we adapted a two-stage second-site suppressor (SSS) strategy (**Fig. 2f,g**). First, we used Rosetta to perform in-silico site-saturation mutagenesis (SSM)^32^ across all interfacial positions on the α and β chain (**Fig. 2f**). Each single-mutant model was evaluated for its effect on interface stability using the dG_separated score obtained with Rosetta’s InterfaceAnalyzer protocol^33^. Subsequently, each SSM mutation was treated as a seed mutation and processed by a mutation cluster workflow adapted for interface design, in which the FastDesign protocol^34^ stochastically samples sequence and rotamer space within a compact interfacial shell around the seed mutation (**Fig. 2g**). Designs where compensatory mutations on the opposite chain rescued the destabilization caused by the seed mutation were identified as candidates for orthogonal pairing. The mutation cluster search employed asymmetric native-retention biases, restricting mutations to residues proximal to the seed mutation while promoting compensatory mutations on the opposite chain to achieve orthogonal pairing. Designs were filtered by scores such as total_score, dG_separated, and number of mutations. Comprehensive SSM and SSS scoring results for TCR-Cα, TCR-Cβ, and TCR-Vβ are shown in **Supplementary Fig. S3**.

We also incorporated rational design based on trends observed across MSD, SSM, and SSS analyses. Key residues identified through SSM analysis informed the selection of additional MSD clusters. These hybrid designs complemented the automated searches by expanding the accessible sequence space guided by structural insights.

Collectively, MSD provided deep optimization within computationally accessible clusters, while the SSM/SSS strategy systematically scanned the remainder of the interface to identify compensatory pairs at a fraction of the computational cost per position. From over a million models generated through these computational design pipelines, we advanced to experimental characterization.

### Experimental validation of designed orthogonal TCR α/β interfaces

We experimentally characterized more than 250 designs (encompassing over 750 pairing states), including 128 variants in the constant domains (TCR-C), 73 in the variable domains (TCR-V), and over 50 combination designs. Given that TCRs are unstable and prone to aggregate^35,36^ when not anchored to the membrane, we used a stabilized TCR scaffold (stTCR) as previously described^30^. Soluble TCR constructs comprising the extracellular domains of α and β chains (truncated at Cα-C213 and Cβ-C247; construct details in Methods) were co-expressed in Expi293F cells as MT/MT, MT/WT, and WT/MT combinations, and purified by Ni-NTA affinity chromatography. Expression levels were assessed by SDS-PAGE with equal amounts of protein lysate from the different samples. The SDS-PAGE of TCRs showed glycosylation-dependent smear patterns characteristic of TCRs^30^. To assess α/β interface stability, we measured thermal unfolding using nano-differential scanning fluorimetry (nanoDSF). The unfolding transition exhibited a single cooperative melting peak (**Supplementary Fig. S4**), reflecting the coupled denaturation of α and β subunits^30^. Comprehensive analyses of an extensive panel of designed constructs, including desC1–128, combiC1–48, desV1–73, and desVcombiC variants, are presented in **Supplementary Figs. S5–6**, with the corresponding design sets listed in **Supplementary Tables S2–4**. Representative examples are described below, and a subset of experimentally validated designs exhibiting favorable properties across assays is summarized in **Supplementary Tables S5–S7**.

Among the TCR-C designs, Design C127 (desC127) employs a charge-swap strategy where Cα-D124 and Cα-F205 are mutated to Arg and Lys, respectively, while Cβ-R139 is mutated to Glu (**Fig. 3a**, **top**). In the MT/MT state, the reversed charge distribution maintains favorable electrostatic interactions, whereas MT/WT and WT/MT combinations create repulsive charge that destabilize the interface. Design C43 (desC43) exemplifies a knob-into-hole approach where Cβ-R195 is mutated to Ser and Cα-S179 is mutated to Arg (**Fig. 3a**, **middle**). In the MT/MT state, the mutant Arg forms stabilizing H-bonds, while in the MT/WT state, electrostatic repulsions between two Arg residues destabilize the interface, and in the WT/MT pair, the interface is underpacked. NanoDSF analysis revealed that desC43 destabilized the MT/WT and WT/MT mispaired states by 7 °C and 5 °C, respectively. SDS-PAGE showed substantially reduced expression of both mispaired states, with a more pronounced reduction for MT/WT. Because interface mutations were designed in spatially distinct residue clusters across the TCR α/β interface, we next tested whether combining mutations from separate clusters into a single construct could further enhance orthogonality. Combining desC127 and desC43 to generate a combinatorial design (combiC46) substantially enhanced orthogonality, with nanoDSF showing destabilization of MT/WT and WT/MT by 7 °C and 11 °C, respectively. SDS-PAGE revealed substantially reduced expression of both mispaired states, with MT/WT expression below detection limits (**Fig. 3b**). To benchmark our designs against previous approaches, we similarly performed nanoDSF and SDS-PAGE analysis of human (huC) and murine TCR-C (muC) pairs, as it is a commonly used strategy to prevent TCR mispairing. Both huC and muC exhibited lower stability and expression than the stabilized TCR constant domain (stTCR-C) used as the starting point for our design process. Despite the widespread use of murine TCR-C, the mispaired huCα/muCβ pair was more abundant than either of the correctly paired huCα/huCβ or muCα/muCβ complexes, indicating substantial mispairing (**Fig. 3c**).

**Figure 3.**
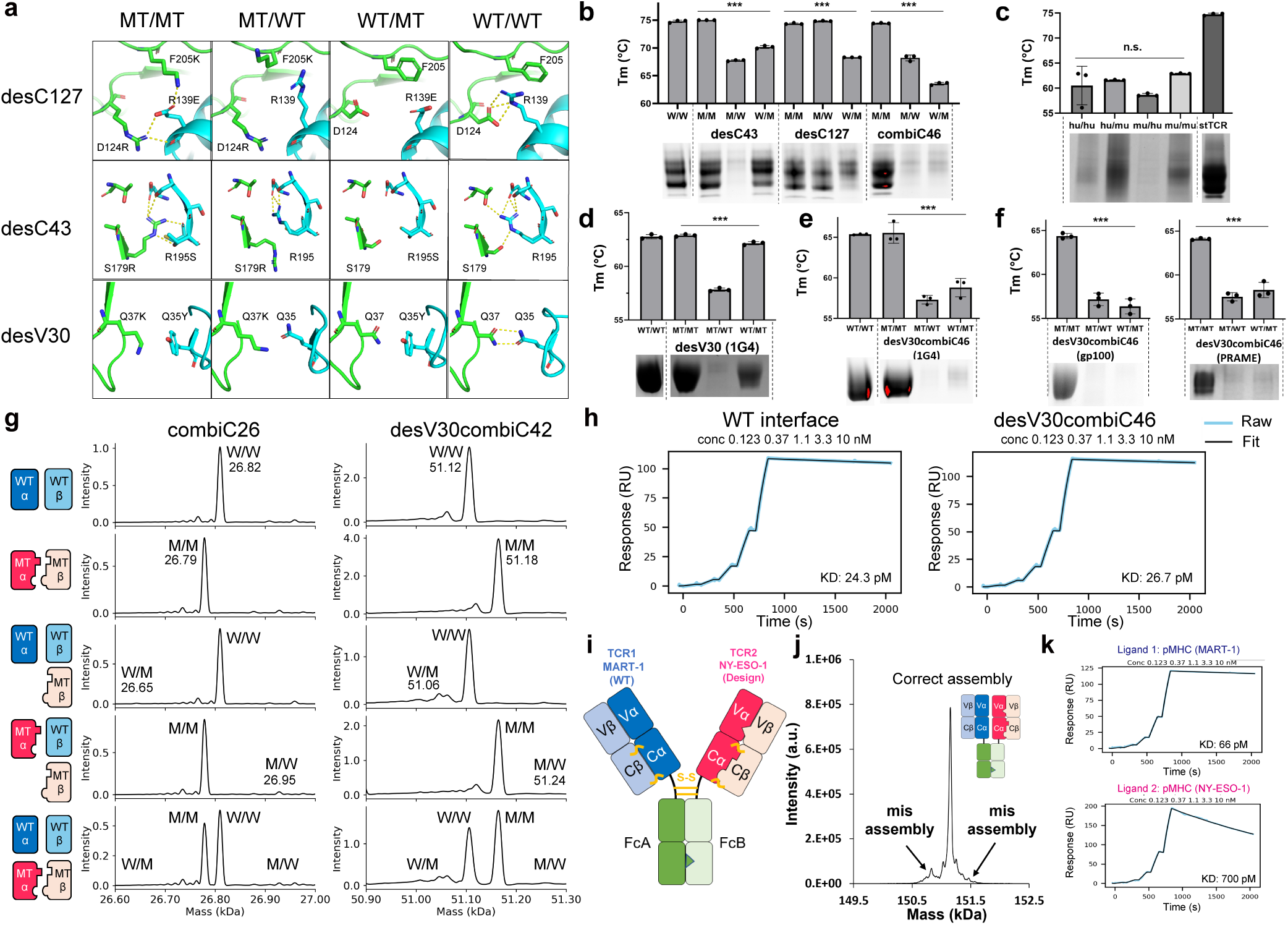
Experimental validation of designed orthogonal TCR α/β interfaces. **a**, Representative Rosetta models of constant-domain designs (desC127 and desC43) and a variable-framework design (desV30), shown across MT/MT, MT/WT, WT/MT, and WT/WT pairing states. **b**, Thermal stability measured by nano-differential scanning fluorimetry (nanoDSF) and SDS-PAGE analysis of desC43, desC127 and the combined constant-domain design combiC46 across pairing states. **c**, Human (hu) and murine (mu) constant-domain pairing (hu/hu, hu/mu, mu/hu and mu/mu), assessed by nanoDSF and SDS-PAGE. **d**, nanoDSF and SDS-PAGE analysis of the variable-framework design desV30 introduced into the NY-ESO-1-specific 1G4 TCR. **e**, nanoDSF and SDS-PAGE analysis of the combined variable and constant-domain design desV30combiC46 in the 1G4 TCR. **f**, Applicability of desV30combiC46 to additional TCRs targeting gp100 and PRAME, assessed by nanoDSF and SDS-PAGE. Bars show mean ± s.d. (n = 3 replicates from 2–3 independent experiments). Statistical analysis was performed by one-way ANOVA; ***P < 0.001; n.s., not significant. **g**, Subunit competition assays quantified by intact mass spectrometry following co-expression of wild-type and mutant TCR α and β chains. The left schematic summarizes five expression conditions, including two reference conditions (WT and MT α/β pairs) and three competitive conditions in which WT and MT subunits are co-expressed to assess preferential α/β pairing. Representative intact-mass spectra are shown for combiC26 and desV30combiC42. Peaks corresponding to α/β pairing states are annotated as W/W, M/M, W/M and M/W (W: WT, M: MT), and the indicated masses correspond to the expected molecular weights of each pairing state. **h**, Surface plasmon resonance (SPR) single-cycle kinetics comparing a wild-type interface and a designed interface (desV30combiC46) for binding to cognate pMHC ligands (five 1:3 serial dilutions, 0.123–10 nM). **i**, Schematic representation of a bispecific TCR-IgG (bsTCR-IgG) incorporating two orthogonal TCRs, TCR1 specific for MART-1 with a wild-type α/β interface and TCR2 specific for NY-ESO-1 with a designed orthogonal interface, fused to a heterodimeric Fc. **j**, Intact-mass spectrometry analysis of bsTCR-IgG showing predominant correctly assembled species together with minor misassembled species. **k**, Surface plasmon resonance (SPR) single-cycle kinetics measuring binding of bsTCR-IgG to MART-1 and NY-ESO-1 pMHC ligands (five 1:3 serial dilutions, 0.123–10 nM).

Within the TCR-V designs, Design V30 (desV30) introduces a compensatory cation-π interaction by mutating a conserved Gln pair (Vα-Q37 and Vβ-Q35) to Lys and Tyr (**Fig. 3a**, **bottom**). In the correctly paired MT/MT complex, Lys-Tyr cation-π pairing is expected to stabilize the interface, whereas in mispaired complexes disruption of the conserved Gln-Gln interaction and, in the MT/WT state, an unpaired Lys charge are expected to be destabilizing. Consistent with these expectations, nanoDSF showed that desV30 destabilized the MT/WT complex by approximately 6 °C while preserving the stability of the MT/MT complex, and SDS-PAGE indicated reduced expression for both mispaired states, with a more pronounced reduction for MT/WT (**Fig. 3d**). This Gln pair is conserved in approximately 90% of TRAV and TRBV sequences in the ImMunoGeneTics (IMGT) database^37^, as illustrated by sequence logo analysis (**Supplementary Fig. S7**), suggesting that the desV30 mutation set is broadly applicable across diverse TCR repertoires. Combining desV30 with combiC46 (desV30combiC46) further enhanced orthogonality across both TCR-C and TCR-V interfaces, as confirmed by nanoDSF and SDS-PAGE. We also confirmed applicability with multiple TCRs, including TCRs targeting NY-ESO-1 (1G4)^31^, gp100^38^, and PRAME^39^ (**Fig. 3e,f**). Framework annotations for all TCRs used throughout this study are summarized in **Supplementary Table S8**.

To further assess pairing fidelity under competitive conditions, we performed subunit competition assays quantified by intact-mass spectrometry (**Fig. 3g**), with the corresponding calculated expected mass values summarized in **Supplementary Table S9**. TCR α and β chains were co-expressed under defined conditions, including two non-competitive reference conditions (WT-only and MT-only α/β pairs) and three competitive conditions: 1) WTα + WTβ + MTβ, 2) MTα + WTβ + MTβ, and 3) WTα + WTβ + MTα + MTβ, enabling direct evaluation of preferential α/β pairing based on the expected molecular weights of assembled complexes.

Following affinity purification, intact-mass spectrometry of the α/β complexes was used to quantify the relative abundances of distinct pairing states. The assays were performed using two representative orthogonal designs, combiC26 and desV30combiC42, comprising engineered mutations in the constant domains alone (desC21, desC27, and desC100 for combiC26) or in combination with variable-domain mutations (desC43, desC125, and desV30 for desV30combiC42). For both designs, intact-mass spectra under competitive conditions were dominated by the correctly paired complex, with no obvious off-target peaks exceeding ∼5% relative abundance above features observed in the non-competitive controls, indicating an estimated ∼95% correct pairing.

To evaluate whether orthogonalization of the TCR α/β interface affected antigen recognition, we assessed peptide MHC (pMHC) binding by surface plasmon resonance (SPR) analysis (**Fig. 3h**). SPR single-cycle kinetic experiments were performed using immobilized pMHC ligands and soluble TCR analytes. The orthogonal TCRs exhibited binding affinities and kinetic profiles to their cognate pMHCs that were comparable to those of the corresponding parental TCRs, indicating that the engineered interface mutations did not measurably perturb antigen recognition.

Next, to validate orthogonality in a therapeutically relevant format, we adopted an antibody-like architecture in which two orthogonal TCRs replace the Fab domains and are fused to a previously-described heterodimeric Fc (7.8.60 Fc)^24,30^. Using this framework, we explored bispecific TCR-IgG (bsTCR-IgG) architectures incorporating two orthogonal TCRs, identifying formats that enable efficient assembly and correct pairing (**Supplementary Fig. S8**). Based on this analysis, we constructed a representative bsTCR-IgG comprising two orthogonal TCRs derived from parental MART-1 specific (KD: ∼100 pM) and NY-ESO-1 specific (KD: ∼1 nM) TCRs. We obtained a high yield of fully-assembled bsTCR-IgG by co-expressing the following domains in Expi293F cells: 1) TCR1α-Fc (7.8.60-A), 2) TCR2α-Fc (7.8.60-B), 3) TCR1β, and 4) TCR2β (**Fig. 3i**). Mass spectrometry showed ∼95% of the α and β chains were correctly paired (**Fig. 3j**). An SPR binding assay further showed that the bsTCR-IgG bound their cognate pMHCs with the expected affinities, indicating that the mutations do not affect the binding to pMHC (**Fig. 3k**).

### Crystal structure of an orthogonal constant-domain interface

To obtain atomic-level validation of the orthogonal TCR interface, we determined the X-ray crystal structure of the combiC26 constant-domain heterodimer, which combines three independently engineered design modules (desC21, desC27, and desC100). Crystal structure data collection and refinement statistics are provided in **Supplementary Table S10**. The structure was refined to 2.05 Å resolution and preserves the overall Cα/Cβ constant-domain architecture, demonstrating that orthogonality is achieved through local remodeling of the interchain interface rather than global distortion of the fold. Mapping the engineered residues onto the constant-domain scaffold revealed three spatially separated patches at the Cα/Cβ interface, consistent with a modular design strategy (**Fig. 4a**).

**Figure 4.**
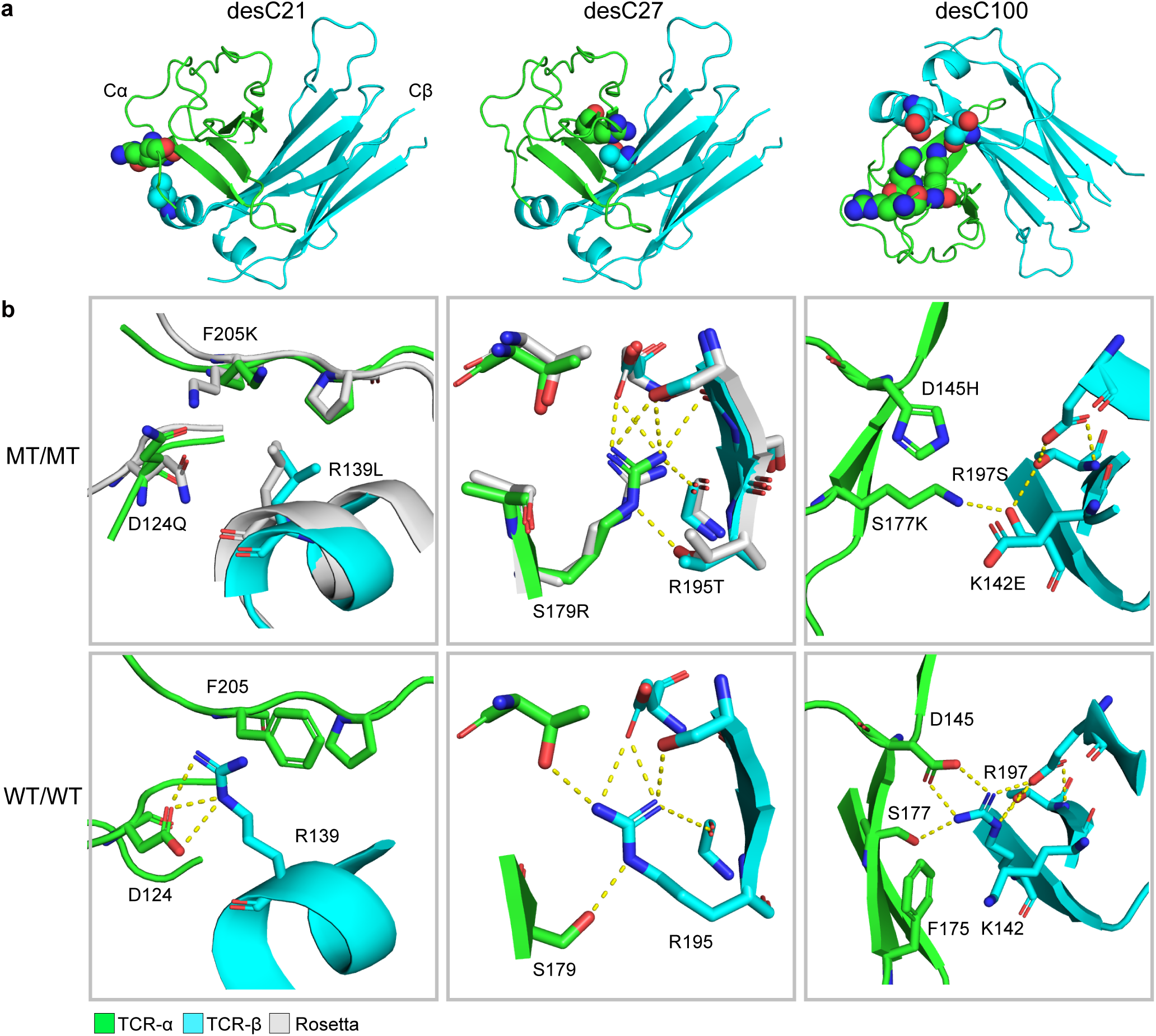
Crystal structure of designed orthogonal TCR α/β interfaces. **a**, X-ray crystal structure of the engineered constant-domain heterodimer combiC26, with the three design modules desC21, desC27, and desC100 mapped onto the TCR Cα/Cβ interface. Engineered residues are shown as spheres. **b**, Close-up structural comparisons of the Rosetta design models (gray) and the experimentally determined crystal structure (green: Cα, cyan: Cβ) for each design patch, shown alongside the corresponding wild-type interface. The desC100 patch was generated by a rational design and is shown without a corresponding Rosetta model.

We then analyzed each design module by comparing the designed MT/MT interface to the parental WT/WT interface and by overlaying the Rosetta design models (gray) with the crystallographic model (**Fig. 4b**). In the desC21 design (Cα-D124Q/F205K, Cβ-R139L), the WT interface is stabilized by a Cβ-R139-centered interaction network, including a salt bridge to Cα-D124 and a cation-π contact with Cα-F205. In combiC26, these polar/aromatic interactions are eliminated and replaced by a repacked hydrophobic interface in which mutant Cβ-L139 engages packing interactions that involve wild-type Cα-P207, with the observed side-chain conformations in close agreement with the Rosetta model. In the desC27 knob-into-hole design (Cα-S179R, Cβ-R195T), the crystallographic structure closely matches the Rosetta model, with Cα-R179 inserting into the pocket created by Cβ-T195. Notably, Cβ-T195 participates in an extended hydrogen-bond network involving Cα-R179 and neighboring residues, providing a structural basis for the high stability of the correctly paired complex. Finally, in the desC100 charge-swap design (Cα-D145H/F175R/S177K, Cβ-K142E/R197S), which was generated through a rational approach rather than Rosetta-based modeling, a new interchain hydrogen bond between mutant Cα-K177 and Cβ-S197 is observed, whereas several polar interactions present in WT/WT are not preserved. Two adjacent loops display weak or absent electron density, indicating local flexibility in this region. Together, these structures provide direct validation that orthogonal interfaces can be achieved through distinct, modular mechanisms, and that Rosetta models accurately capture the intended geometry and stabilizing chemical interactions for the computationally designed interfaces.

### Development of Trispecific T-cell engagers (TriTE) comprising two orthogonal TCRs

Single TCRs have been employed in T-cell redirection strategies by fusing them to an antibody single chain Fv (scFv) that binds T-cell activating subunits such as CD3ε^40,41^. This is exemplified by immune mobilizing monoclonal T-cell receptors against cancer (ImmTACs)^42,43^, such as gp100-targeting tebentafusp, which has received FDA approval for unresectable or metastatic uveal melanoma^44^. Nonetheless, the intratumoral heterogeneity of solid tumors poses an inherent challenge for single-antigen-targeting T-cell redirection therapies, which are often limited by the heterogeneous antigen expression profile and the possibility of immune escape via antigen-loss variants. Trispecific T-cell engagers (TriTEs) can address this limitation by targeting two distinct tumor antigens, enabling elimination of heterogeneous tumor cell populations expressing either or both antigens, while potentially providing enhanced potency through bivalent engagement.

To explore the TriTE format, prior to incorporation of anti-CD3 domains, we first validated tandem dual-TCR architectures comprising two orthogonal TCRs and identified the α1-α2 + β1 + β2 configuration as the most favorable in terms of assembly and expression (**Supplementary Fig. S9**). Next, to develop functional TriTE formats, we tested 24 architectural variants, reflecting three permutations of domain order, two anti-CD3 domain formats (scFv or Fab), two permutations of heavy chain (HC) and light chain (LC) order, and two anti-CD3 domains: SP34 ^30,45^ or humanized UCHT-1 anti-CD3^42,46^. Based on SDS-PAGE, mass photometry, and cytotoxicity profiling (**Supplementary Fig. S10–11**), we identified an optimal architecture consisting of one principal subunit (TCR1α-SP34LC-TCR2α) and three auxiliary subunits (TCR1β, SP34HC, TCR2β) (**Fig. 5a**). Using this format, we constructed TriTEs incorporating TCRs with picomolar affinities for the cancer-testis antigens NY-ESO-1 (KD = 40 pM), MAGE-A4 (KD = 74 pM), and PRAME (KD = 24 pM), based on previously described high-affinity TCRs developed for the ImmTAC platform^47–49^. Mass photometry demonstrated complexes with molecular weight ∼160 kDa as expected (**Fig. 5b**). Using the desV30combiC46 design across multiple TCR pairs (NY-ESO-1/MAGE-A4, NY-ESO-1/PRAME, and MAGE-A4/PRAME), nanoDSF and SDS-PAGE analyses showed higher thermal stability and expression for the on-target α/β pairs (MT/MT and WT/WT) compared with the off-target pairs (MT/WT and WT/MT), demonstrating that the orthogonal interface mutations are effective for these TCR pairs (**Fig. 5c**). Framework annotations for the TCRs evaluated here are included in **Supplementary Table S8**.

**Figure 5.**
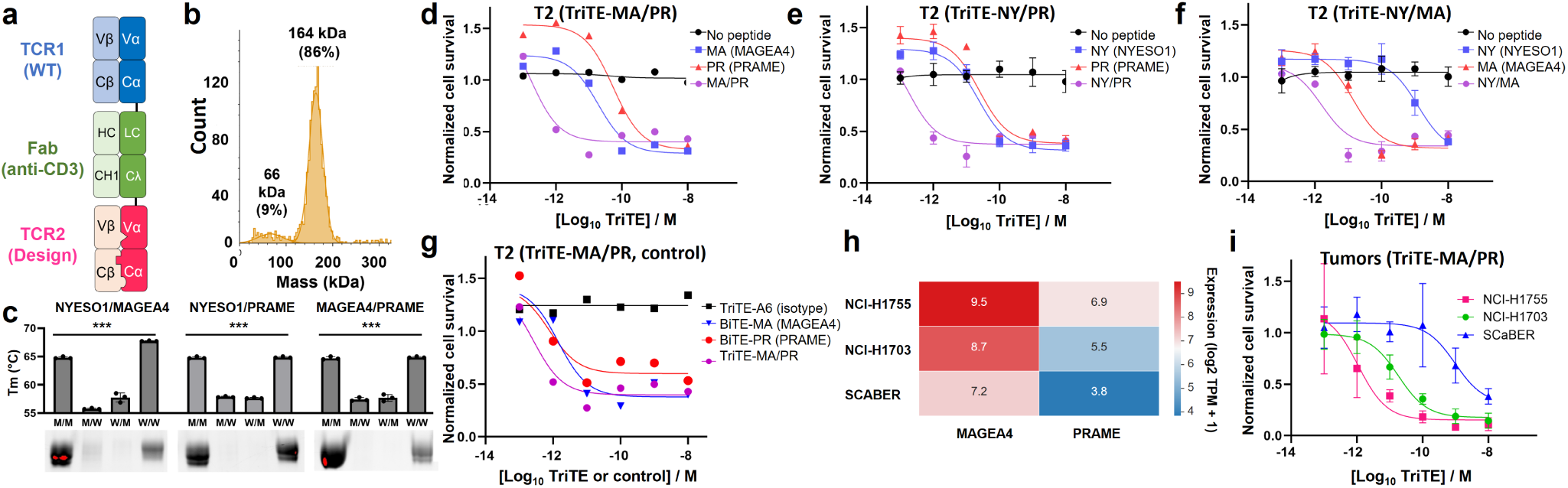
Trispecific T-cell engagers (TriTEs) comprising two orthogonal TCRs. **a**, Schematic representation of a trispecific T-cell engager (TriTE) comprising a TCR with a wild-type α/β interface (TCR1), a TCR with a designed orthogonal α/β interface (TCR2), and an anti-CD3ε Fab. **b**, Mass photometry analysis of purified TriTE showing a predominant species at the expected molecular mass and minor lower-molecular-weight species. **c**, Thermal stability assessed by nano-differential scanning fluorimetry (nanoDSF) and SDS-PAGE analysis of orthogonal TCR pairs (NY-ESO-1/MAGE-A4, NY-ESO-1/PRAME, and MAGE-A4/PRAME) using the desV30combiC46 design, across M/M, M/W, W/M, and W/W pairing states (W: WT, M: MT). Bars show mean ± s.d. (n = 3 replicates); statistical analysis by one-way ANOVA; ***P < 0.001. **d–f**, Redirected T-cell cytotoxicity mediated by TriTE-MA/PR **(d)**, TriTE-NY/PR **(e)**, and TriTE-NY/MA **(f)** against TAP-deficient T2 cells, either unpulsed or pulsed with a single peptide or both peptides. Data are shown as mean ± s.d. (n = 3 replicates). **g**, Comparison of TriTE-MA/PR with a TriTE isotype control (A6-TCR) and single-TCR BiTEs (BiTE-MA and BiTE-PR) in T2 cell killing assays. Data are shown as mean ± s.d. (n = 3 replicates). **h**, Expression of MAGE-A4 and PRAME across human tumor cell lines derived from Cancer Cell Line Encyclopedia (CCLE) RNA-seq dataset, shown as log2-transformed transcripts per million (TPM) values with a pseudocount of 1. Selected HLA-A*02:01-positive tumor cell lines used for functional assays are highlighted. **i**, TriTE-MA/PR-mediated cytotoxicity against HLA-A*02:01 positive tumor cell lines endogenously expressing MAGE-A4 and PRAME (NCI-H1755, NCI-H1703, and ScaBER). Data are shown as mean ± s.d. (n = 3 replicates).

To evaluate antigen-specific cytotoxicity, we used TAP-deficient T2 cells^50^, which present exogenously loaded but not endogenous peptides, providing a controlled system to assess peptide-dependent activity. Our lead construct, TriTE-MA/PR, targeting the MAGE-A4 (230–239; GVYDGREHTV) and PRAME (425–433; SLLQHLIGL) peptides presented by HLA-A*02:01, demonstrated highly potent cytotoxic activity. TriTE-MA/PR exhibited sub-picomolar potency (EC50: 380 fM) when T2 cells were pulsed with both MAGE-A4 and PRAME peptides and retained activity when pulsed with either MAGE-A4 (EC50: 48 pM) or PRAME (EC50: 20 pM) alone. In contrast, no detectable cytotoxicity was observed on unpulsed T2 cells even at the highest TriTE concentration tested (10 nM), indicating a large activity window between peptide-positive and peptide-negative conditions in this assay. Together, these results demonstrate enhanced potency under dual-peptide conditions, highlighting the functional advantage conferred by bivalent engagement (**Fig. 5d**). Similarly, TriTE-NY/PR and TriTE-NY/MA showed enhanced dual-antigen-dependent cytotoxicity (**Fig. 5e,f**). Further comparisons under dual-peptide-pulsed conditions showed that TriTE-MA/PR exhibited greater cytotoxic potency than the corresponding single-TCR BiTEs (BiTE-MA and BiTE-PR), whereas a TriTE isotype control targeting an irrelevant antigen (A6-TCR^51^, specific for the HTLV-1 Tax protein) showed no measurable activity (**Fig. 5g**).

We further assessed the efficacy of TriTE-MA/PR against HLA-A*02:01-positive cancer cell lines endogenously expressing MAGE-A4 and PRAME, including squamous cell lung cancer (NCI-H1703), lung adenocarcinoma (NCI-H1755), and squamous cell carcinoma of the bladder (ScaBER). Expression of MAGE-A4 and PRAME across human tumor cell lines, derived from publicly available RNA-seq datasets^52^, is summarized in a heatmap (**Fig. 5h**). TriTE-MA/PR mediated potent cytotoxicity across all lines tested. Notably, against the lung adenocarcinoma cell line NCI-H1755, TriTE-MA/PR achieved an EC50 of 1.1 pM, confirming its ability to mediate robust cytolysis based on endogenous antigen presentation (**Fig. 5i**). These results demonstrate that orthogonal TCR designs enable functional assembly of TriTEs capable of potent cytotoxicity against cancer cells expressing endogenous tumor antigens, providing a platform for dual-antigen targeting in heterogeneous tumors.

## Discussion

We combined systematic computational design with quantitative biophysical and biochemical characterization to engineer orthogonal TCR interfaces, enabling assembly of dual-TCR therapeutics with high (>95%) pairing accuracy. By focusing designs on constant domains and the highly conserved framework regions within variable domains, we generated a broadly applicable design set across diverse TCR sequences without compromising antigen recognition. Using these orthogonal interfaces, we developed trispecific T-cell engagers (TriTEs) composed of two TCRs and an anti-CD3ε domain. TriTEs targeting two cancer-testis antigens exhibited enhanced cytotoxic potency under dual-antigen engagement with a widened activity window in peptide-pulsed assays, while retaining activity against single-antigen-positive cells, thereby addressing tumor heterogeneity.

Achieving this level of orthogonality required an integrated pipeline of computational design and experimental validation. Multistate design (MSD) provided deep optimization within selected interface clusters, while site-saturation mutagenesis and second-site suppressor (SSM/SSS) searches enabled broader exploration across all interface positions, together providing comprehensive coverage of the interface design space. By characterizing designs with mutations in separate interface regions, our results indicate that robust orthogonality requires coordinated engineering across both variable and constant domain interfaces. Quantitative biophysical characterization combining nanoDSF, SDS-PAGE, mass spectrometry, and X-ray crystallography was essential to resolve subtle energetic and structural differences and quantify pairing fidelity among design modules. This modular engineering strategy provides a generalizable framework for constructing fully orthogonal heterodimeric proteins. These design principles are expected to extend beyond physics-based scoring to deep learning-guided approaches such as AlphaFold^53^ confidence metrics and ProteinMPNN^54^ sequence probabilities, broadening the scope of orthogonal interface engineering across therapeutic applications.

Comparison with prior approaches revealed important mechanistic differences in preventing mispairing. Despite their widespread use to mitigate mispairing, murinized constant domains did not effectively suppress mispaired assemblies, with huα/muβ mispaired complexes showing higher expression yields and comparable thermal stability to correctly paired complexes (Fig. 3c). Given that reported minimally murinized variants, which achieve T-cell surface expression comparable to full murinization, place mutations outside the canonical α/β interface, instead localizing primarily to hinge-adjacent and transmembrane regions of the constant domains, murinization likely operates primarily through expression enhancement rather than interface-level orthogonality. In contrast, our orthogonal designs yielded diverse interface mechanisms including knob-into-hole packing, charge swaps, polar-to-hydrophobic substitutions, and cation–π contacts. These complementary mutations, where α-chain modifications are specifically matched by compensating changes in the β chain, directly destabilize mispaired assemblies and reduce their expression, as confirmed by quantitative biophysical and structural validation.

Among our designs, desC27 (Cα-S179R/Cβ-R195T) and desC43 (Cα-S179R/Cβ-R195S), both independently generated from MSD cluster 4, adopted a knob-into-hole configuration^10^ similar to a TCR redesign reported by Voss et al. However, Voss et al. used a Cβ-R195G mutation instead of R195T/S. To test the relative benefits of R195T/S versus R195G, we created desC41 that includes R195G instead of the serine or threonine at position 195. Compared with desC43, which showed Tm values of 75.0 °C (MT/MT), 66.8 °C (MT/WT), and 69.9 °C (WT/MT), desC41 exhibited substantially lower stability and orthogonality with Tm values of 64.7 °C (MT/MT), 67.5 °C (MT/WT), and 64.1 °C (WT/MT), corresponding to an approximately 10 °C loss in thermal stability of the correctly paired complex. Crystal structure analysis of desC27 revealed that Thr195 (Cβ) forms a stabilizing hydrogen-bond network with Arg179 (Cα) and neighboring residues, providing a structural basis for its enhanced stability. This structural validation demonstrates that achieving orthogonal interfaces requires not only geometric complementarity but also preservation of stabilizing chemical interactions.

These orthogonal interfaces enabled dual-TCR therapeutics with substantial therapeutic advantages. Simultaneous engagement of two target antigens on the same cell induces avidity effects that markedly enhance potency and specificity. TriTE-MA/PR targeting MAGE-A4 and PRAME-derived pMHC achieved an EC50 of 380 fM when both antigens were present on peptide-pulsed T2 cells, compared to 48 pM and 20 pM for individual antigens, demonstrating enhanced potency under dual-antigen engagement. Robust picomolar potency was also observed against cancer cell lines with endogenous antigen expression, with an EC50 of 1.1 pM against NCI-H1755. In addition, TriTE-MA/PR retained potency comparable to single-TCR BiTEs against cells presenting only a single antigen, providing a strategy to mitigate antigen escape.

These orthogonal design principles pave the way for dual-TCR-based multispecific immunotherapies with enhanced tumor selectivity and broader antigen coverage.

## Methods

### Computational multistate design (MSD) of orthogonal TCR interfaces

Computational multistate design (MSD) was applied to two human T-cell receptor (TCR) interfaces: the constant-domain (PDB: 6U07) and the framework region of variable domain (PDB: 2F53). MSD simulations were performed using a previously described workflow (https://github.com/aleaverfay/generic_msd). To ensure broad applicability, mutations were confined to constant domains and conserved framework residues while CDR loops remained fixed. Designable positions were selected by structural inspection of contact distance (<6 Å), buried surface area, and inter-chain hydrogen-bond or salt-bridge networks, excluding residues critical for intra-domain packing or disulfide geometry. Residues were grouped into clusters of 2–12 residues for tractable optimization.

Each cluster was optimized across multiple binding states following established MSD protocols. The positive state corresponded to the mutant-mutant (MT/MT) heterodimer, and the negative states to mixed dimers (MT/WT and WT/MT). Sequence optimization used Rosetta ref2015 all-atom scoring with side-chain packing and minimization. Interface energies (dG_separated), buried surface area (dSASA_int), and unsatisfied H-bonds were evaluated using InterfaceAnalyzer. Sequences were ranked by a custom multi-objective fitness function that favored MT/MT stability while penalizing mixed-state stabilization and excessive mutation load (details in Supplementary Methods). Top-ranked designs were relaxed with FastRelax under backbone restraints, and selected clusters were reanalyzed with expanded state ensembles to improve backbone accommodation. Representative Rosetta inputs, resfiles, and parameters will be provided in Supplementary Methods.

### In silico site-saturation mutagenesis and second-site suppressor screening

To explore the broader sequence landscape of the TCR variable-domain interface, we performed a two-stage computational workflow combining site-saturation mutagenesis (SSM) and second-site suppressor (SSS) screening, adapted from the workflow by Thieker et al^32^. In the SSM step, all interfacial residues (within 8 Å of the partner chain) on Vα or Vβ were individually mutated to all canonical amino acids except cysteine. For each variant, side-chain and local backbone conformations were sampled within a 12 Å sphere using Rosetta Cartesian FastRelax (ref2015), while outer regions were held fixed under coordinate constraints. Changes in total energy (ΔE) and interface binding energy (dG_separated, InterfaceAnalyzer) identified stabilizing mutations as seeds for combinatorial design. In the SSS screening step, each seed was subjected to combinatorial optimization using Rosetta’s InterfaceDesign2019 FastDesign protocol. Residues within 15 Å of the seed were allowed conformational freedom, with an inner design shell defined by atomic contact (CloseContact selector, 4.5–8 Å) and interface geometry (InterfaceByVector). Asymmetric native-retention biases (ResidueTypeConstraintGenerator; stronger near seed, weaker on partner chain) promoted compensatory mutations while limiting excessive substitutions. Designs were ranked by dG_separated, packing quality, and mutation count.

### TCR sequence annotation and analysis

Human TRAV and TRBV germline sequences were obtained from the ImMunoGeneTics (IMGT) database^37^. Conservation of TCR variable-domain framework residues was assessed by generating sequence logos for TRAV and TRBV gene sets using WebLogo^55^. IMGT unique numbering was assigned to representative TCR structures, and Colliers de Perles representations were generated using IMGT tools to define the topological context of conserved framework and CDR regions, including positions relevant to the TCR-V interface design strategy^56^. TCR variable domain sequences were annotated and numbered using the Ab-Numbering tool (NovoPro Labs; https://www.novoprolabs.com/tools/ab-numbering). ANARCI was used to identify TCR V-gene segments and assign IMGT numbering schemes for sequence annotation purposes^57^. Explicit residue numbering correspondence maps between IMGT, Kabat, and PDB schemes were generated for the TCR constant-domain structure (PDB: 6U07) and the TCR variable-domain structure (PDB: 2F53), and are provided in **Supplementary Tables S11–S14**.

### Plasmid construction

For the construction of soluble TCR variants, the constant regions were truncated at C213 of the TCR α constant domain (Cα; gene TRAC, UniProt P01848) and C247 in the human TCR β2 constant domain (Cβ; gene TRBC2, UniProt A0A5B9), thus retaining their native disulfide bond. For the stabilized TCR variants (stTCR), an additional interchain disulfide bond was introduced by incorporating T166C in Cα and S173C in Cβ mutations as previously reported^58^. To further enhance stability, additional mutations were introduced: T150I, A190T, and S139F (Cα), and E134K, H139R, D155P, and S170D (Cβ), as previously described^30^. The residue numbering follows the Kabat numbering scheme. For bispecific TCR-IgG constructs, a heterodimeric Fc scaffold (7.8.60 Fc) was used to enable orthogonal pairing of the two Fc chains. Individual TCRα chains were fused to the N terminus of the 7.8.60-A or 7.8.60-B Fc chains, while the corresponding TCRβ chains were expressed as separate polypeptides. For TriTE constructs, tandemly linked TCR ectodomains were fused to anti-CD3 modules. Anti-CD3 modules were derived from either the humanized UCHT-1 clone (GenBank accession numbers AAB24133.1 and AAB24132.1) or the SP34 clone, as indicated. DNA fragments encoding the design sequences were synthesized (Twist Bioscience) and cloned into the modified pαH mammalian expression vector, originally derived from the pHLSec vector^59^. A human serum albumin signal peptide for secreted expression was incorporated into all constructs. Soluble TCR and TriTE constructs contained an 8×His tag at either the N terminus or the C terminus for affinity purification.

### Protein expression and purification

Constructs were expressed in Expi293F cells (Thermo Fisher Scientific, A14527) cultured in Expi293 Expression Medium on an orbital shaker (125 rpm, 37 °C, 8% CO₂). Cells were seeded at 2.5 × 10⁶ cells/mL one day before transfection and adjusted to 3.0 × 10⁶ cells/mL on the day of transfection. Transfection was performed using either ExpiFectamine 293 Transfection Kit (Thermo Fisher Scientific, A14525) or PEI-MAX (Kyfora Bio, 24765-100) according to manufacturer’s protocols with appropriate enhancers added at ∼18 h post-transfection. Cultures were harvested ∼96 h post-transfection, and supernatants were collected by centrifugation (3428 × g, 4 °C, 15 min) and filtered through 0.22 µm PVDF membranes (MilliporeSigma, SLGVR33RS). Proteins were purified by affinity chromatography using penta-Ni resin (Marvelgent Biosciences, 11-0228-100) with sequential washes in Tris buffer (pH 8.1) or PBS (pH 7.4) containing increasing imidazole concentrations (typically 20 mM, 40 mM) and elution with 500 mM imidazole. Typical yields of stabilized TCR-C constructs were approximately 80–100 mg/L, and yields of TCR-VC constructs varied depending on the design and were generally 5–20 mg/L. bsTCR-IgG constructs were purified using Protein A affinity resin (GenScript, L00210). Eluted proteins were buffer-exchanged into PBS (pH 7.4), concentrated using Amicon Ultra Centrifugal Filters (10 kDa MWCO, MilliporeSigma, UFC801008). When necessary, proteins were further purified by size exclusion chromatography on Superdex 200 Increase 10/300 GL (Cytiva, 28990944) using an ÄKTA FPLC system at 4 °C. Fractions containing target proteins were pooled and concentrated.

### Assessment of protein stability by differential scanning fluorimetry

Thermal stability of purified TCR proteins was assessed by nano-differential scanning fluorimetry (nanoDSF) using a Prometheus NT.48 instrument (NanoTemper Technologies). Samples (10 µL) were prepared at 10 µM or the highest achievable concentration after purification, then heated from 20 to 95 °C at 1 °C/min while monitoring intrinsic tryptophan fluorescence at 330 nm and 350 nm. Melting temperatures (Tm) were calculated using PR.ThermControl software.

### Evaluation of TCR subunit pairing fidelity by ZipChip MS analysis

Samples were submitted in a 50 mM ammonium acetate buffer and were each diluted 1:1 with a background electrolyte (BGE) made of 49.5% acetonitrile, 49.5% water, 1% formic acid prior to CE-MS. Samples were injected onto an HR Chip (908Devices Inc.) using a ZipChip system (908Devices Inc.) interfaced to a Thermo QExactive HF Biopharma mass spectrometer. The MS data were acquired through Tune (v. 2.9) and settings included: scan range 500-2500 m/z, in-source CID 15eV, mass resolution 15,000, 3 microscans, AGC target 3e6, 10 ms maximum injection time. The CE settings included: BGE type = peptides, field strength 500 V/cm, injection volume 1 nL, pressure assist start time 0.5 min, analysis run time 5 min. Intact mass spectra were deconvoluted using the Byos protein characterization software (Protein Metrics Inc.).

### Affinity measurement with Surface Plasmon Resonance (SPR)

Experiments were conducted using a Biacore 8K instrument (Cytiva) at 25 °C in PBS containing 0.05% Tween 20, pH 7.4 (PBS-T). For binding assays, biotinylated human pMHC monomer was diluted to ∼1.5 µg/mL in PBS-T and immobilized on a NeutrAvidin chip (Cytiva, 29407997) to a level of ∼200 response units (RUs). Following immobilization, affinity measurements were performed using five successive 1:3 serial dilutions of soluble TCRs (at concentrations indicated in figures) using a single-cycle kinetics method with 120 s of association and 1,200 s for the final dissociation time at 25 °C and a flow rate of 30 µL/min. All data were analyzed with the Biacore Insight Evaluation software version 3.0.12 with the general single-cycle kinetics method and a 1:1 binding kinetics fit model.

### **X-** ray crystallography

Protein was concentrated to 55 mg/mL and then tested for crystallization against common commercially available crystal screens using a Mosquito robot (SPT Labtech) with drops composed of 150 nL protein and 150 nL reservoir solution set over 30 µL reservoir volumes. Crystals were observed with a solution composed of 100 mM sodium cacodylate pH 6.5 and 1.26 M ammonium sulfate. The crystals were briefly soaked in reservoir supplemented with 15% ethylene glycol and then cryocooled in liquid nitrogen. Diffraction data were collected on a Bruker D8 Venture equipped with an IµS copper sealed microfocus source, a Photon III C14 pixel array detector, and an Oxford Cryostream 800. Data were collected and reduced in Bruker Proteum^60,61^. The structure was phased by molecular replacement using Phaser^62^ with PDB 6U07 as the search model^30^. Real-space rebuilding was done in Coot^63^, and reciprocal space refinements and validations were done in PHENIX^64^. Coordinates and structure factors will be deposited in the Protein Data Bank (PDB).

### T-cell-redirected tumor cell cytotoxicity assay

T-cell-redirected tumor cell killing was assessed using primary human pan-CD3+ T cells (StemExpress, PB03020C) expanded for 4–14 days in complete RPMI 1640 medium supplemented with anti-CD28 (BD Biosciences, 555725, 2.5 µg/mL) and IL-2 (R&D Systems, 202-IL/CF, 2 ng/mL) in flasks precoated with anti-CD3 antibody (BD Biosciences, 555329, 5 µg/mL). Target tumor cells were seeded in 96-well plates and incubated for 4–6 h or overnight at 37 °C with 5% CO₂. For peptide-pulsed conditions, cells were washed, incubated with peptide at 1 µM for 3 h, then washed twice. Expanded T cells were added at indicated effector-to-target ratios together with serially diluted BiTE or TriTE molecules and incubated for 24–48 h. Plates were washed twice to remove non-adherent T cells and remaining viable tumor cells were quantified using CellTiter-Glo 2.0 (Promega, G9242) according to manufacturer’s instructions.

Luminescence was measured using a CLARIOstar Plus microplate reader (BMG Labtech). Cytotoxicity assays were conducted in technical triplicates. EC50 values were calculated by nonlinear regression using a four-parameter logistic equation in GraphPad Prism 10.

### Statistical analysis

Data are presented as mean ± standard deviation (SD) or standard error of the mean (SEM) as indicated. Statistical comparisons were performed using unpaired two-tailed Welch’s t-test or one-way ANOVA with post-hoc Tukey’s test for multiple comparisons. P-values <0.05 were considered statistically significant.

## Data availability

The data supporting the findings of this study are available within the paper. Orthogonal TCR interface designs and mutations are detailed in **Supplementary Tables S2–4** and have been described in patent application WO2024036166A1, published February 15, 2024^65^. Coordinates and structure factors for designed orthogonal T-cell receptors will be deposited in the Protein Data Bank upon publication. Plasmids encoding the major constructs will be deposited with Addgene and made available to academic researchers under a standard material transfer agreement.

## Code availability

Representative Rosetta scripts used for multistate design (MSD) and second-site suppressor (SSS) screening are available from the corresponding author upon reasonable request. Additional analysis workflows used for data analysis are available upon reasonable request.

## Acknowledgments

The authors are grateful to members of the Kuhlman Laboratory for their valuable input and assistance. This crystallography was in part supported by a grant from UNC Lineberger Comprehensive Cancer Center (P30 CA016086). The mass spectrometry is based in part upon work conducted using the UNC Metabolomics and Proteomics Core Facility, which is supported in part by NCI Center Core Support Grant (2P30CA016086-45) from the UNC Lineberger Comprehensive Cancer Center. The authors also thank the UNC Macromolecular Crystallography core facility, UNC Metabolomics and Proteomics Core Facility, UNC Research Computing, UNC Macromolecular Interactions Facility, UNC High-Throughput Peptide Synthesis Core Facility, and UNC Flow Cytometry Core Facility for technical assistance and access to instrumentation. The authors thank Duygu Koldere Vilain (Life Science Editors) and BioRender.com for assistance with figure illustrations. This work was supported by NIH grant R35GM131923 (B.K.); the Takeda Science Foundation Fellowship, the Murata Overseas Scholarship Foundation Fellowship, and the JSPS Overseas Research Fellowship from the Japan Society for the Promotion of Science (T.K.); NIH training grant T32GM086330, the Fred J. Ansfield, MD, Endowed Young Investigator Award from Conquer Cancer, the ASCO Foundation, and the American Cancer Society Postdoctoral Fellowship (PF-25-1412409-01-PFCDET) (S.Y.); and the UNC School of Medicine Translational Team Science Award (TTSA033P1) (B.K., W.Y.K., and S.Y.).

## Author contributions

T.K. and B.K. conceived and designed the study; T.K. performed computational design and experimental validation of orthogonal TCR interfaces and conducted data analysis under the supervision of A.L. and B.K.; T.K. designed and characterized bispecific TCR-IgG constructs; T.K. led the design and characterization of TCR-TriTE molecules with assistance from S.Y., under the supervision of B.K. and W.K.; N.N. supervised and performed X-ray crystallography with assistance from T.K.; B.K. supervised the overall project; T.K. wrote the original draft of the manuscript under the supervision of B.K., and all authors contributed to review and editing.

---

**Supplementary Figure S1.**
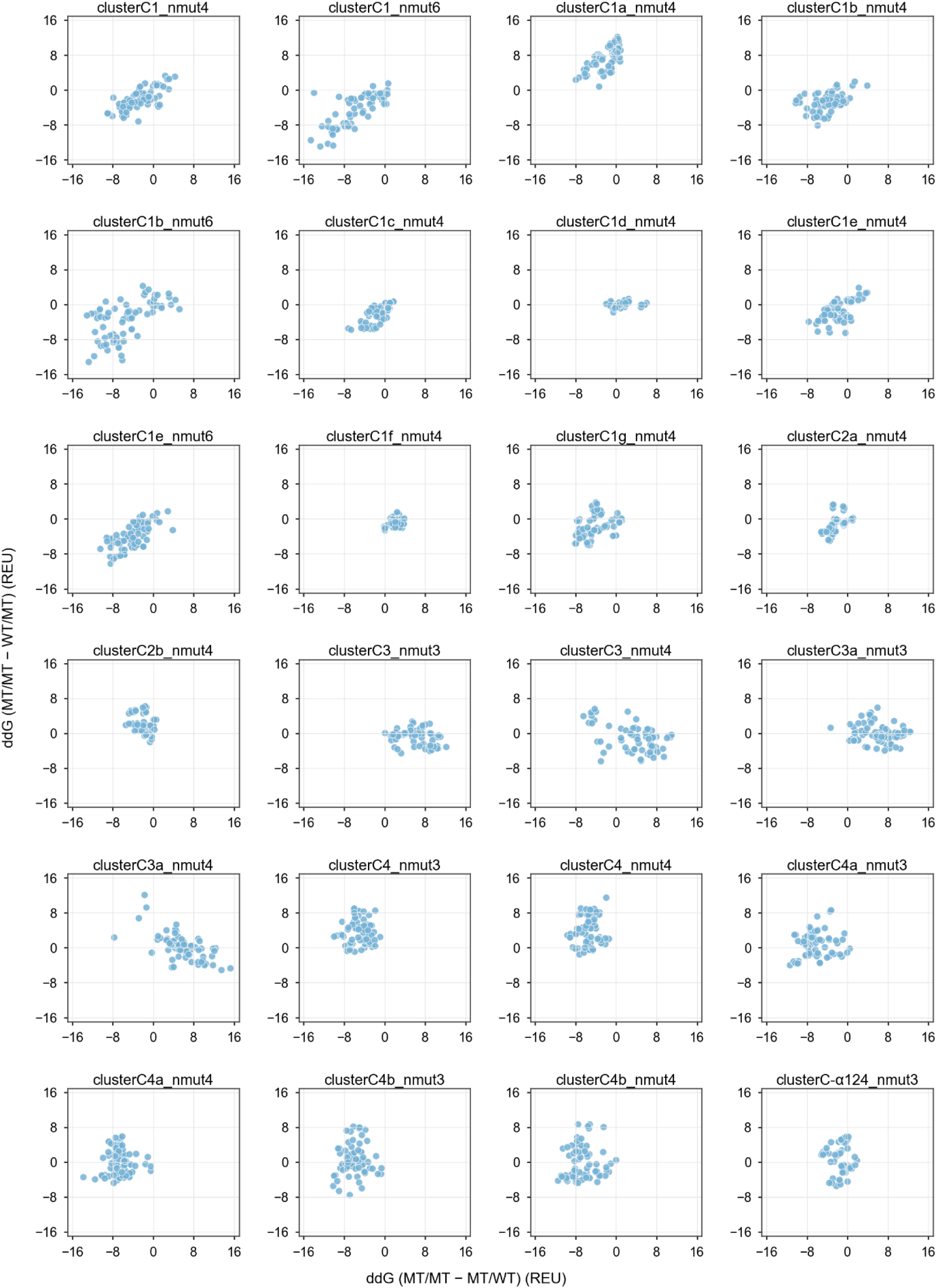
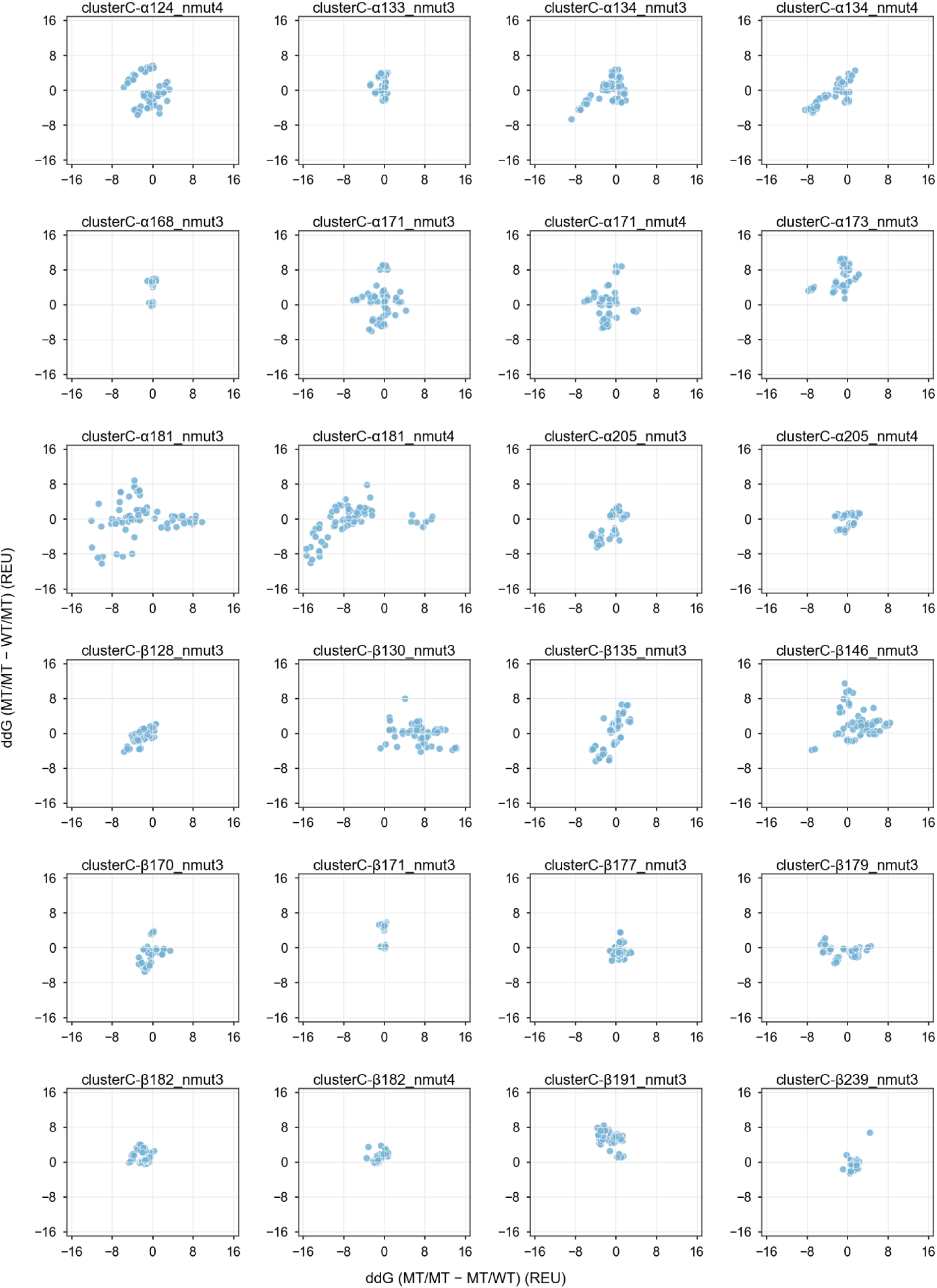
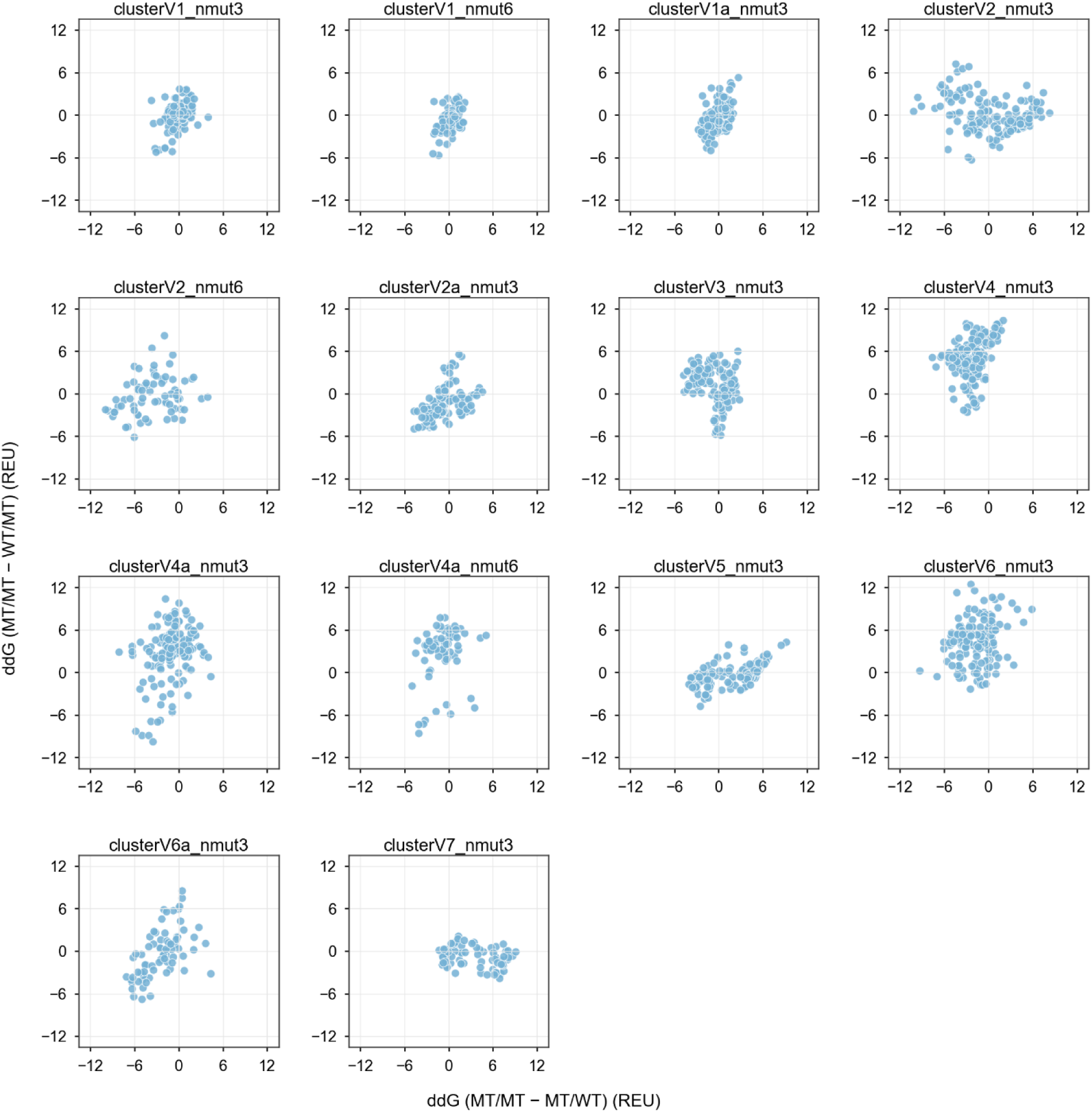
MSD interface energy differences across all residue clusters. Multistate design (MSD) interface energy scores (ΔΔG) are shown for all residue clusters at the TCR constant-domain interface and the conserved variable-framework interface. Scores show interface energy differences of off-target state (MT/WT and WT/MT) relative to the on-target MT/MT state for each cluster. MSD run include an nmut parameter that permits a number of mutations without penalty, with additional mutations incurring a progressive energetic penalty.

**Supplementary Figure S2.**
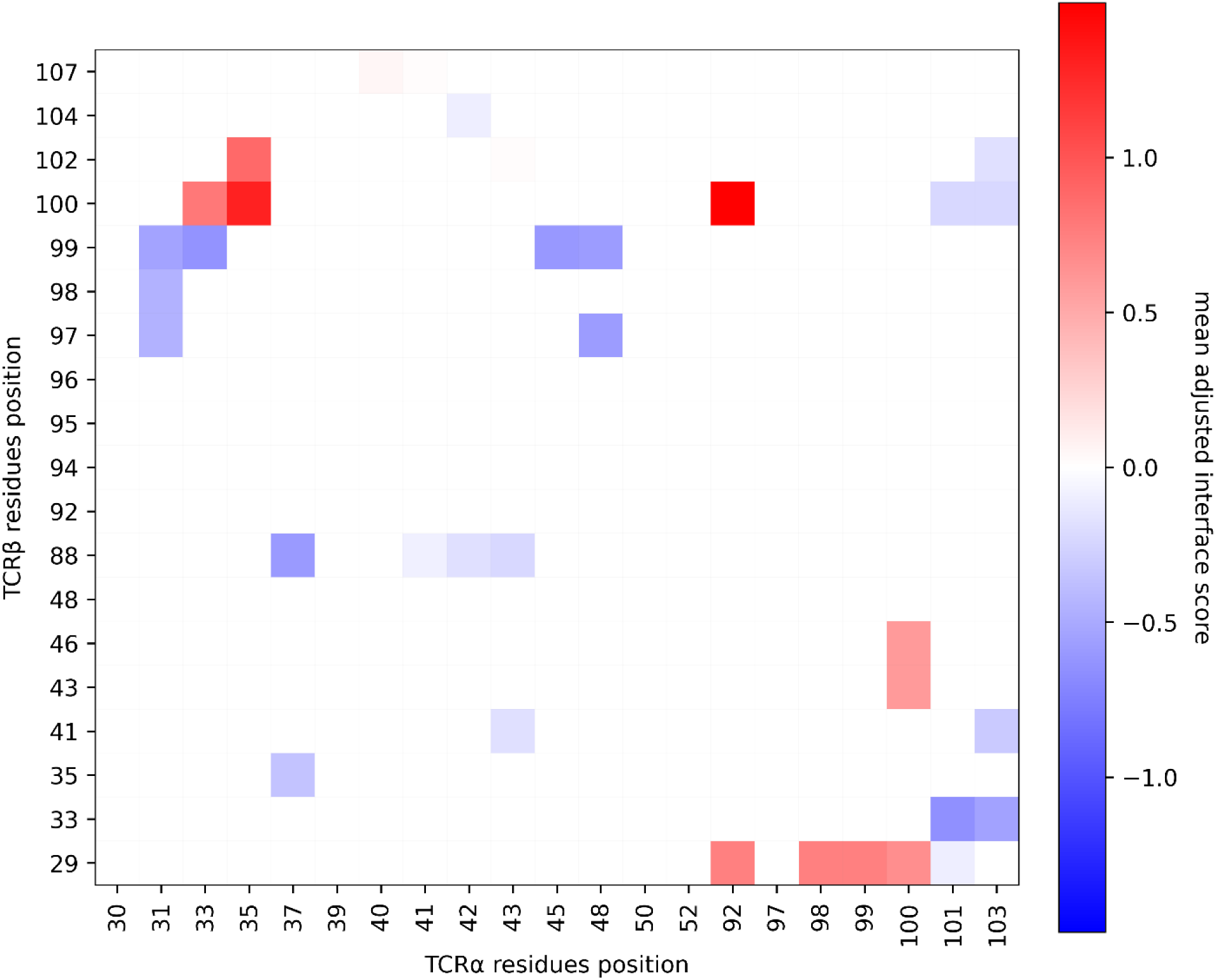
Mean adjusted MSD interface energy differences on the TCR-Vα/Vβ contact map. Residue-residue map of mean adjusted interface energy differences between on-target and off-target pairing states projected onto inter-chain contacting TCR-V α/β residue pairs, with colder colors indicating residue pairs for which off-target pairings are energetically less favorable relative to the on-target state. MSD scores, residue numbering, and inter-chain contacts are defined based on the TCR Vα/Vβ domains in the structure PDB: 2F53.

**Supplementary Figure S3.**
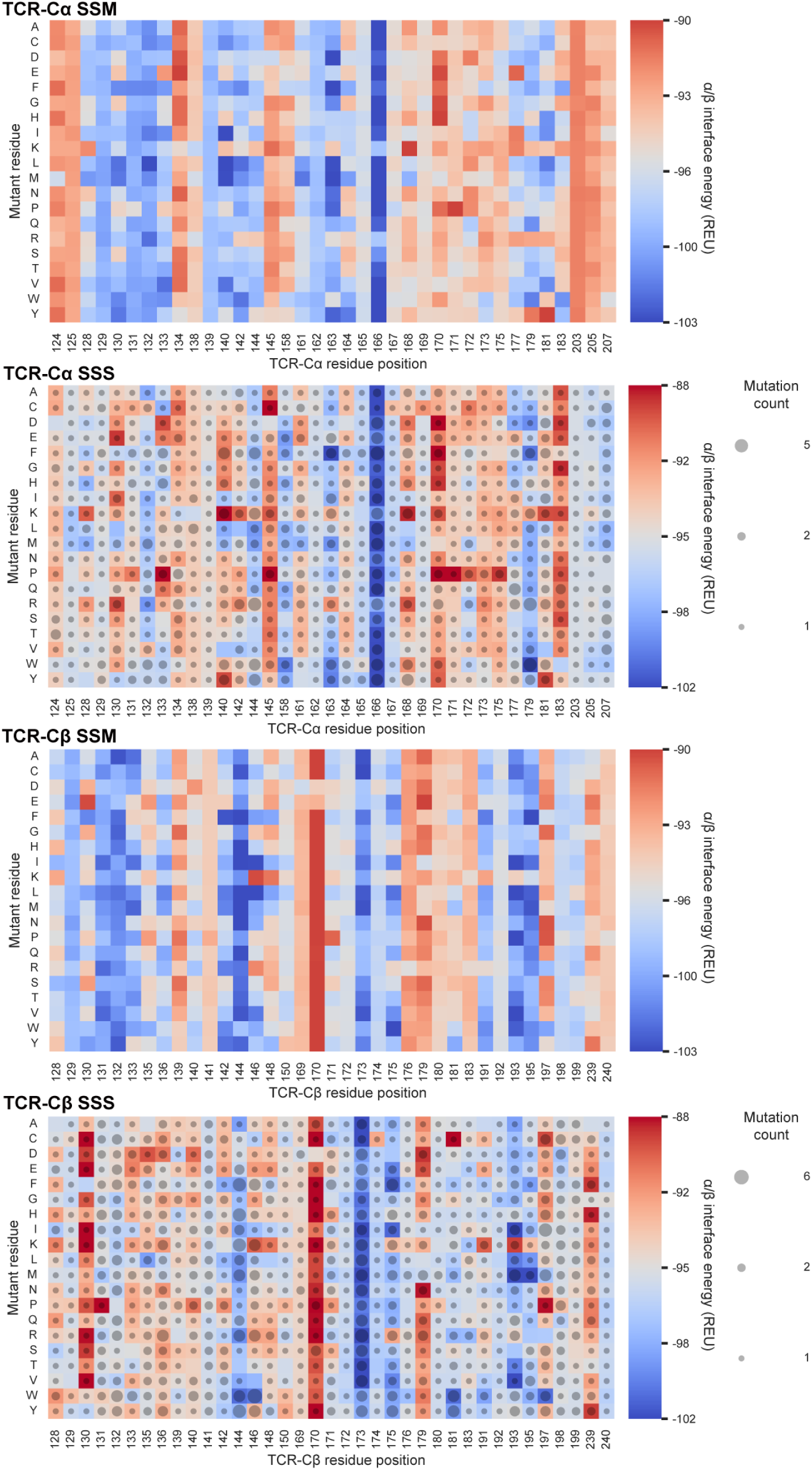
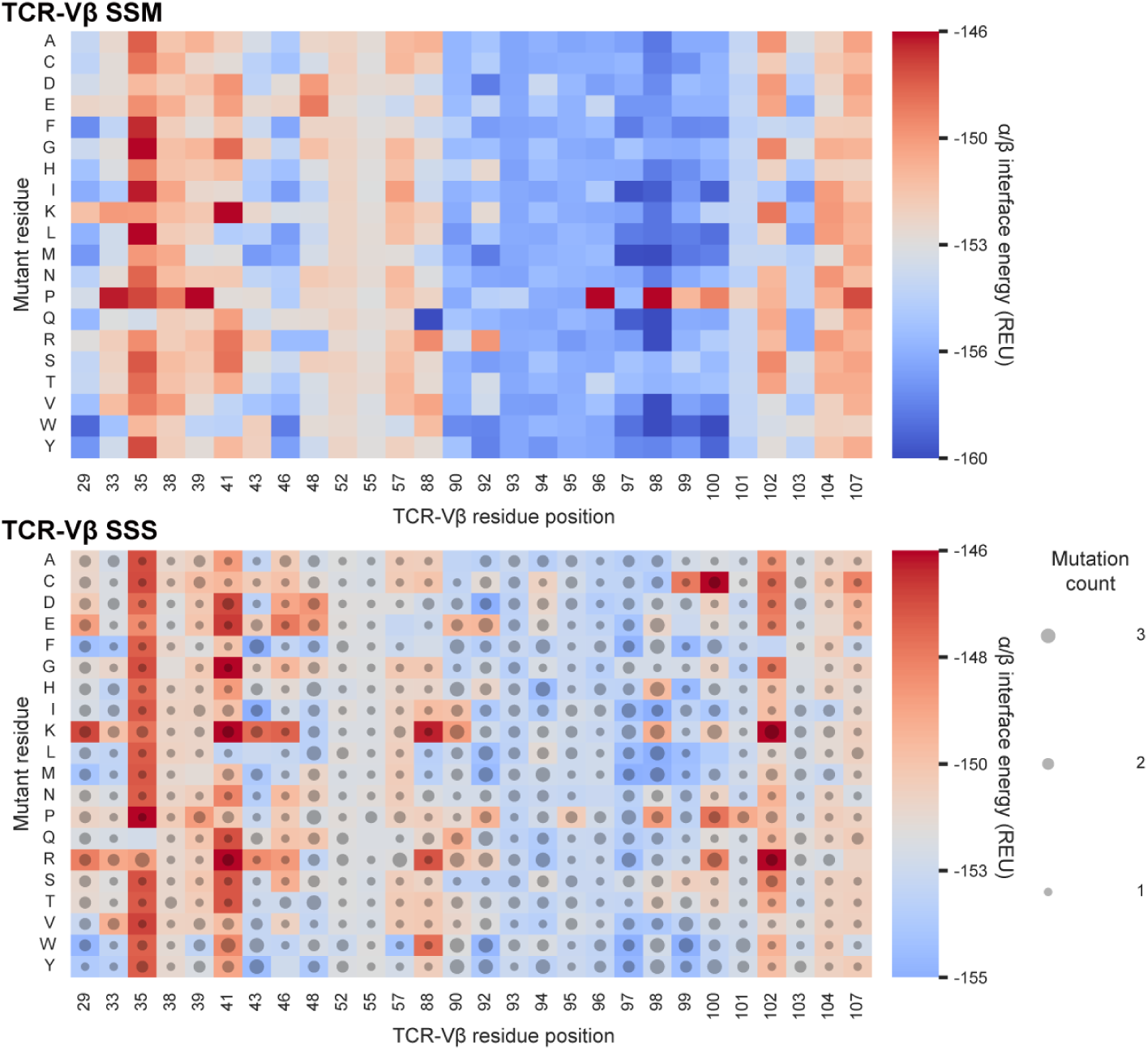
In silico SSM and SSS at TCR α/β interfaces. Heatmaps show in silico site-saturation mutagenesis (SSM) and second-site suppressor screening (SSS) at the TCR α/β interface. Panels labeled TCR-Cα SSM/SSS and TCR-Cβ SSM/SSS show constant-domain interface scans, and panels labeled TCR-Vβ SSM/SSS show variable-domain framework interface scans. In both SSM and SSS panels, colors indicate α/β interface energy, corresponding to the Rosetta dG_separated value (REU: Rosetta energy units), reported as the mean score for designs represented in each cell, with dots size reflecting the median number of mutations in the redesign. SSM/SSS results for TCR-Vα are shown in Fig. 2f,g.

**Supplementary Figure S4.**
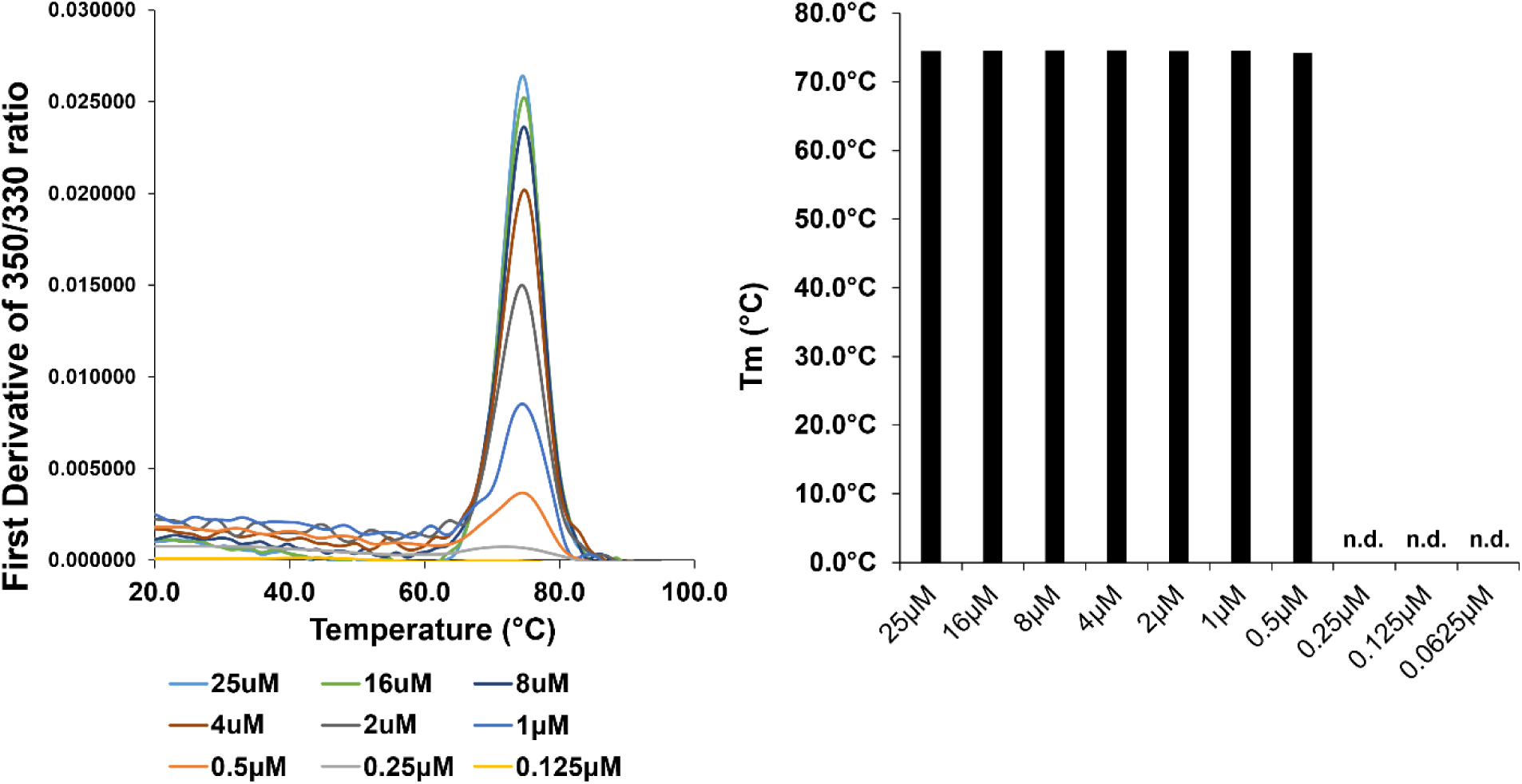
Validation of nanoDSF thermal unfolding measurements. Thermal unfolding of a purified TCR-C α/β complex with a wild-type α/β interface (WT/WT) in 1×PBS (pH 7.4) was measured by nano-differential scanning fluorimetry (nanoDSF) at the indicated protein concentrations. Melting curves show a single cooperative unfolding transition and minimal dependence of the melting temperature (Tm) on protein concentration. Tm values were determined from the peak of the first derivative of the F350/F330 fluorescence ratio. n.d., not determined.

**Supplementary Figure S5.**
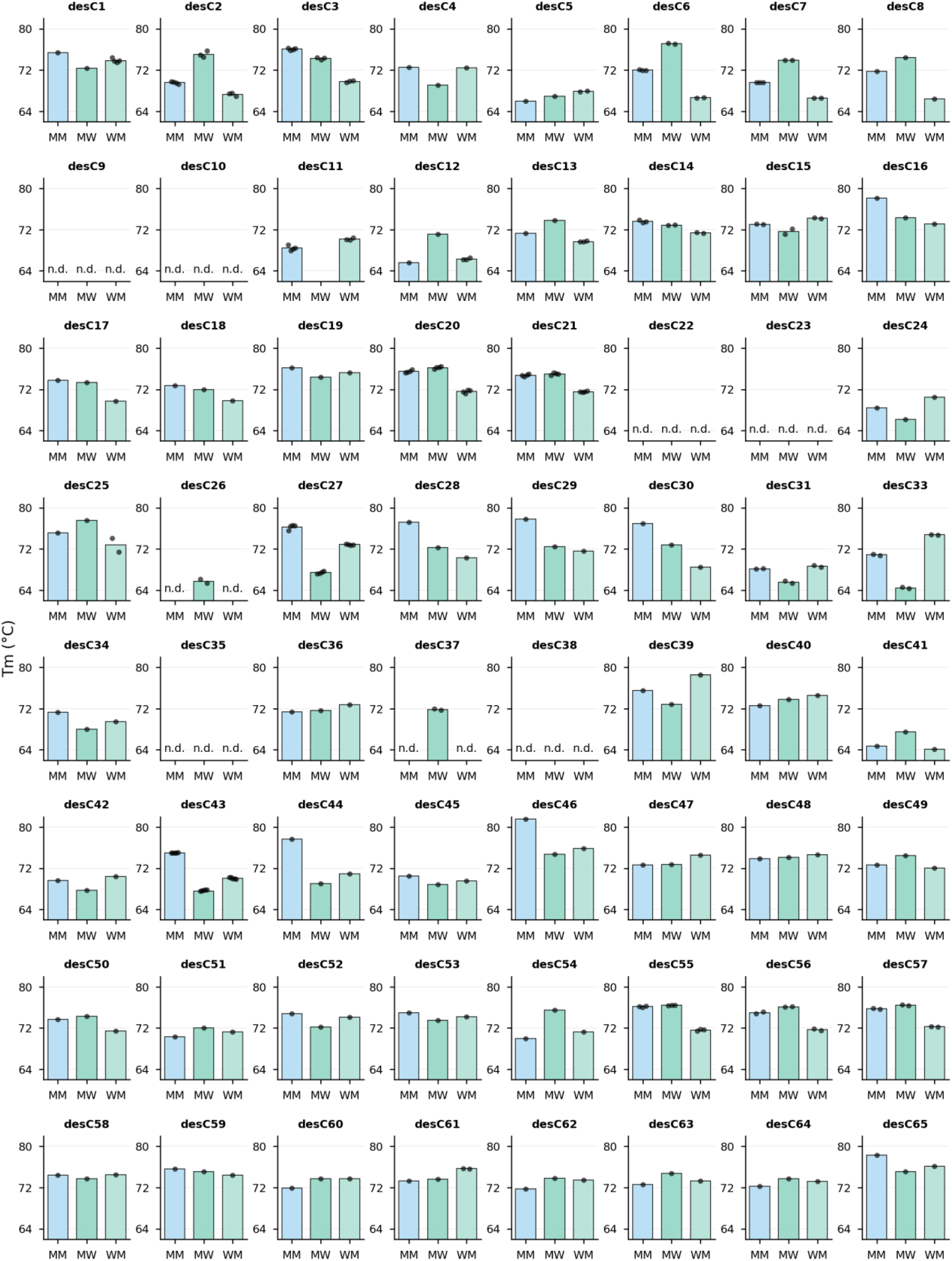
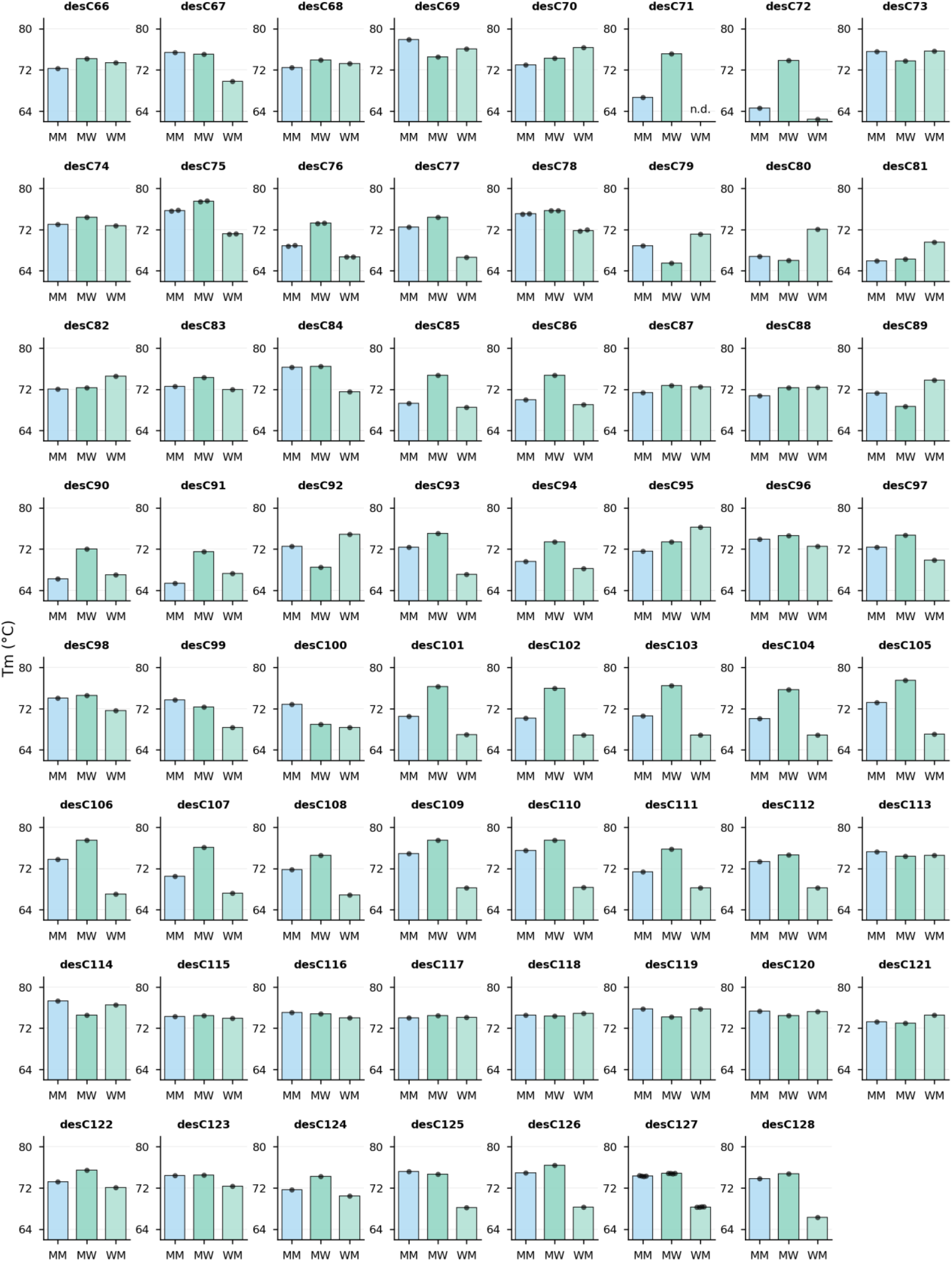
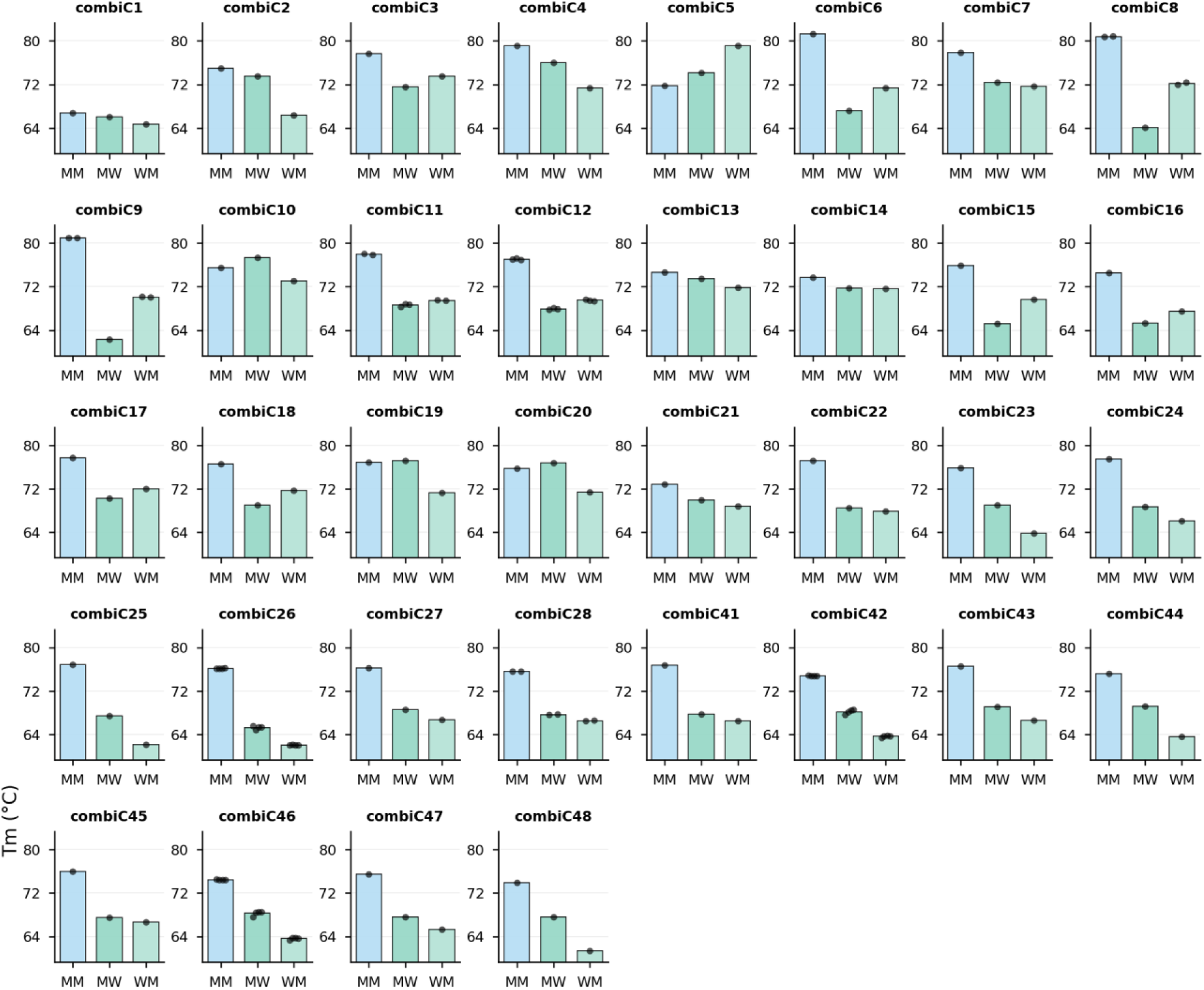
Thermal stability of designed TCR-C α/β complexes. Thermal melting temperatures (Tm) were measured by nano-differential scanning fluorimetry (nanoDSF) for co-expressed and purified TCR constant-domain (TCR-C) α/β complexes across MM, MW, and WM pairing states (MM: mutant α + mutant β, MW: mutant α + wild-type β, WM: wild-type α + mutant β). Data are shown for single-interface designs (desC1–128) and combinatorial TCR-C designs (combiC1–48). Bars represent mean Tm values, with individual measurements shown as dots where applicable. n.d., not determined.

**Supplementary Figure S6.**
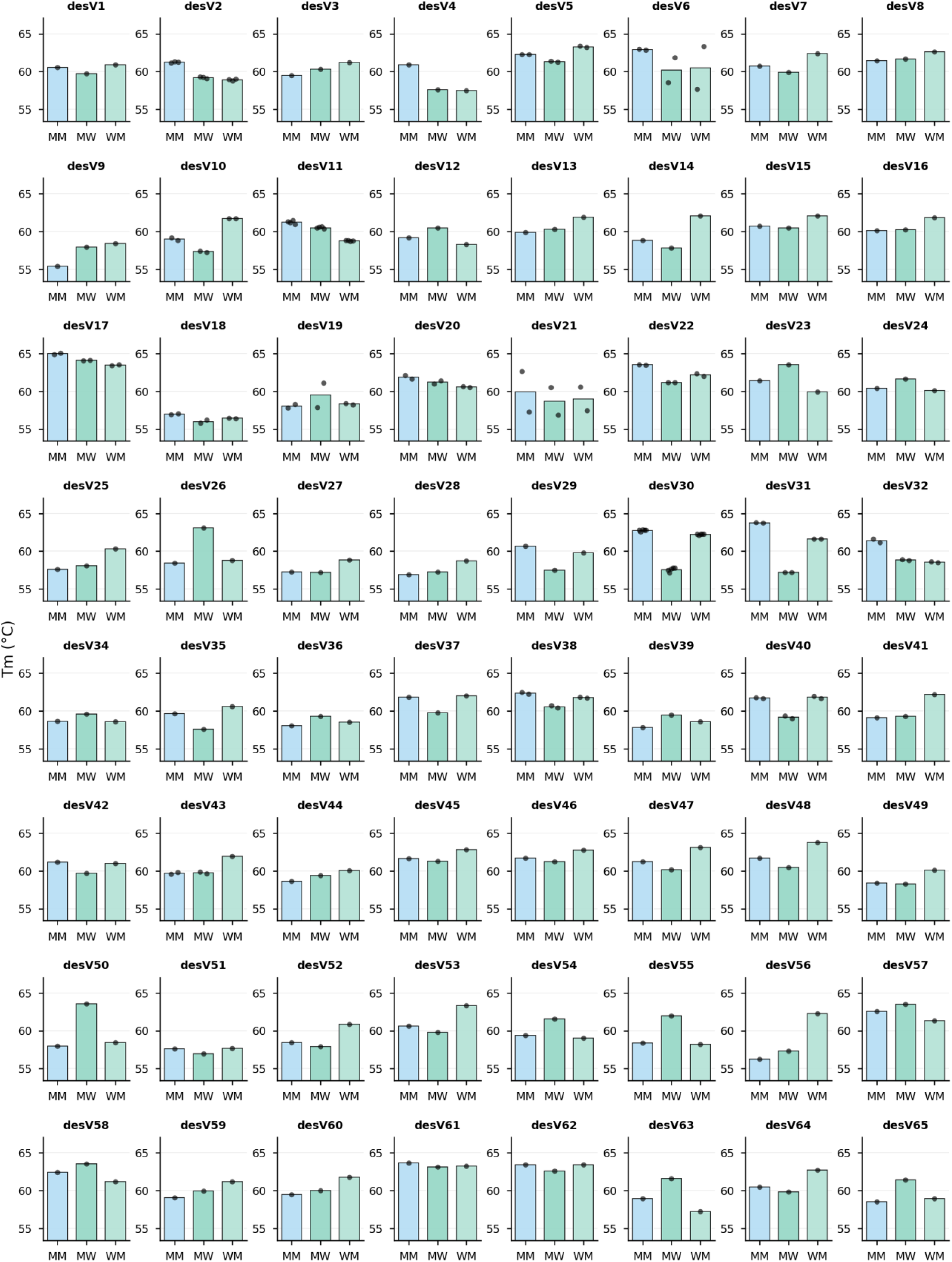
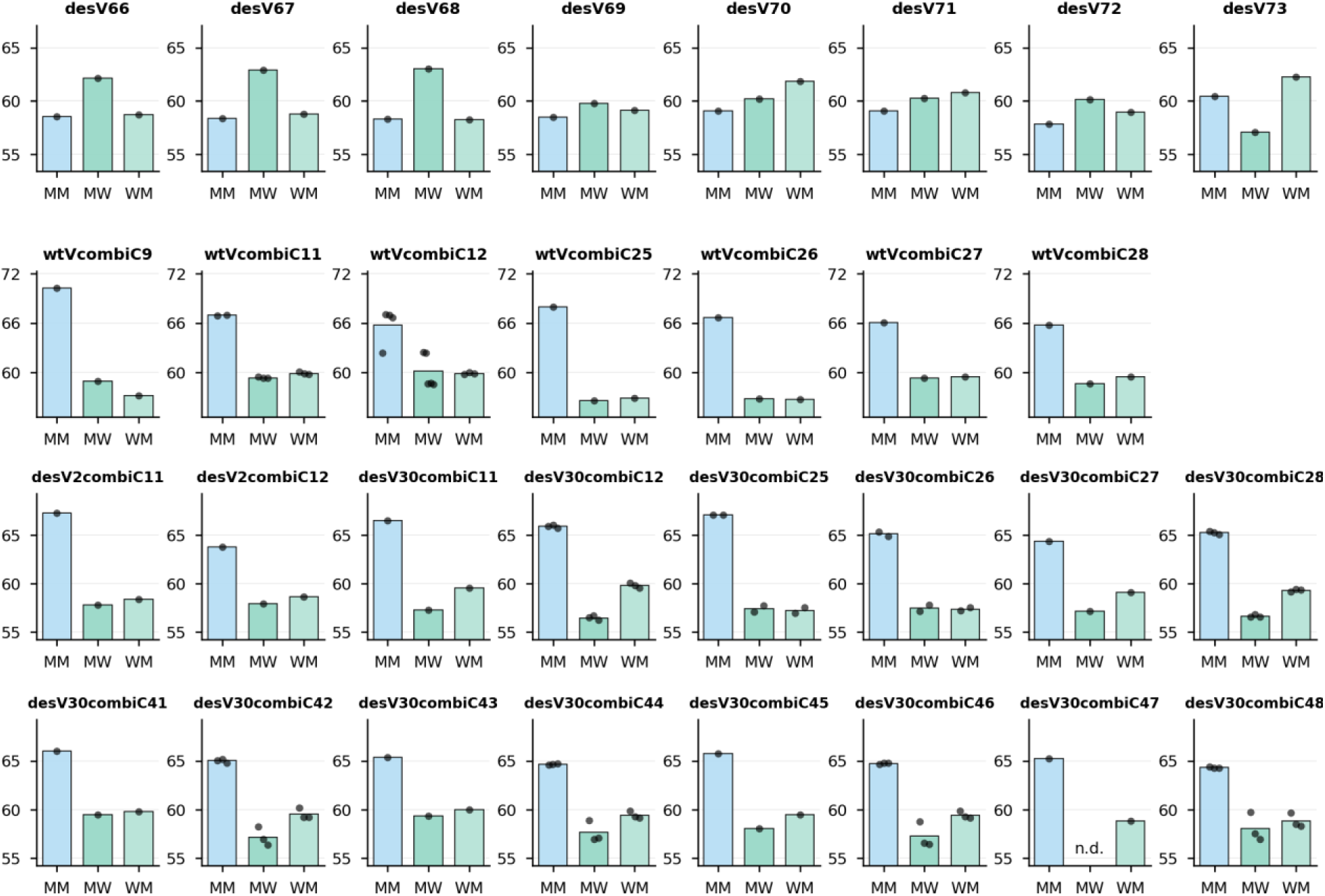
Thermal stability of designed TCR-VC α/β complexes. Thermal melting temperatures (Tm) were measured by nano-differential scanning fluorimetry (nanoDSF) for co-expressed and purified TCR α/β complexes across MM, MW, and WM pairing states (MM: mutant α + mutant β, MW: mutant α + wild-type β, WM: wild-type α + mutant β). Data include variable-domain designs (desV1–73) evaluated in a TCR constant-domain background containing an engineered disulfide but no stabilizing or interface mutations, as well as wild-type or designed variable domains combined with designed constant domains derived from a stabilized TCR-C background (wtVcombiC and desVcombiC). Bars represent mean Tm values, with individual measurements shown as dots where applicable. n.d., not determined.

**Supplementary Figure S7.**
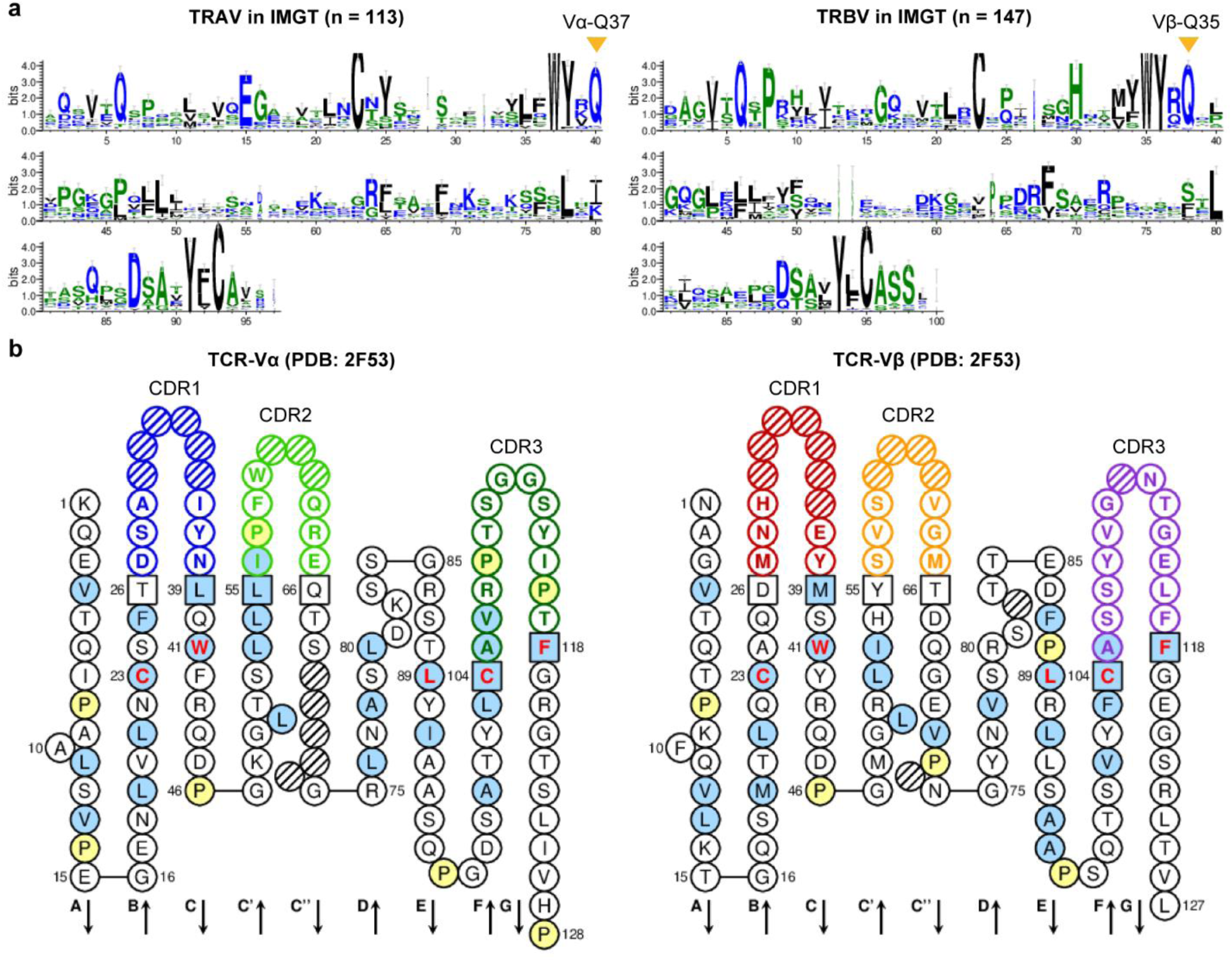
Conservation and topology of TCR-V framework residues. **a**, Sequence logos of human TRAV (n = 113) and TRBV (n = 147) genes from sequences deposited in IMGT, highlighting conserved framework positions relevant to the TCR-V interface design strategy, including the residues mutated in Design V30 (desV30: Vα-Q37 and Vβ-Q35). **b**, Topology diagram illustrating the location of framework and complementarity-determining regions (CDRs) in the TCR-V, based on the structure of the 1G4 TCR variant (PDB: 2F53).

**Supplementary Figure S8.**
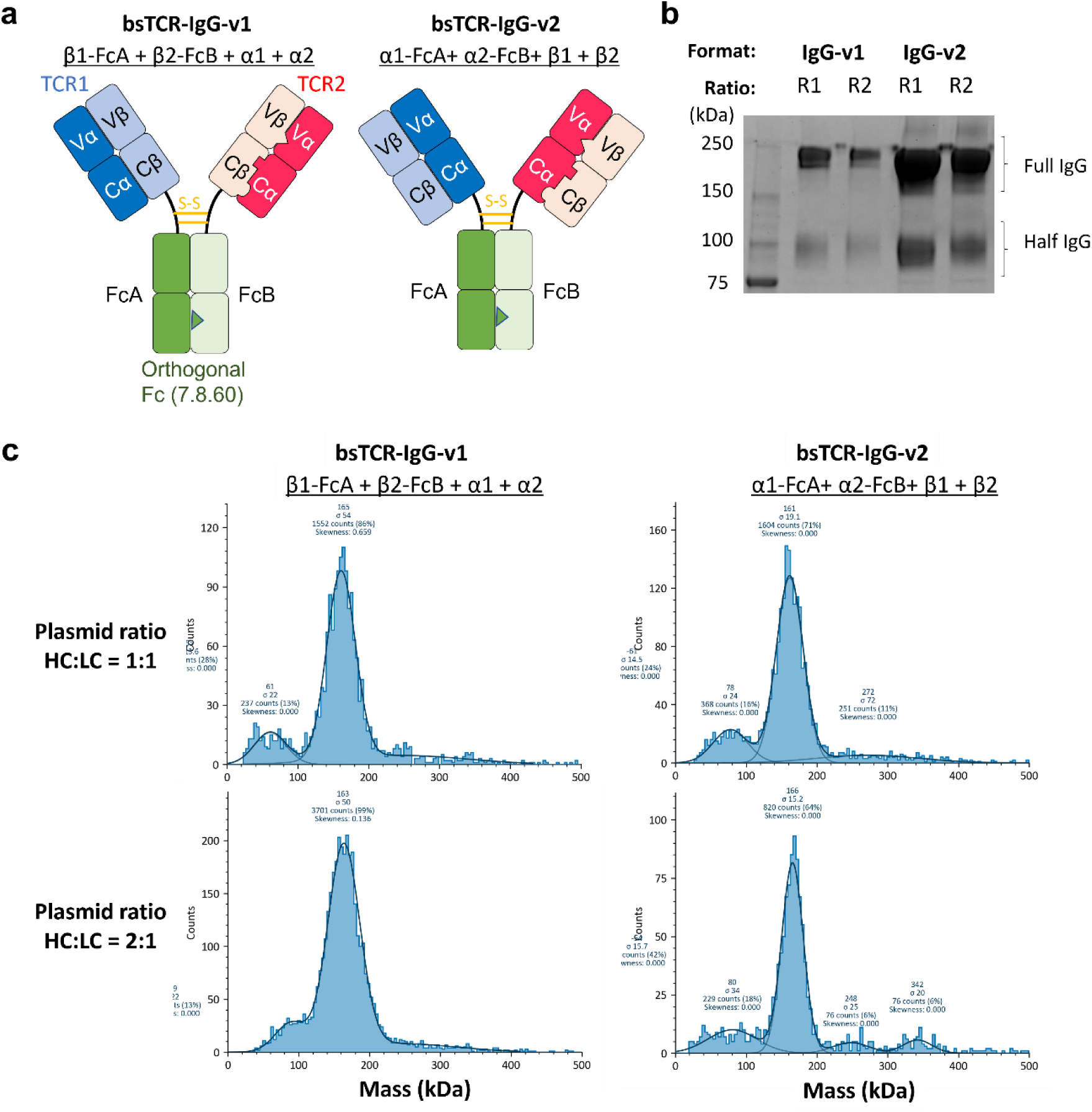
Characterization of a bispecific TCR-IgG architecture. **a**, Schematics of two bispecific TCR-IgG (bsTCR-IgG) formats in which two soluble TCRs are fused to an orthogonal Fc heterodimer (7.8.60); the TCR β chain is Fc-fused in TCR-IgG-v1, whereas the TCR α chain is Fc-fused in TCR-IgG-v2. **b**, SDS-PAGE analysis of bsTCR-IgG expressed using two DNA transfection ratios (R1: Fc-fused chain 1 : Fc-fused chain 2 : unfused chain 1 : unfused chain 2 = 1:1:1:1, R2: 2:2:1:1). Bands corresponding to fully assembled IgG and half-IgG species are indicated. **c**, Mass photometry of bsTCR-IgG produced under R1 and R2, showing mass distributions consistent with differences in assembly state between formats.

**Supplementary Figure S9.**
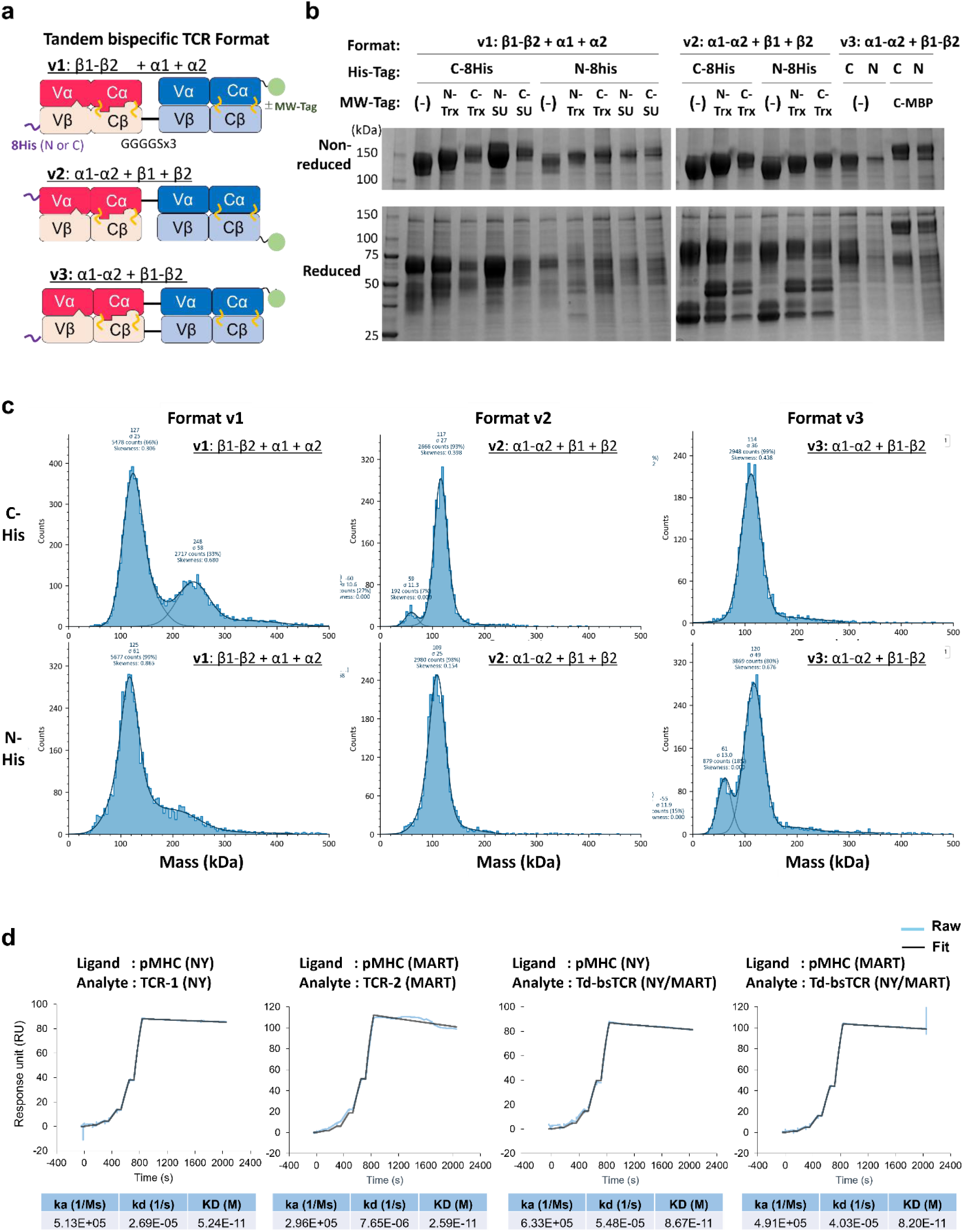
Characterization of a tandem bispecific TCR format. **a**, Schematics of three tandem bispecific TCR architectures (v1–v3), defined by the pattern of TCR chains linked by a flexible GS linker (v1: β1-β2/α1/α2, v2: α1-α2/β1/β2, v3: α1-α2/β1-β2). In all formats, the tandem bispecific TCR comprises two αβ TCRs, one carrying a designed α/β interface (desV30combiC12) and the other retaining a wild-type α/β interface. **b**, SDS-PAGE analysis of expressed tandem bsTCR formats under non-reducing and reducing conditions. To assess sub-chain assembly, selected α or β chains were fused to molecular-weight (MW) shift tags to alter apparent mobility. The TCR variable-constant domains have expected molecular weights of ∼23 kDa (Vα-Cα) and ∼27 kDa (Vβ-Cβ), with apparent migration influenced by glycosylation under reducing SDS-PAGE conditions. MW-shift tags include thioredoxin (Trx, ∼12 kDa), SUMOstar (∼10 kDa), and MBP (∼41 kDa), enabling evaluation of relative sub-chain stoichiometry by reduced SDS-PAGE. **c**, Surface plasmon resonance (SPR) binding of the selected tandem format (v2: α1-α2/β1-β2) to pMHC complexes presenting NY-ESO-1-derived or MART-1-derived peptides, corresponding to the targets of TCR1 and TCR2, respectively.

**Supplementary Figure S10.**
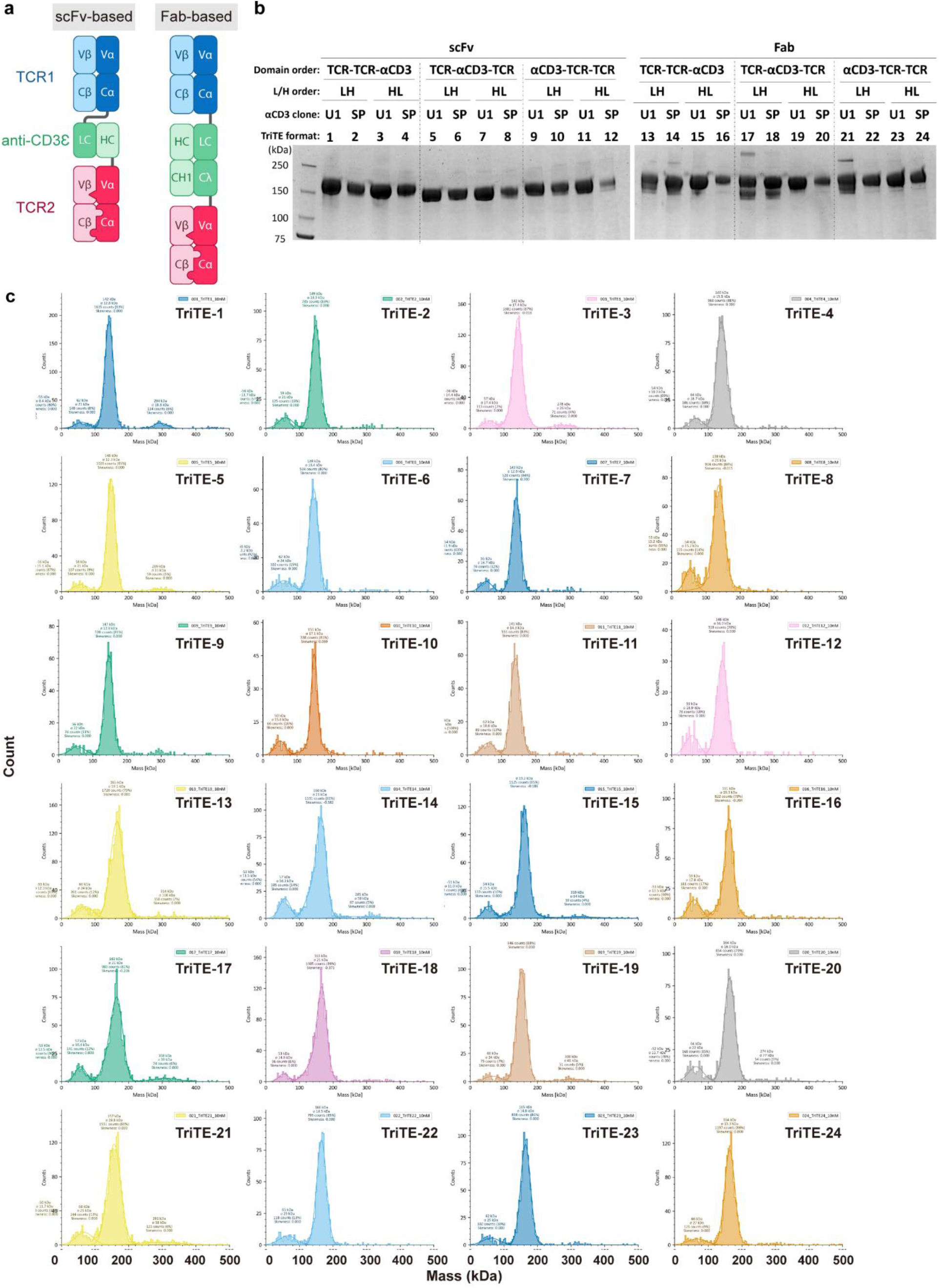
TriTE format design and assembly screening. **a**, Schematics of dual-TCR trispecific T-cell engager (TriTE) architectures comprising two orthogonal TCRs (TCR1 and TCR2) and an anti-CD3ε module configured as either an scFv (TriTE1–12) or a Fab (TriTE13–24). For each anti-CD3 (αCD3) configuration, three domain orders (TCR1-TCR2-αCD3, TCR1-αCD3-TCR2, and αCD3-TCR1-TCR2) were evaluated. Within each domain order, four variants were generated by combining two anti-CD3 clones (U1: humanized UCHT-1, SP: SP34) with two orientations (LH/HL for scFv: VL-VH vs VH-VL order; LH/HL for Fab: LC/HC placement in the main chain). **b**, SDS-PAGE analysis of TriTE variants expressed and purified in Expi293 cells (format numbering and design matrix are indicated in the figure), showing predominant expression of full-length TriTE constructs across formats. **c**, Mass photometry of purified TriTE variants, used to assess assembly state and stoichiometry. Dominant species are consistent with the expected mass, with minor lower-molecular-weight species observed in some formats.

**Supplementary Figure S11.**
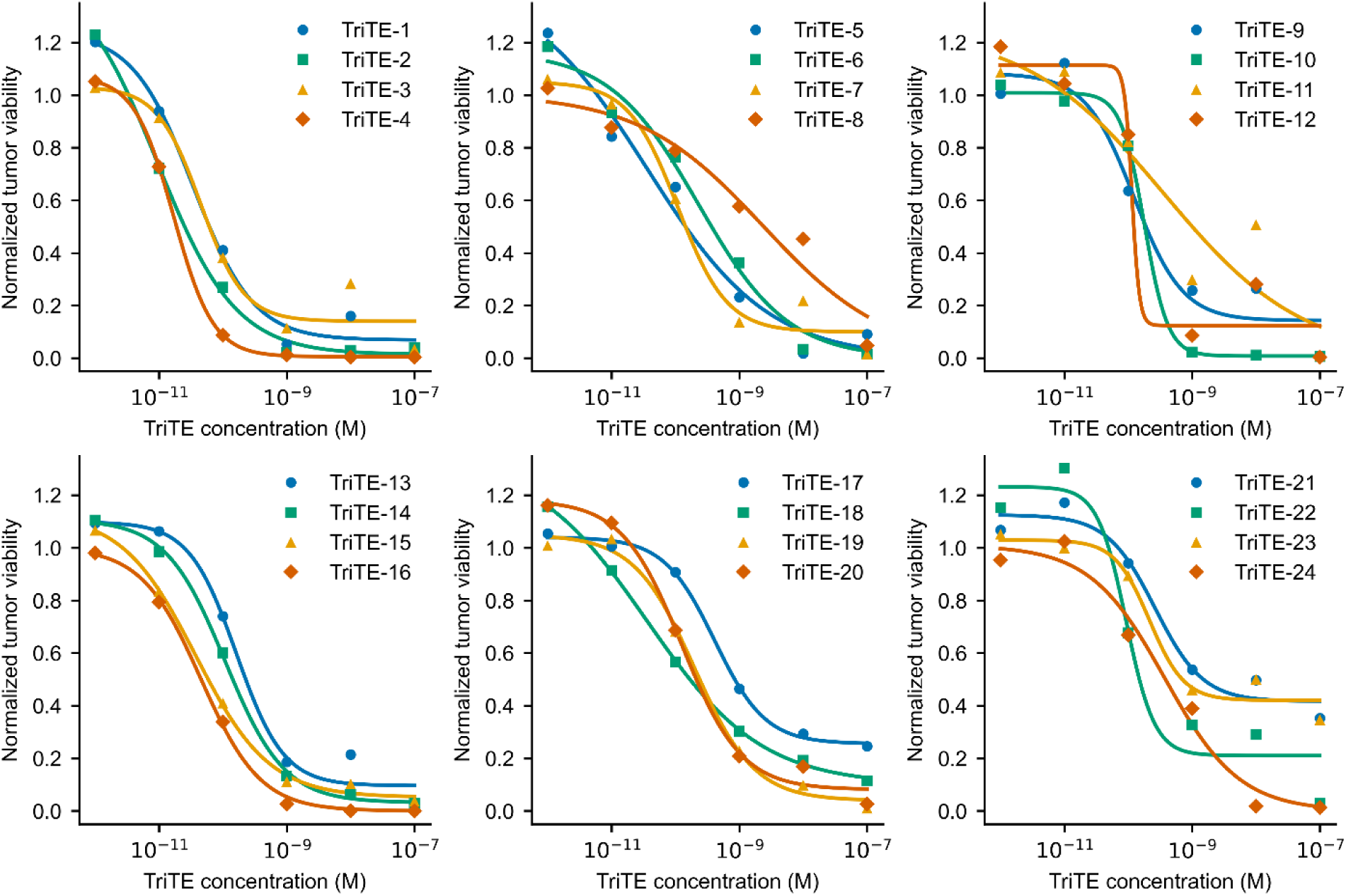
T-cell redirected cytotoxicity screening of TriTE formats. T-cell redirected tumor killing assays using A375 melanoma cells simultaneously pulsed with NY-ESO-1 peptide (SLLMWITQC) and MART-1 peptide (ELAGIGILTV), corresponding to the targets of TCR1 and TCR2, respectively. Dose-response curves (TriTE concentration versus normalized tumor cell viability) and EC50 values are shown to compare cytotoxic potency across TriTE formats. Curves are grouped as TriTE1–4, 5–8, and 9–12 (scFv-based formats), and TriTE13–16, 17–20, and 21–24 (Fab-based formats), matching the domain-order groupings defined in Supplementary Figure S10.

**Supplementary Table S1.** All residue clusters for multistate design (MSD)

| Domain | Cluster_name | Residue_TCRα | Residue_TCRβ | N_residues | PDB |
| --- | --- | --- | --- | --- | --- |
| TCR-C | clusterC1 | 144, 145, 168, 175, 177 | 136, 140, 142, 144, 197 | 10 | 6U07 |
| TCR-C | clusterC1a | 170, 173, 175 | 142 | 4 | 6U07 |
| TCR-C | clusterC1b | 144, 145, 168, 170, 175, 177 | 136, 140, 142, 144, 197 | 11 | 6U07 |
| TCR-C | clusterC1c | 144, 145, 177 | 136, 144, 197 | 6 | 6U07 |
| TCR-C | clusterC1d | 126 | 136 | 2 | 6U07 |
| TCR-C | clusterC1e | 145, 168, 175, 177 | 140, 142, 144, 197 | 8 | 6U07 |
| TCR-C | clusterC1f | 145 | 197 | 2 | 6U07 |
| TCR-C | clusterC1g | 128, 142, 144, 145, 177 | 136, 140, 144, 197 | 9 | 6U07 |
| TCR-C | clusterC2 | 124, 125, 205 | 139 | 4 | 6U07 |
| TCR-C | clusterC2a | 124, 205 | 139 | 3 | 6U07 |
| TCR-C | clusterC2b | 124, 128, 205 | 139 | 4 | 6U07 |
| TCR-C | clusterC2c | 124, 128, 205, 207, 208 | 133, 135, 136, 139 | 9 | 6U07 |
| TCR-C | clusterC3 | 128, 140, 142, 181, 183 | 130, 146, 148 | 8 | 6U07 |
| TCR-C | clusterC3a | 140, 181, 183 | 130, 146, 148 | 6 | 6U07 |
| TCR-C | clusterC4 | 163, 181 | 175, 193, 195 | 5 | 6U07 |
| TCR-C | clusterC4a | 163, 179, 181 | 175, 193, 195 | 6 | 6U07 |
| TCR-C | clusterC4b | 163, 179, 180, 181 | 175, 193, 195 | 7 | 6U07 |
| TCR-C | clusterC-α124 | 124, 126, 128, 205 | 139, 140 | 6 | 6U07 |
| TCR-C | clusterC-α133 | 133 | 129 | 2 | 6U07 |
| TCR-C | clusterC-α134 | 132, 133, 134 | 129, 131, 239, 240, 241 | 8 | 6U07 |
| TCR-C | clusterC-α168 | 168, 169 | 171, 172 | 4 | 6U07 |
| TCR-C | clusterC-α171 | 169, 170, 171, 172, 173, 174 | 169, 170, 171 | 9 | 6U07 |
| TCR-C | clusterC-α173 | 170, 173 | 199 | 3 | 6U07 |
| TCR-C | clusterC-α181 | 140, 163, 180, 181, 183 | 146, 148, 175, 193, 195 | 10 | 6U07 |
| TCR-C | clusterC-α205 | 205, 207, 208 | 133, 135, 136 | 6 | 6U07 |
| TCR-C | clusterC-β128 | 132, 138 | 128 | 3 | 6U07 |
| TCR-C | clusterC-β130 | 132, 138, 140 | 130, 148 | 5 | 6U07 |
| TCR-C | clusterC-β135 | 128, 205, 207, 208 | 135 | 5 | 6U07 |
| TCR-C | clusterC-β146 | 130, 140, 181 | 146 | 4 | 6U07 |
| TCR-C | clusterC-β170 | 169, 171, 172 | 170, 171 | 5 | 6U07 |
| TCR-C | clusterC-β171 | 168, 169 | 171 | 3 | 6U07 |
| TCR-C | clusterC-β177 | 164, 165 | 176, 177 | 4 | 6U07 |
| TCR-C | clusterC-β179 | 161, 162 | 179, 181 | 4 | 6U07 |
| TCR-C | clusterC-β182 | 158, 159, 161 | 181, 182, 183, 184 | 7 | 6U07 |
| TCR-C | clusterC-β191 | 183 | 191, 192 | 3 | 6U07 |
| TCR-C | clusterC-β239 | 133, 134 | 239, 240, 241 | 5 | 6U07 |
| TCR-V | clusterV1 | 40, 41, 42 | 7, 104, 107 | 6 | 2F53 |
| TCR-V | clusterV1a | 41 | 104 | 2 | 2F53 |
| TCR-V | clusterV2 | 37, 43, 88 | 35, 41, 88 | 6 | 2F53 |
| TCR-V | clusterV2a | 37 | 35 | 2 | 2F53 |
| TCR-V | clusterV3 | 105 | 38, 39 | 3 | 2F53 |
| TCR-V | clusterV4 | 101, 103 | 33 | 3 | 2F53 |
| TCR-V | clusterV4a | 35, 43, 101, 103 | 33, 41, 102 | 7 | 2F53 |
| TCR-V | clusterV5 | 100 | 29, 43 | 3 | 2F53 |
| TCR-V | clusterV6 | 31, 33, 45, 48 | 99 | 5 | 2F53 |
| TCR-V | clusterV6a | 45, 48 | 99 | 3 | 2F53 |
| TCR-V | clusterV7 | 35 | 100, 101, 102 | 4 | 2F53 |

**Supplementary Table S2.** Orthogonal designs of the TCR-C (desC1–128).

| Design_name | Designed mutations_α | Designed mutations_β | Design_type |
| --- | --- | --- | --- |
| desC1 | Y128F T144M D145F L168A F175L S177L | K142L R197I | MSD+rational |
| desC2 | T144M L168A F175K S177M | T140K K142D R197I | MSD+rational |
| desC3 | T144M D145N F175R S177L | T140Y K142E R197T | MSD |
| desC4 | Y128F D145M L168I F175I S177V | K142Q R197M | MSD+rational |
| desC5 | Y128F D145K L168I F175V S177I | K142E R197M | MSD+rational |
| desC6 | D124R F205E | R139D | MSD+rational |
| desC7 | D124E F205K | R139D | MSD+rational |
| desC8 | D124Q F205K | R139D | MSD+rational |
| desC9 | V140T V181L W183K | V146A L148W | MSD+rational |
| desC10 | V140M L142M V181A | V146G L148F | MSD+rational |
| desC11 | S179I V181Q | D175Q R195A | MSD+rational |
| desC12 | K134E | E240K | rational |
| desC13 | F175K | K142D | rational/SSS |
| desC14 | S179A | R195M | rational |
| desC15 | K134E | E126R | rational |
| desC16 | T144M D145N F175R S177L | E136Q T140Y K142E R197T | MSD |
| desC17 | T144E D145N F175R S177K | E136Q T140Y K142E R197S | MSD |
| desC18 | D145H F175R S177Q | T140Y K142E R197T | MSD |
| desC19 | S177H | K142W R197T | MSD |
| desC20 | D124F F205W | R139L | MSD |
| desC21 | D124Q F205K | R139L | MSD |
| desC22 | V140M L142M V181A W183T | F130Y V146G L148W | MSD |
| desC23 | V140T L142S V181L W183K | F130Y V146A L148W | MSD |
| desC24 | T163G V181R | D175L R195A | MSD |
| desC25 | Y128H V140A L142S V181M | V146A L148F | MSD |
| desC26 | V140T V181A W183R | L148W | MSD |
| desC27 | S179R | R195T | MSD |
| desC28 | S179I V181I | R195A | MSD |
| desC29 | S179L V181I | D175M R195A | MSD |
| desC30 | S179I V181I | D175E R195A | MSD |
| desC31 | W183S | L148A S193Y | MSD |
| desC32 | V181M W183Q | L148W D175I R195A | MSD |
| desC33 | V140L W183G | L148A D175I S193Y R195M | MSD |
| desC34 | T163G V181L | V146A S193F R195L | MSD |
| desC35 | V181Q W183K | L148W D175V R195A | MSD |
| desC36 | F205R P207G | A135E | MSD |
| desC37 | F205K P207G | A135E | MSD |
| desC38 | L130A V181Q | V146Q | MSD |
| desC39 | Y161V | L179M E181Y | MSD |
| desC40 | Y161I | L179M E181H | MSD |
| desC41 | S179R | R195G | previous report |
| desC42 | S179R | R195A | MSD |
| desC43 | S179R | R195S | MSD |
| desC44 | T163G S179I V181I | D175L R195A | MSD |
| desC45 | T163G S179I V181I | D175L S193A R195A | MSD |
| desC46 | T163M S179L | D175A S193I R195T | MSD |
| desC47 | S133H |  | MSD |
| desC48 | S133V |  | MSD |
| desC49 | R171P S172W | D170A | MSD |
| desC50 | R171D S172W | D170K | MSD |
| desC51 | D132L | A128G | MSD |
| desC52 | D132A K138L | A128I | MSD |
| desC53 | D132S K138L | A128I | MSD |
| desC54 | D132L K138R | A128G | MSD |

**Supplementary Table S2. Orthogonal designs of the TCR-C (desC1–128).**
| Design_name | Designed mutations_α | Designed mutations_β | Design_type |
| --- | --- | --- | --- |
| desC55 | R171E S172D | D170K | MSD |
| desC56 | R171Q S172D | D170K | MSD |
| desC57 | R171E S172D | D170R | MSD |
| desC58 | D164N | Q177E | MSD |
| desC59 | D164N K165T | Q177E | MSD |
| desC60 | Y161V | L179M E181H | MSD |
| desC61 | Y161V | L179M E181F | MSD |
| desC62 | Y161V | E181H Q182Y | MSD |
| desC63 | Y161I | E181H Q182Y | MSD |
| desC64 | Y161V | E181W Q182Y | MSD |
| desC65 | F175R S177M | K142T R197T | MSD |
| desC66 | T144M F175D S177L | R197S | MSD |
| desC67 | T144M D145N F175K S177L | K142D R197T | MSD |
| desC68 | T144E F175K S177L | E136Q K142D R197T | MSD |
| desC69 | F175R S177L | K142T R197T | MSD |
| desC70 | T144M F175D S177L | R197V | MSD |
| desC71 | S177H | K142W T144I R197T | MSD |
| desC72 | S177T | K142Y T144I R197G | MSD |
| desC73 | S177A | K142L R197T | MSD |
| desC74 | L168I F175V S177Q | K142Q R197M | MSD |
| desC75 | D124R F205E | R139N | MSD |
| desC76 | F205K | R139D | MSD |
| desC77 | D124A F205R | R139D | MSD |
| desC78 | D124W F205W | R139L | MSD |
| desC79 | V181Q | D175Q R195A | MSD |
| desC80 | A126Q Y128V F205K | R139L | MSD |
| desC81 | Y128M F205K | R139L T140Y | MSD |
| desC82 | S133R |  | MSD |
| desC83 | R171P S172W | D170A | MSD |
| desC84 | R171E S172D | D170K | MSD |
| desC85 | P207G | S133T A135Q | MSD |
| desC86 | P207G | S133T A135E | MSD |
| desC87 | F205R P207G | A135Q | MSD |
| desC88 | F205R P207G | A135D | MSD |
| desC89 | S133E | E131A | SSS |
| desC90 | K134E | E240R | rational |
| desC91 | K134D | E240K | rational |
| desC92 | K138E | E126R | rational |
| desC93 | D124T F205K | R139D | SSS |
| desC94 | F205K | R139E | SSS |
| desC95 | F175E S177E | R197V | SSS |
| desC96 | F175R S177Q | T140K K142E R197T | MSD |
| desC97 | F175K S177L | K142D R197T | rational |
| desC98 | F175R S177L | K142E R197T | rational |
| desC99 | F175R S177K | K142E R197S | rational |
| desC100 | D145H F175R S177K | K142E R197S | rational |
| desC101 | D124F | R139D | SSS |
| desC102 | D124W | R139D | SSS |
| desC103 | D124F F205W | R139D | rational |
| desC104 | D124W F205W | R139D | rational |
| desC105 | D124K | R139D | rational |
| desC106 | D124R | R139D | rational |
| desC107 | D124Y | R139D | rational |

**Supplementary Table S2. Orthogonal designs of the TCR-C (desC1–128).**
| **Design_name** | **Designed mutations_α** | **Designed mutations_β** | **Design_type** |
| --- | --- | --- | --- |
| desC108 | D124Y F205K | R139D | rational |
| desC109 | D124K | R139E | SSS |
| desC110 | D124R | R139E | rational |
| desC111 | D124W | R139E | rational |
| desC112 | D124S F205R | R139E | rational |
| desC113 | S158R | A184D | SSS |
| desC114 | S158R | A184D L185R | SSS |
| desC115 | S158D | A184K | SSS |
| desC116 | S158E | A184K | SSS |
| desC117 | S158Y | A184K | SSS |
| desC118 | S158Y | A184E | SSS |
| desC119 | S158W | L185R | SSS |
| desC120 | S158W | L185K | SSS |
| desC121 | S172R | F202E | SSS |
| desC122 | S172E | F202R | rational |
| desC123 | F175R | K142E | SSS |
| desC124 | M173Y F175K | K142D | SSS |
| desC125 | D124R F205R | R139E | rational |
| desC126 | D124R F205H | R139E | rational |
| desC127 | D124R F205K | R139E | rational |
| desC128 | D124R F205K | R139D | rational |

**Supplementary Table S3.** Combination designs of the TCR-C (combiC1–48).

| **Design_name** | **Design_combination** |
| --- | --- |
| combiC1 | desC3_11 |
| combiC2 | desC3_14 |
| combiC3 | desC16_15 |
| combiC4 | desC16_20 |
| combiC5 | desC16_21 |
| combiC6 | desC16_27 |
| combiC7 | desC16_15_20 |
| combiC8 | desC16_15_27 |
| combiC9 | desC16_15_20_27 |
| combiC10 | desC27_15 |
| combiC11 | desC27_20 |
| combiC12 | desC27_21 |
| combiC13 | desC15_20 |
| combiC14 | desC15_21 |
| combiC15 | desC15_21_27 |
| combiC16 | desC15_21_43 |
| combiC17 | desC55_27 |
| combiC18 | desC56_27 |
| combiC19 | desC55_21 |
| combiC20 | desC56_21 |
| combiC21 | desC27_21_95 |
| combiC22 | desC27_21_96 |
| combiC23 | desC27_21_97 |
| combiC24 | desC27_21_98 |
| combiC25 | desC27_21_99 |
| combiC26 | desC27_21_100 |
| combiC27 | desC43_20 |
| combiC28 | desC43_21 |
| combiC29 | desC27_77 |
| combiC30 | desC43_77 |
| combiC31 | desC27_93 |
| combiC32 | desC43_93 |
| combiC33 | desC27_99 |
| combiC34 | desC43_99 |
| combiC35 | desC27_100 |
| combiC36 | desC43_100 |
| combiC37 | desC27_110 |
| combiC38 | desC43_110 |
| combiC39 | desC27_112 |
| combiC40 | desC43_112 |
| combiC41 | desC27_125 |
| combiC42 | desC43_125 |
| combiC43 | desC27_126 |
| combiC44 | desC43_126 |
| combiC45 | desC27_127 |
| combiC46 | desC43_127 |
| combiC47 | desC27_128 |
| combiC48 | desC43_128 |

**Supplementary Table S4.** Orthogonal designs of the TCR-V (desV1–73).

| Design_name | Designed mutations_Vα | Designed mutations_Vβ | Design_type |
| --- | --- | --- | --- |
| desV1 | K41D | E104R | MSD |
| desV2 | Q37E | Q35K | MSD |
| desV3 | R105E | M39R | MSD |
| desV4 | F103E | L41G F102Y | MSD |
| desV5 | K41Q | R107K | MSD |
| desV6 | K41R G42A | R107K | MSD |
| desV7 | K41D | E104S | MSD |
| desV8 | K41Q |  | MSD |
| desV9 | Q37N | Q35R | MSD |
| desV10 | Q37R | Q35S | rational |
| desV11 | Q37Y | Q35K | rational |
| desV12 | Q37Y | Q35R | rational |
| desV13 | R105D | M39T | MSD/SSS |
| desV14 | R105S | M39T | MSD |
| desV15 | R105D | M39K | MSD |
| desV16 | R105E | M39S | MSD |
| desV17 | P101W F103M | Y33W | MSD |
| desV18 | P101Y F103S | Y33A L41I | MSD |
| desV19 | I100L | L43T | MSD |
| desV20 | I100L | L43S | MSD |
| desV21 | I100L | Y29F L43A | MSD |
| desV22 | I100L | Y29W L43T | MSD |
| desV23 | N31L L48V | E99T | MSD |
| desV24 | L48S | E99T | MSD |
| desV25 | S45L L48S | E99T | MSD |
| desV26 | L48F | E99F | MSD |
| desV27 | F35A | F102W | MSD |
| desV28 | F35N | F102W | MSD |
| desV29 | Q37K | Q35E | rational |
| desV30 | Q37K | Q35Y | rational |
| desV31 | Q37K | Q35D | rational |
| desV32 | Q37D | Q35K | rational |
| desV33 | Q37R | Q35T | rational |
| desV34 | Q37T | Q35R | rational |
| desV35 | Q37R | Q35A | MSD |
| desV36 | Q37A | Q35R | MSD |
| desV37 | Q37I | Q35M | MSD |
| desV38 | Q37L | Q35M | MSD |
| desV39 | Q37S | Q35R | MSD |
| desV40 | Q37V | Q35M | MSD |
| desV41 | K41D | E104D | MSD |
| desV42 | K41D | E104K | MSD |
| desV43 | K41D | E104T | MSD |
| desV44 | K41D | E104V | MSD |
| desV45 | K41Q | F8Y | rational |
| desV46 | K41Q | K7R | rational |
| desV47 | K41T | V86L | rational |
| desV48 | K41T | V86W | rational |

**Supplementary Table S4. Orthogonal designs of the TCR-V (desV1–73).**
| Design_name | Designed mutations_Vα | Designed mutations_Vβ | Design_type |
| --- | --- | --- | --- |
| desV49 | L43A | L41F | MSD |
| desV50 | L43F | L41A | MSD |
| desV51 | Q37V L43A | Q35R L41M | MSD |
| desV52 | Q37V L43A | L41M | MSD |
| desV53 | G40E | R107K | SSS |
| desV54 | G40K | R107F | SSS |
| desV55 | F103W | R42I | SSS |
| desV56 | F103Y | S31A | SSS |
| desV57 | S108K | G38D | SSS |
| desV58 | S108K | G38E | SSS |
| desV59 | R105D | M39N | SSS |
| desV60 | R105D | M39R | SSS |
| desV61 | T86L | M39W | SSS |
| desV62 | T86V | M39Y | SSS |
| desV63 | S45K | F101D | SSS |
| desV64 | S45P | F101V | SSS |
| desV65 | G40M | R107F D150P | SSS |
| desV66 | G40N | R107F D150P | SSS |
| desV67 | G40R | R107F D150P | SSS |
| desV68 | G40W | R107I D150P | SSS |
| desV69 | R105D | M39K R42I | SSS |
| desV70 | R105D | G38D M39K | SSS |
| desV71 | R105D | G38N M39K | SSS |
| desV72 | R105D | M39R R42I | SSS |
| desV73 | Q37R | Q35Y | rational |

**Supplementary Table S5.** Representative orthogonal design of TCR-C (desC and combiC).

| Design (TCR-C) | Design type | Mutation |  | T<sub>m</sub> (°C) |  |  |
| --- | --- | --- | --- | --- | --- | --- |
|  |  | chain A | chain B | MM (WW) | MW | WM |
| stTCR-C | (no design) | NA | NA | 74.4°C | NA | NA |
| desC20 | charge to hydrophobic | chA D124F F205W | chB R139L | 75.3°C | 75.9°C | 71.5°C |
| desC21 | charge to hydrophobic | chA D124Q F205K | chB R139L | 74.7°C | 74.7°C | 71.5°C |
| desC27 | knob-into-hole | chA S179R | chB R195T | 75.6°C | 67.2°C | 73.0°C |
| desC43 | knob-into-hole | chA S179R | chB R195S | 75.0°C | 67.6°C | 69.9°C |
| desC56 | charge swap | chA R171Q S172D | chB D170K | 74.8°C | 76.1°C | 71.8°C |
| desC99 | charge swap | chA F175R S177K | chB K142E R197S | 73.7°C | 72.3°C | 68.4°C |
| desC100 | charge swap | chA D145H F175R S177K | chB K142E R197S | 72.9°C | 69.0°C | 68.4°C |
| desC125 | charge swap | chA D124R F205R | chB R139E | 75.2°C | 74.7°C | 68.2°C |
| desC127 | charge swap | chA D124R F205K | chB R139E | 74.5°C | 74.8°C | 68.3°C |
| desC128 | charge swap | chA D124R F205K | chB R139D | 73.8°C | 74.8°C | 66.3°C |
| combiC12 (desC21_C27) | combination | chA D124Q S179R F205K | chB R139L R195T | 77.2°C | 68.1°C | 69.5°C |
| combiC25 (desC21_27_99) | combination | chA D124Q F175R S177K S179R F205K | chB R139L K142E R195T R197S | 76.9°C | 67.4°C | 62.2°C |
| combiC26 (desC21_27_100) | combination | chA D124Q D145H F175R S177K S179R F205K | chB R139L K142E R195T R197S | 76.1°C | 65.5°C | 62.1°C |
| combiC42 (desC43_125) | combination | chA D124R S179R F205R | chB R139E R195S | 74.9°C | 67.6°C | 63.4°C |
| combiC46 (desC43_127) | combination | chA D124R S179R F205K | chB R139E R195S | 74.5°C | 67.6°C | 63.3°C |
| combiC48 (desC43_128) | combination | chA D124R S179R F205K | chB R139D R195S | 73.9°C | 67.6°C | 61.3°C |

TCR-C: TCR constant domain
stTCR-C: wild type TCR variable domain and TCR constant domain with stabilizing mutations
desC: TCR constant domain design
combiC: combinative design of TCR constant
T<sub>m</sub>: melting temperature determined by nanoDSF
MM: mutant/mutant
MW: mutant/wild type
WM: wild type/mutant
NA: Not applicable

**Supplementary Table S6.** Representative orthogonal design of TCR-V (desV).

| Design (TCR variant) | Design type | Mutation |  | T<sub>m</sub> (°C) |  |  |
| --- | --- | --- | --- | --- | --- | --- |
|  |  | chain A | chain B | MM (WW) | MW | WM |
| wtVdsC | (no design) | NA | NA | 62.5°C | NA | NA |
| desV11dsC | polar to cation-pi | chA Q37Y | chB Q35K | 61.3°C | 60.5°C | 58.8°C |
| desV30dsC | polar to cation-pi | chA Q37K | chB Q35Y | 62.8°C | 57.5°C | 62.3°C |
| desV31dsC | polar to charge | chA Q37K | chB Q35D | 63.8°C | 57.2°C | 61.6°C |
| desV32dsC | polar to charge | chA Q37D | chB Q35K | 61.6°C | 58.9°C | 58.6°C |
| desV38dsC | polar to hydrophobic | chA Q37L | chB Q35M | 62.5°C | 60.7°C | 61.8°C |
| desV40dsC | polar to hydrophobic | chA Q37V | chB Q35M | 61.8°C | 59.3°C | 62.0°C |
| desV58dsC | new salt bridge | chA S108K | chB G38E | 62.4°C | 63.6°C | 61.2°C |

desV: TCR variable domain design
wtVdsC: wild type TCR variable domain and TCR constant domain with an extra disulfide bond (Cα -T166C and Cβ -S173C)
T<sub>m</sub>: melting temperature determined by nano DSF
MM: mutant/mutant
MW: mutant/wild type
WM: wild type/mutant
NA: Not applicable

**Supplementary Table S7.**
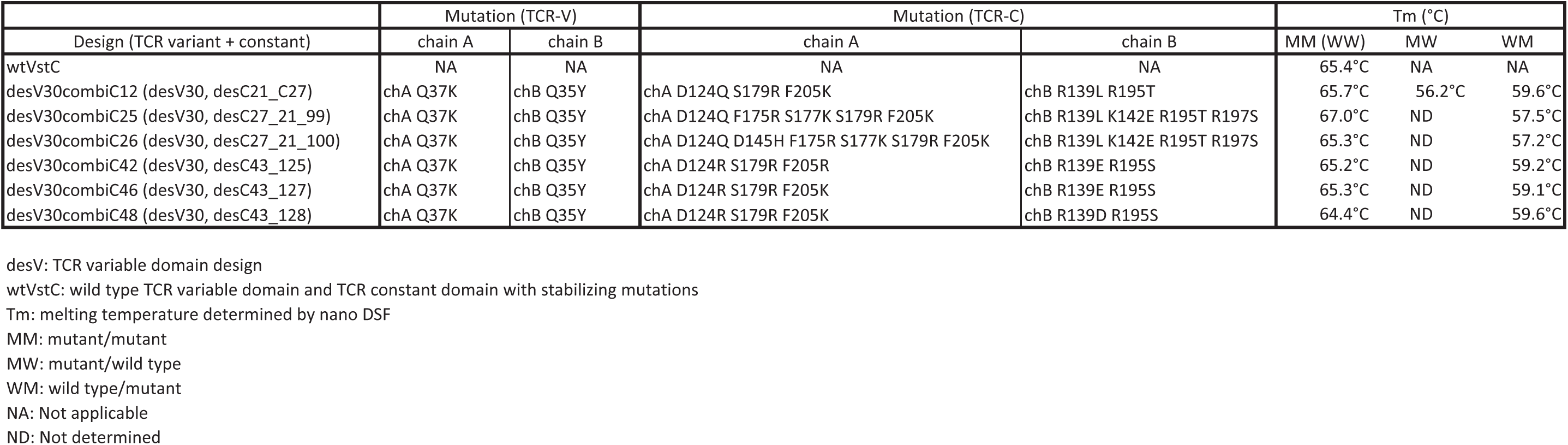
Representative orthogonal design of TCR-VC (wtVcombiC and desVcombiC).

**Supplementary Table S8.**
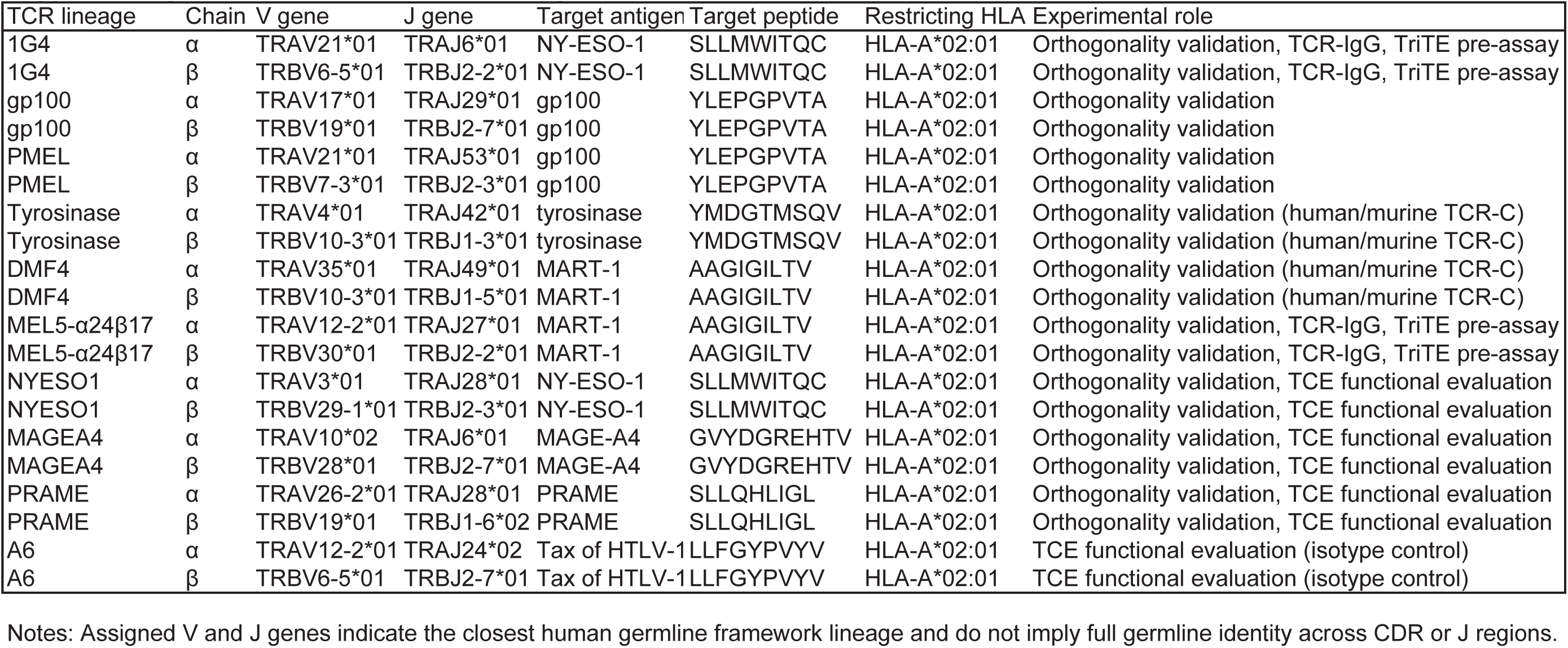
TCRs used in this study and their framework annotations.

| TCR lineage | Chain | V gene | J gene | Target antigen | Target peptide | Restricting HLA | Experimental role |
| --- | --- | --- | --- | --- | --- | --- | --- |
| 1G4 | α | TRAV21*01 | TRAJ6*01 | NY-ESO-1 | SLLMWITQC | HLA-A*02:01 | Orthogonality validation, TCR-IgG, TriTE pre-assay |
| 1G4 | β | TRBV6-5*01 | TRBJ2-2*01 | NY-ESO-1 | SLLMWITQC | HLA-A*02:01 | Orthogonality validation, TCR-IgG, TriTE pre-assay |
| gp100 | α | TRAV17*01 | TRAJ29*01 | gp100 | YLEPGPVTA | HLA-A*02:01 | Orthogonality validation |
| gp100 | β | TRBV19*01 | TRBJ2-7*01 | gp100 | YLEPGPVTA | HLA-A*02:01 | Orthogonality validation |
| PMEL | α | TRAV21*01 | TRAJ53*01 | gp100 | YLEPGPVTA | HLA-A*02:01 | Orthogonality validation |
| PMEL | β | TRBV7-3*01 | TRBJ2-3*01 | gp100 | YLEPGPVTA | HLA-A*02:01 | Orthogonality validation |
| Tyrosinase | α | TRAV4*01 | TRAJ42*01 | tyrosinase | YMDGTMSQV | HLA-A*02:01 | Orthogonality validation (human/murine TCR-C) |
| Tyrosinase | β | TRBV10-3*01 | TRBJ1-3*01 | tyrosinase | YMDGTMSQV | HLA-A*02:01 | Orthogonality validation (human/murine TCR-C) |
| DMF4 | α | TRAV35*01 | TRAJ49*01 | MART-1 | AAGIGILTV | HLA-A*02:01 | Orthogonality validation (human/murine TCR-C) |
| DMF4 | β | TRBV10-3*01 | TRBJ1-5*01 | MART-1 | AAGIGILTV | HLA-A*02:01 | Orthogonality validation (human/murine TCR-C) |
| MEL5-α24β17 | α | TRAV12-2*01 | TRAJ27*01 | MART-1 | AAGIGILTV | HLA-A*02:01 | Orthogonality validation, TCR-IgG, TriTE pre-assay |
| MEL5-α24β17 | β | TRBV30*01 | TRBJ2-2*01 | MART-1 | AAGIGILTV | HLA-A*02:01 | Orthogonality validation, TCR-IgG, TriTE pre-assay |
| NYESO1 | α | TRAV3*01 | TRAJ28*01 | NY-ESO-1 | SLLMWITQC | HLA-A*02:01 | Orthogonality validation, TCE functional evaluation |
| NYESO1 | β | TRBV29-1*01 | TRBJ2-3*01 | NY-ESO-1 | SLLMWITQC | HLA-A*02:01 | Orthogonality validation, TCE functional evaluation |
| MAGEA4 | α | TRAV10*02 | TRAJ6*01 | MAGE-A4 | GVYDGREHTV | HLA-A*02:01 | Orthogonality validation, TCE functional evaluation |
| MAGEA4 | β | TRBV28*01 | TRBJ2-7*01 | MAGE-A4 | GVYDGREHTV | HLA-A*02:01 | Orthogonality validation, TCE functional evaluation |
| PRAME | α | TRAV26-2*01 | TRAJ28*01 | PRAME | SLLQHLIGL | HLA-A*02:01 | Orthogonality validation, TCE functional evaluation |
| PRAME | β | TRBV19*01 | TRBJ1-6*02 | PRAME | SLLQHLIGL | HLA-A*02:01 | Orthogonality validation, TCE functional evaluation |
| A6 | α | TRAV12-2*01 | TRAJ24*02 | Tax of HTLV-1 | LLFGYPVYV | HLA-A*02:01 | TCE functional evaluation (isotype control) |
| A6 | β | TRBV6-5*01 | TRBJ2-7*01 | Tax of HTLV-1 | LLFGYPVYV | HLA-A*02:01 | TCE functional evaluation (isotype control) |

Notes: Assigned V and J genes indicate the closest human germline framework lineage and do not imply full germline identity across CDR or J regions.

**Supplementary Table S9.** Expected molecular masses for α/β pairing states used in intact-mass spectrometry analysis (Fig. 3g).

| No | TCR type | Co-expressed TCR chains | WTα (Da) | WTβ (Da) | MTα (Da) | MTβ (Da) | WW (Da) | WM (Da) | MW (Da) | MM (Da) |
| --- | --- | --- | --- | --- | --- | --- | --- | --- | --- | --- |
| 1 | TCR-C | 8His-stCa/stCb | 11926 | 14892 | N/A | N/A | 26818 | N/A | N/A | N/A |
| 2 | TCR-C | 8His-combiC42a/combiC42b | N/A | N/A | 12045 | 14796 | N/A | N/A | N/A | 26841 |
| 3 | TCR-C | 8His-combiC26a/combiC26b | N/A | N/A | 12061 | 14726 | N/A | N/A | N/A | 26787 |
| 4 | TCR-C | 8His-stCa/stCb/combiC42b | 11926 | 14892 | N/A | 14796 | 26818 | 26722 | N/A | N/A |
| 5 | TCR-C | 8His-combiC42a/stCb/combiC42b | N/A | 14892 | 12045 | 14796 | N/A | N/A | 26938 | 26841 |
| 6 | TCR-C | 8His-stCa/stCb/combiC26b | 11926 | 14892 | N/A | 14726 | 26818 | 26652 | N/A | N/A |
| 7 | TCR-C | 8His-combiC26a/stCb/combiC26b | N/A | 14892 | 12061 | 14726 | N/A | N/A | 26954 | 26787 |
| 8 | TCR-C | 8His-stCa/stCb/8His-combiC26a/combiC26b | 11926 | 14892 | 12061 | 14726 | 26818 | 26652 | 26954 | 26787 |
| 9 | TCR-VC | 8His-wtVstCa/wtVstCb | 23964 | 27154 | N/A | N/A | 51119 | N/A | N/A | N/A |
| 10 | TCR-VC | 8His-desV30combiC42a/desV30combiC42b | N/A | N/A | 24084 | 27093 | N/A | N/A | N/A | 51177 |
| 11 | TCR-VC | 8His-desV30combiC26a/desV30combiC26b | N/A | N/A | 24100 | 27023 | N/A | N/A | N/A | 51123 |
| 12 | TCR-VC | 8His-wtVstCa/wtVstCb/desV30combiC42b | 23964 | 27154 | N/A | 27093 | 51119 | 51057 | N/A | N/A |
| 13 | TCR-VC | 8His-desV30combiC42a/wtVstCb/desV30combiC42b | N/A | 27154 | 24084 | 27093 | N/A | N/A | 51238 | 51177 |
| 14 | TCR-VC | 8His-wtVstCa/wtVstCb/desV30combiC26b | 23964 | 27154 | N/A | 27023 | 51119 | 50987 | N/A | N/A |
| 15 | TCR-VC | 8His-desV30combiC26a/wtVstCb/desV30combiC26b | N/A | 27154 | 24100 | 27023 | N/A | N/A | 51254 | 51123 |
| 16 | TCR-VC | 8His-wtVstCa/wtVstCb/8His-desV30combiC42a/desV30combiC42b | 23964 | 27154 | 24084 | 27093 | 51119 | 51057 | 51238 | 51177 |

TCR-C: TCR constant domain.
TCR-V: TCR variable domain.
TCR-VC: TCR construct containing both variable and constant domains.
stC: Stabilized TCR constant domain containing additional stabilizing mutations.
combiC: Combinatorial orthogonal TCR constant-domain design.
desV: Designed TCR variable-domain interface.
wtVstC: Wild-type TCR variable domain fused to a stabilized TCR constant domain.
WT: Wild type.
MT: Mutant (orthogonally designed).
MM: Mutant $\alpha$ chain paired with mutant $\beta$ chain (on-target pairing).
MW: Mutant $\alpha$ chain paired with wild-type $\beta$ chain.
WM: Wild-type $\alpha$ chain paired with mutant $\beta$ chain.

**Supplementary Table S10.** Structure information.

|  |  |
| --- | --- |
| **Data reduction** |  |
| Space group | P61 |
| *Cell dimensions* |  |
| a, b, c (Å) | 53.458, 53.458, 163.788 |
| $\alpha$, $\beta$, $\gamma$ (°) | 90, 90, 120 |
| Resolution (Å)* | 23.15 - 2.05 (2.12-2.05) |
| Rmerge* | 12.1 (116.1) |
| Rmeas* | 12.4 (120.2) |
| CC½ (%)* | 99.9 (79.4) |
| I / σI* | 19.5 (2.0) |
| Completeness (%)* | 99.98 (100) |
| Total reflections | 404171 (24637) |
| Unique reflections* | 16557 (1650) |
| Redundancy* | 24.4 (14.9) |
| VM (Å³/Da) | 2.52 |
| **Refinement** |  |
| Resolution (Å) | 23.15-2.05 (2.08-2.05) |
| No. reflections | 32741 (1253) |
| Rwork / Rfree | 19.61 / 24.77 (29.27 / 32.66) |
| *No. atoms* |  |
| Protein | 1529 |
| Water | 105 |
| *Mean B-factors* |  |
| Overall (Å²) | 39.89 |
| Protein (Å²) | 40.03 |
| Water (Å²) | 37.80 |
| *R.m.s. deviations* |  |
| Bond lengths (Å) | 0.007 |
| Bond angles (°) | 0.849 |
| *Ramachandran* |  |
| Favored (%) | 88.3 |
| Allowed (%) | 9.4 |
| Outliers (%) | 2.3 |
| Rotamer outliers (%) | 2.3 |
| Clashscore | 2.34 |
| Wilson B-factor (Å²) | 30.29 |

\* Figures for the highest resolution shell are shown in parentheses.

**Supplementary Table S11.** IMGT, Kabat and PDB numbering for TCR-Cα.

| IMGT DOMAIN | IMGT | Kabat | PDB (6U07) | Residue [1 letter] | Residue [3 letter] |
| --- | --- | --- | --- | --- | --- |
| V-ALPHA [D1] | 128V | - | 118 | P | PRO |
| C-ALPHA [D1] | 1.5C | 117 | 119 | Y | TYR |
| C-ALPHA [D1] | 1.4C | 118 | 120 | I | ILE |
| C-ALPHA [D1] | 1.3C | 119 | 121 | Q | GLN |
| C-ALPHA [D1] | 1.2C | 120 | 122 | N | ASN |
| C-ALPHA [D1] | 1.1C | 121 | 123 | P | PRO |
| C-ALPHA [D1] | 1C | 122 | 124 | D | ASP |
| C-ALPHA [D1] | 2C | 123 | 125 | P | PRO |
| C-ALPHA [D1] | 3C | 124 | 126 | A | ALA |
| C-ALPHA [D1] | 4C | 125 | 127 | V | VAL |
| C-ALPHA [D1] | 5C | 126 | 128 | Y | TYR |
| C-ALPHA [D1] | 6C | 127 | 129 | Q | GLN |
| C-ALPHA [D1] | 7C | 128 | 130 | L | LEU |
| C-ALPHA [D1] | 8C | 129 | 131 | R | ARG |
| C-ALPHA [D1] | 9C | 132 | 132 | D | ASP |
| C-ALPHA [D1] | 10C | 133 | 133 | S | SER |
| C-ALPHA [D1] | 11C | 134 | 134 | K | LYS |
| C-ALPHA [D1] | 17C | 135 | 135 | S | SER |
| C-ALPHA [D1] | 18C | 136 | 136 | S | SER |
| C-ALPHA [D1] | 19C | 137 | 137 | D | ASP |
| C-ALPHA [D1] | 20C | 138 | 138 | K | LYS |
| C-ALPHA [D1] | 21C | 139 | 139 | F | PHE |
| C-ALPHA [D1] | 22C | 140 | 140 | V | VAL |
| C-ALPHA [D1] | 23C | 141 | 141 | C | CYS |
| C-ALPHA [D1] | 24C | 142 | 142 | L | LEU |
| C-ALPHA [D1] | 25C | 143 | 143 | F | PHE |
| C-ALPHA [D1] | 26C | 144 | 144 | T | THR |
| C-ALPHA [D1] | 27C | 145 | 145 | D | ASP |
| C-ALPHA [D1] | 28C | 146 | 146 | F | PHE |
| C-ALPHA [D1] | 29C | 147 | 147 | D | ASP |
| C-ALPHA [D1] | 30C | 148 | 148 | S | SER |
| C-ALPHA [D1] | 36C | 149 | 149 | Q | GLN |
| C-ALPHA [D1] | 37C | 150 | 150 | I | ILE |
| C-ALPHA [D1] | 38C | 151 | 151 | N | ASN |
| C-ALPHA [D1] | 39C | 152 | 152 | V | VAL |
| C-ALPHA [D1] | 40C | 153 | 153 | S | SER |
| C-ALPHA [D1] | 41C | 154 | 154 | Q | GLN |
| C-ALPHA [D1] | 42C | 155 | 155 | S | SER |
| C-ALPHA [D1] | 43C | 156 | 156 | K | LYS |
| C-ALPHA [D1] | 44C | 157 | 157 | D | ASP |
| C-ALPHA [D1] | 45C | 158 | 158 | S | SER |
| C-ALPHA [D1] | 77C | 159 | 159 | D | ASP |
| C-ALPHA [D1] | 78C | 160 | 160 | V | VAL |
| C-ALPHA [D1] | 79C | 161 | 161 | Y | TYR |
| C-ALPHA [D1] | 80C | 162 | 162 | I | ILE |
| C-ALPHA [D1] | 81C | 163 | 163 | T | THR |
| C-ALPHA [D1] | 82C | 164 | 164 | D | ASP |
| C-ALPHA [D1] | 83C | 165 | 165 | K | LYS |

**Supplementary Table S11. IMGT, Kabat and PDB numbering for TCR- $\alpha$ .**
| IMGT DOMAIN | IMGT | Kabat | PDB (6U07) | Residue [1 letter] | Residue [3 letter] |
| --- | --- | --- | --- | --- | --- |
| C-ALPHA [D1] | 84C | 166 | 166 | C | CYS |
| C-ALPHA [D1] | 84.1C | 167 | 167 | V | VAL |
| C-ALPHA [D1] | 84.2C | 168 | 168 | L | LEU |
| C-ALPHA [D1] | 84.3C | 169 | 169 | D | ASP |
| C-ALPHA [D1] | 84.4C | 170 | 170 | M | MET |
| C-ALPHA [D1] | 84.5C | 171 | 171 | R | ARG |
| C-ALPHA [D1] | 84.6C | 172 | 172 | S | SER |
| C-ALPHA [D1] | 84.7C | 173 | 173 | M | MET |
| C-ALPHA [D1] | 85.6C | 174 | 174 | D | ASP |
| C-ALPHA [D1] | 85.5C | 175 | 175 | F | PHE |
| C-ALPHA [D1] | 85.4C | 176 | 176 | K | LYS |
| C-ALPHA [D1] | 85.3C | 177 | 177 | S | SER |
| C-ALPHA [D1] | 85.2C | 178 | 178 | N | ASN |
| C-ALPHA [D1] | 85.1C | 179 | 179 | S | SER |
| C-ALPHA [D1] | 85C | 180 | 180 | A | ALA |
| C-ALPHA [D1] | 86C | 181 | 181 | V | VAL |
| C-ALPHA [D1] | 87C | 182 | 182 | A | ALA |
| C-ALPHA [D1] | 88C | 183 | 183 | W | TRP |
| C-ALPHA [D1] | 89C | 184 | 184 | S | SER |
| C-ALPHA [D1] | 90C | 185 | 185 | N | ASN |
| C-ALPHA [D1] | 91C | 186 | 186 | K | LYS |
| C-ALPHA [D1] | 92C | 187 | 187 | S | SER |
| C-ALPHA [D1] | 101C | 188 | 188 | D | ASP |
| C-ALPHA [D1] | 102C | 189 | 189 | F | PHE |
| C-ALPHA [D1] | 103C | 190 | 190 | T | THR |
| C-ALPHA [D1] | 104C | 191 | 191 | C | CYS |
| C-ALPHA [D1] | 105C | 192 | 192 | A | ALA |
| C-ALPHA [D1] | 106C | 193 | 193 | N | ASN |
| C-ALPHA [D1] | 107C | 194 | 194 | A | ALA |
| C-ALPHA [D1] | 108C | 195 | 195 | F | PHE |
| C-ALPHA [D1] | 109C | 196 | 196 | N | ASN |
| C-ALPHA [D1] | 110C | 197 | 197 | N | ASN |
| C-ALPHA [D1] | 113C | 198 | 198 | S | SER |
| C-ALPHA [D1] | 114C | 199 | 199 | I | ILE |
| C-ALPHA [D1] | 115C | 200 | 200 | I | ILE |
| C-ALPHA [D1] | 116C | 201 | 201 | P | PRO |
| C-ALPHA [D1] | 117C | 202 | 202 | E | GLU |
| C-ALPHA [D1] | 118C | 203 | 203 | D | ASP |
| C-ALPHA [D1] | 119C | 204 | 204 | T | THR |
| C-ALPHA [D1] | 120C | 205 | 205 | F | PHE |
| C-ALPHA [D1] | 121C | 206 | 206 | F | PHE |
| C-ALPHA [D1] | 122C | 207 | 207 | P | PRO |
| C-ALPHA [D1] | 123C | 208 | 208 | S | SER |
| C-ALPHA [D1] | 124C | 209 | 209 | P | PRO |
| C-ALPHA [D1] | 125C | 210 | 210 | E | GLU |
| C-ALPHA [D1] | 126C | 211 | 211 | S | SER |
| C-ALPHA [D1] | 127C | 212 | 212 | S | SER |
| C-ALPHA [D1] | 128C | 213 | 213 | C | CYS |

**Supplementary Table S12.** IMGT, Kabat and PDB numbering for TCR-Cβ.

| IMGT DOMAIN | IMGT | Kabat | PDB (6U07) | Residue [1 letter] | Residue [3 letter] |
| --- | --- | --- | --- | --- | --- |
| C-BETA-2 [D1] | 1.7C | 117 | 117 | E | GLU |
| C-BETA-2 [D1] | 1.6C | 118 | 118 | D | ASP |
| C-BETA-2 [D1] | 1.5C | 119 | 119 | L | LEU |
| C-BETA-2 [D1] | 1.4C | 120 | 120 | K | LYS |
| C-BETA-2 [D1] | 1.3C | 121 | 121 | N | ASN |
| C-BETA-2 [D1] | 1.2C | 122 | 122 | V | VAL |
| C-BETA-2 [D1] | 1.1C | 123 | 123 | F | PHE |
| C-BETA-2 [D1] | 1C | 124 | 124 | P | PRO |
| C-BETA-2 [D1] | 2C | 125 | 125 | P | PRO |
| C-BETA-2 [D1] | 3C | 126 | 126 | E | GLU |
| C-BETA-2 [D1] | 4C | 127 | 127 | V | VAL |
| C-BETA-2 [D1] | 5C | 128 | 128 | A | ALA |
| C-BETA-2 [D1] | 6C | 129 | 129 | V | VAL |
| C-BETA-2 [D1] | 7C | 130 | 130 | F | PHE |
| C-BETA-2 [D1] | 8C | 131 | 131 | E | GLU |
| C-BETA-2 [D1] | 9C | 132 | 132 | P | PRO |
| C-BETA-2 [D1] | 10C | 133 | 133 | S | SER |
| C-BETA-2 [D1] | 11C | 134 | 134 | K | LYS |
| C-BETA-2 [D1] | 12C | 135 | 135 | A | ALA |
| C-BETA-2 [D1] | 13C | 136 | 136 | E | GLU |
| C-BETA-2 [D1] | 14C | 137 | 137 | I | ILE |
| C-BETA-2 [D1] | 15C | 138 | 138 | S | SER |
| C-BETA-2 [D1] | 15.1C | 139 | 139 | R | ARG |
| C-BETA-2 [D1] | 16C | 140 | 140 | T | THR |
| C-BETA-2 [D1] | 17C | 141 | 141 | Q | GLN |
| C-BETA-2 [D1] | 18C | 142 | 142 | K | LYS |
| C-BETA-2 [D1] | 19C | 143 | 143 | A | ALA |
| C-BETA-2 [D1] | 20C | 144 | 144 | T | THR |
| C-BETA-2 [D1] | 21C | 145 | 145 | L | LEU |
| C-BETA-2 [D1] | 22C | 146 | 146 | V | VAL |
| C-BETA-2 [D1] | 23C | 147 | 147 | C | CYS |
| C-BETA-2 [D1] | 24C | 148 | 148 | L | LEU |
| C-BETA-2 [D1] | 25C | 149 | 149 | A | ALA |
| C-BETA-2 [D1] | 26C | 150 | 150 | T | THR |
| C-BETA-2 [D1] | 27C | 151 | 151 | G | GLY |
| C-BETA-2 [D1] | 28C | 152 | 152 | F | PHE |
| C-BETA-2 [D1] | 29C | 153 | 153 | Y | TYR |
| C-BETA-2 [D1] | 30C | 154 | 154 | P | PRO |
| C-BETA-2 [D1] | 35C | 155 | 155 | P | PRO |
| C-BETA-2 [D1] | 36C | 156 | 156 | H | HIS |
| C-BETA-2 [D1] | 37C | 157 | 157 | V | VAL |
| C-BETA-2 [D1] | 38C | 158 | 158 | E | GLU |
| C-BETA-2 [D1] | 39C | 159 | 159 | L | LEU |
| C-BETA-2 [D1] | 40C | 160 | 160 | S | SER |
| C-BETA-2 [D1] | 41C | 161 | 161 | W | TRP |
| C-BETA-2 [D1] | 42C | 162 | 162 | W | TRP |
| C-BETA-2 [D1] | 43C | 163 | 163 | V | VAL |
| C-BETA-2 [D1] | 44C | 164 | 164 | N | ASN |
| C-BETA-2 [D1] | 45C | 165 | 165 | G | GLY |

**Supplementary Table S12. IMGT, Kabat and PDB numbering for TCR-Cβ .**
| IMGT DOMAIN | IMGT | Kabat | PDB (6U07) | Residue [1 letter] | Residue [3 letter] |
| --- | --- | --- | --- | --- | --- |
| C-BETA-2 [D1] | 45.1C | 166 | 166 | K | LYS |
| C-BETA-2 [D1] | 45.2C | 167 | 167 | E | GLU |
| C-BETA-2 [D1] | 45.3C | 168 | 168 | V | VAL |
| C-BETA-2 [D1] | 45.4C | 169 | 169 | H | HIS |
| C-BETA-2 [D1] | 45.5C | 170 | 170 | D | ASP |
| C-BETA-2 [D1] | 77C | 171 | 171 | G | GLY |
| C-BETA-2 [D1] | 78C | 172 | 172 | V | VAL |
| C-BETA-2 [D1] | 79C | 173 | 173 | C | CYS |
| C-BETA-2 [D1] | 80C | 174 | 174 | T | THR |
| C-BETA-2 [D1] | 81C | 175 | 175 | D | ASP |
| C-BETA-2 [D1] | 82C | 176 | 176 | P | PRO |
| C-BETA-2 [D1] | 83C | 177 | 177 | Q | GLN |
| C-BETA-2 [D1] | 84C | 178 | 178 | P | PRO |
| C-BETA-2 [D1] | 84.1C | 179 | 179 | L | LEU |
| C-BETA-2 [D1] | 84.2C | 180 | 180 | K | LYS |
| C-BETA-2 [D1] | 84.3C | 181 | 181 | E | GLU |
| C-BETA-2 [D1] | 84.4C | 182 | 182 | Q | GLN |
| C-BETA-2 [D1] | 84.5C | 183 | 183 | P | PRO |
| C-BETA-2 [D1] | 84.6C | 184 | 184 | A | ALA |
| C-BETA-2 [D1] | 84.7C | 185 | 185 | L | LEU |
| C-BETA-2 [D1] | 85.6C | 186 | 186 | N | ASN |
| C-BETA-2 [D1] | 85.5C | 187 | 187 | D | ASP |
| C-BETA-2 [D1] | 85.4C | 188 | 188 | S | SER |
| C-BETA-2 [D1] | 85.3C | 189 | 189 | R | ARG |
| C-BETA-2 [D1] | 85.2C | 190 | 190 | Y | TYR |
| C-BETA-2 [D1] | 85.1C | 191 | 191 | A | ALA |
| C-BETA-2 [D1] | 85C | 192 | 192 | L | LEU |
| C-BETA-2 [D1] | 86C | 193 | 193 | S | SER |
| C-BETA-2 [D1] | 87C | 194 | 194 | S | SER |
| C-BETA-2 [D1] | 88C | 195 | 195 | R | ARG |
| C-BETA-2 [D1] | 89C | 196 | 196 | L | LEU |
| C-BETA-2 [D1] | 90C | 197 | 197 | R | ARG |
| C-BETA-2 [D1] | 91C | 198 | 198 | V | VAL |
| C-BETA-2 [D1] | 92C | 199 | 199 | S | SER |
| C-BETA-2 [D1] | 93C | 200 | 200 | A | ALA |
| C-BETA-2 [D1] | 94C | 201 | 201 | T | THR |
| C-BETA-2 [D1] | 95C | 202 | 202 | F | PHE |
| C-BETA-2 [D1] | 96C | 203 | 203 | W | TRP |
| C-BETA-2 [D1] | 96.1C | 204 | 204 | Q | GLN |
| C-BETA-2 [D1] | 97C | 205 | 205 | D | ASP |
| C-BETA-2 [D1] | 98C | 206 | 206 | P | PRO |
| C-BETA-2 [D1] | 99C | 207 | 207 | R | ARG |
| C-BETA-2 [D1] | 100C | 208 | 208 | N | ASN |
| C-BETA-2 [D1] | 101C | 209 | 209 | H | HIS |
| C-BETA-2 [D1] | 102C | 210 | 210 | F | PHE |
| C-BETA-2 [D1] | 103C | 211 | 211 | R | ARG |
| C-BETA-2 [D1] | 104C | 212 | 212 | C | CYS |
| C-BETA-2 [D1] | 105C | 213 | 213 | Q | GLN |
| C-BETA-2 [D1] | 106C | 214 | 214 | V | VAL |

**Supplementary Table S12. IMGT, Kabat and PDB numbering for TCR-Cβ .**
| IMGT DOMAIN | IMGT | Kabat | PDB (6U07) | Residue [1 letter] | Residue [3 letter] |
| --- | --- | --- | --- | --- | --- |
| C-BETA-2 [D1] | 107C | 215 | 215 | Q | GLN |
| C-BETA-2 [D1] | 108C | 216 | 216 | F | PHE |
| C-BETA-2 [D1] | 109C | 217 | 217 | Y | TYR |
| C-BETA-2 [D1] | 110C | 218 | 218 | G | GLY |
| C-BETA-2 [D1] | 111C | 219 | 219 | L | LEU |
| C-BETA-2 [D1] | 111.1C | 220 | 220 | S | SER |
| C-BETA-2 [D1] | 111.2C | 221 | 221 | E | GLU |
| C-BETA-2 [D1] | 111.3C | 222 | 222 | N | ASN |
| C-BETA-2 [D1] | 111.4C | 223 | 223 | D | ASP |
| C-BETA-2 [D1] | 111.5C | 224 | 224 | E | GLU |
| C-BETA-2 [D1] | 111.6C | 225 | 225 | W | TRP |
| C-BETA-2 [D1] | 112.6C | 226 | 226 | T | THR |
| C-BETA-2 [D1] | 112.5C | 227 | 227 | Q | GLN |
| C-BETA-2 [D1] | 112.4C | 228 | 228 | D | ASP |
| C-BETA-2 [D1] | 112.3C | 229 | 229 | R | ARG |
| C-BETA-2 [D1] | 112.2C | 230 | 230 | A | ALA |
| C-BETA-2 [D1] | 112.1C | 231 | 231 | K | LYS |
| C-BETA-2 [D1] | 112C | 232 | 232 | P | PRO |
| C-BETA-2 [D1] | 113C | 233 | 233 | V | VAL |
| C-BETA-2 [D1] | 114C | 234 | 234 | T | THR |
| C-BETA-2 [D1] | 115C | 235 | 235 | Q | GLN |
| C-BETA-2 [D1] | 116C | 236 | 236 | I | ILE |
| C-BETA-2 [D1] | 117C | 237 | 237 | V | VAL |
| C-BETA-2 [D1] | 118C | 238 | 238 | S | SER |
| C-BETA-2 [D1] | 119C | 239 | 239 | A | ALA |
| C-BETA-2 [D1] | 120C | 240 | 240 | E | GLU |
| C-BETA-2 [D1] | 121C | 241 | 241 | A | ALA |
| C-BETA-2 [D1] | 122C | 242 | 242 | W | TRP |
| C-BETA-2 [D1] | 123C | 243 | 243 | G | GLY |
| C-BETA-2 [D1] | 124C | 244 | 244 | R | ARG |
| C-BETA-2 [D1] | 125C | 245 | 245 | A | ALA |
| C-BETA-2 [D1] | 126C | 246 | 246 | D | ASP |
| C-BETA-2 [D1] | 127C | 247 | 247 | C | CYS |

**Supplementary Table S13.** IMGT, Kabat and PDB numbering for TCR-Vα.

| IMGT DOMAIN | FR/CDR | IMGT | Kabat | PDB (2F53) | Residue [1 letter] | Residue [3 letter] |
| --- | --- | --- | --- | --- | --- | --- |
| - | - | - | - | -1 | M | MET |
| V-ALPHA [D1] | FR1 | 1V | 1 | 0 | K | LYS |
| V-ALPHA [D1] | FR1 | 2V | 2 | 1 | Q | GLN |
| V-ALPHA [D1] | FR1 | 3V | 3 | 2 | E | GLU |
| V-ALPHA [D1] | FR1 | 4V | 4 | 3 | V | VAL |
| V-ALPHA [D1] | FR1 | 5V | 5 | 4 | T | THR |
| V-ALPHA [D1] | FR1 | 6V | 6 | 5 | Q | GLN |
| V-ALPHA [D1] | FR1 | 7V | 7 | 6 | I | ILE |
| V-ALPHA [D1] | FR1 | 8V | 8 | 7 | P | PRO |
| V-ALPHA [D1] | FR1 | 9V | 9 | 8 | A | ALA |
| V-ALPHA [D1] | FR1 | 10V | 10 | 9 | A | ALA |
| V-ALPHA [D1] | FR1 | 11V | 11 | 10 | L | LEU |
| V-ALPHA [D1] | FR1 | 12V | 12 | 11 | S | SER |
| V-ALPHA [D1] | FR1 | 13V | 13 | 12 | V | VAL |
| V-ALPHA [D1] | FR1 | 14V | 14 | 13 | P | PRO |
| V-ALPHA [D1] | FR1 | 15V | 15 | 14 | E | GLU |
| V-ALPHA [D1] | FR1 | 16V | 16 | 15 | G | GLY |
| V-ALPHA [D1] | FR1 | 17V | 17 | 16 | E | GLU |
| V-ALPHA [D1] | FR1 | 18V | 18 | 17 | N | ASN |
| V-ALPHA [D1] | FR1 | 19V | 19 | 18 | L | LEU |
| V-ALPHA [D1] | FR1 | 20V | 20 | 19 | V | VAL |
| V-ALPHA [D1] | FR1 | 21V | 21 | 20 | L | LEU |
| V-ALPHA [D1] | FR1 | 22V | 22 | 21 | N | ASN |
| V-ALPHA [D1] | FR1 | 23V | 23 | 22 | C | CYS |
| V-ALPHA [D1] | FR1 | 24V | 24 | 23 | S | SER |
| V-ALPHA [D1] | FR1 | 25V | 25 | 24 | F | PHE |
| V-ALPHA [D1] | FR1 | 26V | 26 | 25 | T | THR |
| V-ALPHA [D1] | CDR1 | 27V | 27 | 26 | D | ASP |
| V-ALPHA [D1] | CDR1 | 28V | 29 | 27 | S | SER |
| V-ALPHA [D1] | CDR1 | 29V | 30 | 28 | A | ALA |
| - | - | 30 | - | - | - | - |
| - | - | 31 | - | - | - | - |
| - | - | 32 | - | - | - | - |
| - | - | 33 | - | - | - | - |
| - | - | 34 | - | - | - | - |
| - | - | 35 | - | - | - | - |
| V-ALPHA [D1] | CDR1 | 36V | 30a | 29 | I | ILE |
| V-ALPHA [D1] | CDR1 | 37V | 30b | 30 | Y | TYR |
| V-ALPHA [D1] | CDR1 | 38V | 31 | 31 | N | ASN |
| V-ALPHA [D1] | FR2 | 39V | 32 | 32 | L | LEU |
| V-ALPHA [D1] | FR2 | 40V | 33 | 33 | Q | GLN |
| V-ALPHA [D1] | FR2 | 41V | 34 | 34 | W | TRP |
| V-ALPHA [D1] | FR2 | 42V | 35 | 35 | F | PHE |
| V-ALPHA [D1] | FR2 | 43V | 36 | 36 | R | ARG |
| V-ALPHA [D1] | FR2 | 44V | 37 | 37 | Q | GLN |
| V-ALPHA [D1] | FR2 | 45V | 38 | 38 | D | ASP |
| V-ALPHA [D1] | FR2 | 46V | 39 | 39 | P | PRO |
| V-ALPHA [D1] | FR2 | 47V | 40 | 40 | G | GLY |

**Supplementary Table S13. IMGT, Kabat and PDB numbering for TCR-V $\alpha$ .**
| IMGT DOMAIN | FR/CDR | IMGT | Kabat | PDB (2F53) | Residue [1 letter] | Residue [3 letter] |
| --- | --- | --- | --- | --- | --- | --- |
| V-ALPHA [D1] | FR2 | 48V | 41 | 41 | K | LYS |
| V-ALPHA [D1] | FR2 | 49V | 42 | 42 | G | GLY |
| V-ALPHA [D1] | FR2 | 50V | 43 | 43 | L | LEU |
| V-ALPHA [D1] | FR2 | 51V | 44 | 44 | T | THR |
| V-ALPHA [D1] | FR2 | 52V | 45 | 45 | S | SER |
| V-ALPHA [D1] | FR2 | 53V | 46 | 46 | L | LEU |
| V-ALPHA [D1] | FR2 | 54V | 47 | 47 | L | LEU |
| V-ALPHA [D1] | FR2 | 55V | 48 | 48 | L | LEU |
| V-ALPHA [D1] | CDR2 | 56V | 49 | 49 | I | ILE |
| V-ALPHA [D1] | CDR2 | 57V | 50 | 50 | P | PRO |
| V-ALPHA [D1] | CDR2 | 58V | - | 51 | F | PHE |
| V-ALPHA [D1] | CDR2 | 59V | - | 52 | W | TRP |
| - | - | 60 | - | - | - | - |
| - | - | 61 | - | - | - | - |
| - | - | 62 | - | - | - | - |
| V-ALPHA [D1] | CDR2 | 63V | 55 | 53 | Q | GLN |
| V-ALPHA [D1] | CDR2 | 64V | 56 | 54 | R | ARG |
| V-ALPHA [D1] | CDR2 | 65V | 57 | 55 | E | GLU |
| V-ALPHA [D1] | FR3 | 66V | 58 | 56 | Q | GLN |
| V-ALPHA [D1] | FR3 | 67V | 59 | 57 | T | THR |
| V-ALPHA [D1] | FR3 | 68V | - | 58 | S | SER |
| - | - | 69 | - | - | - | - |
| - | - | 70 | - | - | - | - |
| - | - | 71 | - | - | - | - |
| - | - | 72 | - | - | - | - |
| - | - | 73 | - | - | - | - |
| V-ALPHA [D1] | FR3 | 74V | 60 | 59 | G | GLY |
| V-ALPHA [D1] | FR3 | 75V | 61 | 60 | R | ARG |
| V-ALPHA [D1] | FR3 | 76V | 62 | 61 | L | LEU |
| V-ALPHA [D1] | FR3 | 77V | 63 | 62 | N | ASN |
| V-ALPHA [D1] | FR3 | 78V | 64 | 63 | A | ALA |
| V-ALPHA [D1] | FR3 | 79V | 65 | 64 | S | SER |
| V-ALPHA [D1] | FR3 | 80V | 66 | 65 | L | LEU |
| V-ALPHA [D1] | FR3 | 81V | 67 | 66 | D | ASP |
| V-ALPHA [D1] | FR3 | 82V | 68 | 67 | K | LYS |
| V-ALPHA [D1] | FR3 | 83V | 69 | 68 | S | SER |
| V-ALPHA [D1] | FR3 | 84V | 70 | 69 | S | SER |
| V-ALPHA [D1] | FR3 | 85V | 71 | 70 | G | GLY |
| V-ALPHA [D1] | FR3 | 86V | 72 | 71 | R | ARG |
| V-ALPHA [D1] | FR3 | 87V | 73 | 72 | S | SER |
| V-ALPHA [D1] | FR3 | 88V | 74 | 73 | T | THR |
| V-ALPHA [D1] | FR3 | 89V | 75 | 74 | L | LEU |
| V-ALPHA [D1] | FR3 | 90V | 76 | 75 | Y | TYR |
| V-ALPHA [D1] | FR3 | 91V | 77 | 76 | I | ILE |
| V-ALPHA [D1] | FR3 | 92V | 78 | 77 | A | ALA |
| V-ALPHA [D1] | FR3 | 93V | 79 | 78 | A | ALA |
| V-ALPHA [D1] | FR3 | 94V | 80 | 79 | S | SER |
| V-ALPHA [D1] | FR3 | 95V | 81 | 80 | Q | GLN |

**Supplementary Table S13. IMGT, Kabat and PDB numbering for TCR-V $\alpha$ .**
| IMGT DOMAIN | FR/CDR | IMGT | Kabat | PDB (2F53) | Residue [1 letter] | Residue [3 letter] |
| --- | --- | --- | --- | --- | --- | --- |
| V-ALPHA [D1] | FR3 | 96V | 82 | 81 | P | PRO |
| V-ALPHA [D1] | FR3 | 97V | 83 | 82 | G | GLY |
| V-ALPHA [D1] | FR3 | 98V | 84 | 83 | D | ASP |
| V-ALPHA [D1] | FR3 | 99V | 85 | 84 | S | SER |
| V-ALPHA [D1] | FR3 | 100V | 86 | 85 | A | ALA |
| V-ALPHA [D1] | FR3 | 101V | 87 | 86 | T | THR |
| V-ALPHA [D1] | FR3 | 102V | 88 | 87 | Y | TYR |
| V-ALPHA [D1] | FR3 | 103V | 89 | 88 | L | LEU |
| V-ALPHA [D1] | FR3 | 104V | 90 | 89 | C | CYS |
| V-ALPHA [D1] | CDR3 | 105V | 91 | 90 | A | ALA |
| V-ALPHA [D1] | CDR3 | 106V | 92 | 91 | V | VAL |
| V-ALPHA [D1] | CDR3 | 107V | 93 | 92 | R | ARG |
| V-ALPHA [D1] | CDR3 | 108V | - | 93 | P | PRO |
| V-ALPHA [D1] | CDR3 | 109V | - | 94 | T | THR |
| V-ALPHA [D1] | CDR3 | 110V | - | 95 | S | SER |
| V-ALPHA [D1] | CDR3 | 111V | - | 96 | G | GLY |
| V-ALPHA [D1] | CDR3 | 112V | - | 97 | G | GLY |
| V-ALPHA [D1] | CDR3 | 113V | - | 98 | S | SER |
| V-ALPHA [D1] | CDR3 | 114V | - | 99 | Y | TYR |
| V-ALPHA [D1] | CDR3 | 115V | - | 100 | I | ILE |
| V-ALPHA [D1] | CDR3 | 116V | - | 101 | P | PRO |
| V-ALPHA [D1] | CDR3 | 117V | - | 102 | T | THR |
| V-ALPHA [D1] | FR4 | 118V | - | 103 | F | PHE |
| V-ALPHA [D1] | FR4 | 119V | - | 104 | G | GLY |
| V-ALPHA [D1] | FR4 | 120V | - | 105 | R | ARG |
| V-ALPHA [D1] | FR4 | 121V | - | 106 | G | GLY |
| V-ALPHA [D1] | FR4 | 122V | - | 107 | T | THR |
| V-ALPHA [D1] | FR4 | 123V | - | 108 | S | SER |
| V-ALPHA [D1] | FR4 | 124V | - | 109 | L | LEU |
| V-ALPHA [D1] | FR4 | 125V | - | 110 | I | ILE |
| V-ALPHA [D1] | FR4 | 126V | - | 111 | V | VAL |
| V-ALPHA [D1] | FR4 | 127V | - | 112 | H | HIS |
| V-ALPHA [D1] | FR4 | 128V | - | 113 | P | PRO |

**Supplementary Table S14.** IMGT, Kabat and PDB numbering for TCR-Vβ.

| IMGT DOMAIN | IMGT CDR | IMGT | Kabat | PDB (2F53) | Residue [1 letter] | Residue [3 letter] |
| --- | --- | --- | --- | --- | --- | --- |
| V-BETA [D1] | FR1 | 1V | 1 | -1 | N | ASN |
| V-BETA [D1] | FR1 | 2V | 2 | 0 | A | ALA |
| V-BETA [D1] | FR1 | 3V | 3 | 1 | G | GLY |
| V-BETA [D1] | FR1 | 4V | 4 | 2 | V | VAL |
| V-BETA [D1] | FR1 | 5V | 5 | 3 | T | THR |
| V-BETA [D1] | FR1 | 6V | 6 | 4 | Q | GLN |
| V-BETA [D1] | FR1 | 7V | 7 | 5 | T | THR |
| V-BETA [D1] | FR1 | 8V | 8 | 6 | P | PRO |
| V-BETA [D1] | FR1 | 9V | 9 | 7 | K | LYS |
| V-BETA [D1] | FR1 | 10V | 10 | 8 | F | PHE |
| V-BETA [D1] | FR1 | 11V | 11 | 9 | Q | GLN |
| V-BETA [D1] | FR1 | 12V | 12 | 10 | V | VAL |
| V-BETA [D1] | FR1 | 13V | 13 | 11 | L | LEU |
| V-BETA [D1] | FR1 | 14V | 14 | 12 | K | LYS |
| V-BETA [D1] | FR1 | 15V | 15 | 13 | T | THR |
| V-BETA [D1] | FR1 | 16V | 16 | 14 | G | GLY |
| V-BETA [D1] | FR1 | 17V | 17 | 15 | Q | GLN |
| V-BETA [D1] | FR1 | 18V | 18 | 16 | S | SER |
| V-BETA [D1] | FR1 | 19V | 19 | 17 | M | MET |
| V-BETA [D1] | FR1 | 20V | 20 | 18 | T | THR |
| V-BETA [D1] | FR1 | 21V | 21 | 19 | L | LEU |
| V-BETA [D1] | FR1 | 22V | 22 | 20 | Q | GLN |
| V-BETA [D1] | FR1 | 23V | 23 | 21 | C | CYS |
| V-BETA [D1] | FR1 | 24V | 24 | 22 | A | ALA |
| V-BETA [D1] | FR1 | 25V | 25 | 23 | Q | GLN |
| V-BETA [D1] | FR1 | 26V | 26 | 24 | D | ASP |
| V-BETA [D1] | CDR1 | 27V | 27 | 25 | M | MET |
| V-BETA [D1] | CDR1 | 28V | 28 | 26 | N | ASN |
| V-BETA [D1] | CDR1 | 29V | 29 | 27 | H | HIS |
| - | CDR1 | 30 | - | - | - | - |
| - | CDR1 | 31 | - | - | - | - |
| - | CDR1 | 32 | - | - | - | - |
| - | CDR1 | 33 | - | - | - | - |
| - | CDR1 | 34 | - | - | - | - |
| - | CDR1 | 35 | - | - | - | - |
| - | CDR1 | 36 | - | - | - | - |
| V-BETA [D1] | CDR1 | 37V | 30 | 28 | E | GLU |
| V-BETA [D1] | CDR1 | 38V | 31 | 29 | Y | TYR |
| V-BETA [D1] | FR2 | 39V | 32 | 30 | M | MET |
| V-BETA [D1] | FR2 | 40V | 33 | 31 | S | SER |
| V-BETA [D1] | FR2 | 41V | 34 | 32 | W | TRP |
| V-BETA [D1] | FR2 | 42V | 35 | 33 | Y | TYR |
| V-BETA [D1] | FR2 | 43V | 36 | 34 | R | ARG |
| V-BETA [D1] | FR2 | 44V | 37 | 35 | Q | GLN |
| V-BETA [D1] | FR2 | 45V | 38 | 36 | D | ASP |
| V-BETA [D1] | FR2 | 46V | 39 | 37 | P | PRO |
| V-BETA [D1] | FR2 | 47V | 40 | 38 | G | GLY |
| V-BETA [D1] | FR2 | 48V | 41 | 39 | M | MET |
| V-BETA [D1] | FR2 | 49V | 42 | 40 | G | GLY |

**Supplementary Table S14. IMGT, Kabat and PDB numbering for TCR-Vβ .**
| IMGT DOMAIN | IMGT CDR | IMGT | Kabat | PDB (2F53) | Residue [1 letter] | Residue [3 letter] |
| --- | --- | --- | --- | --- | --- | --- |
| V-BETA [D1] | FR2 | 50V | 43 | 41 | L | LEU |
| V-BETA [D1] | FR2 | 51V | 44 | 42 | R | ARG |
| V-BETA [D1] | FR2 | 52V | 45 | 43 | L | LEU |
| V-BETA [D1] | FR2 | 53V | 46 | 44 | I | ILE |
| V-BETA [D1] | FR2 | 54V | 47 | 45 | H | HIS |
| V-BETA [D1] | FR2 | 55V | 48 | 46 | Y | TYR |
| V-BETA [D1] | CDR2 | 56V | 49 | 47 | S | SER |
| V-BETA [D1] | CDR2 | 57V | 50 | 48 | V | VAL |
| V-BETA [D1] | CDR2 | 58V | 51 | 49 | S | SER |
| - | CDR2 | 59 | - | - | - | - |
| - | CDR2 | 60 | - | - | - | - |
| - | CDR2 | 61 | - | - | - | - |
| - | CDR2 | 62 | - | - | - | - |
| V-BETA [D1] | CDR2 | 63V | 52 | 50 | V | VAL |
| V-BETA [D1] | CDR2 | 64V | 53 | 51 | G | GLY |
| V-BETA [D1] | CDR2 | 65V | 54 | 52 | M | MET |
| V-BETA [D1] | FR3 | 66V | 55 | 53 | T | THR |
| V-BETA [D1] | FR3 | 67V | 56 | 54 | D | ASP |
| V-BETA [D1] | FR3 | 68V | 57 | 55 | Q | GLN |
| V-BETA [D1] | FR3 | 69V | 58 | 56 | G | GLY |
| V-BETA [D1] | FR3 | 70V | 59 | 57 | E | GLU |
| V-BETA [D1] | FR3 | 71V | 60 | 58 | V | VAL |
| V-BETA [D1] | FR3 | 72V | 61 | 59 | P | PRO |
| - | FR3 | 73 | 62 | - | - | - |
| V-BETA [D1] | FR3 | 74V | 63 | 60 | N | ASN |
| V-BETA [D1] | FR3 | 75V | 64 | 61 | G | GLY |
| V-BETA [D1] | FR3 | 76V | 65 | 62 | Y | TYR |
| V-BETA [D1] | FR3 | 77V | 66 | 63 | N | ASN |
| V-BETA [D1] | FR3 | 78V | 67 | 64 | V | VAL |
| V-BETA [D1] | FR3 | 79V | 68 | 65 | S | SER |
| V-BETA [D1] | FR3 | 80V | 69 | 66 | R | ARG |
| V-BETA [D1] | FR3 | 81V | 70 | 67 | S | SER |
| - | FR3 | 82 | - | - | - | - |
| V-BETA [D1] | FR3 | 83V | 71 | 68 | T | THR |
| V-BETA [D1] | FR3 | 84V | 72 | 69 | T | THR |
| V-BETA [D1] | FR3 | 85V | 73 | 70 | E | GLU |
| V-BETA [D1] | FR3 | 86V | 74 | 71 | D | ASP |
| V-BETA [D1] | FR3 | 87V | 75 | 72 | F | PHE |
| V-BETA [D1] | FR3 | 88V | 76 | 73 | P | PRO |
| V-BETA [D1] | FR3 | 89V | 77 | 74 | L | LEU |
| V-BETA [D1] | FR3 | 90V | 78 | 75 | R | ARG |
| V-BETA [D1] | FR3 | 91V | 79 | 76 | L | LEU |
| V-BETA [D1] | FR3 | 92V | 80 | 77 | L | LEU |
| V-BETA [D1] | FR3 | 93V | 81 | 78 | S | SER |
| V-BETA [D1] | FR3 | 94V | 82 | 79 | A | ALA |
| V-BETA [D1] | FR3 | 95V | 83 | 80 | A | ALA |
| V-BETA [D1] | FR3 | 96V | 84 | 81 | P | PRO |
| V-BETA [D1] | FR3 | 97V | 85 | 82 | S | SER |
| V-BETA [D1] | FR3 | 98V | 86 | 83 | Q | GLN |

**Supplementary Table S14. IMGT, Kabat and PDB numbering for TCR-Vβ .**
| IMGT DOMAIN | IMGT CDR | IMGT | Kabat | PDB (2F53) | Residue [1 letter] | Residue [3 letter] |
| --- | --- | --- | --- | --- | --- | --- |
| V-BETA [D1] | FR3 | 99V | 87 | 84 | T | THR |
| V-BETA [D1] | FR3 | 100V | 88 | 85 | S | SER |
| V-BETA [D1] | FR3 | 101V | 89 | 86 | V | VAL |
| V-BETA [D1] | FR3 | 102V | 90 | 87 | Y | TYR |
| V-BETA [D1] | FR3 | 103V | 91 | 88 | F | PHE |
| V-BETA [D1] | FR3 | 104V | 92 | 89 | C | CYS |
| V-BETA [D1] | CDR3 | 105V | 93 | 90 | A | ALA |
| V-BETA [D1] | CDR3 | 106V | 94 | 91 | S | SER |
| V-BETA [D1] | CDR3 | 107V | 95 | 92 | S | SER |
| V-BETA [D1] | CDR3 | 108V | 96 | 93 | Y | TYR |
| V-BETA [D1] | CDR3 | 109V | - | 94 | V | VAL |
| V-BETA [D1] | CDR3 | 110V | - | 95 | G | GLY |
| - | CDR3 | 111 | - | - | - | - |
| V-BETA [D1] | CDR3 | 112V | - | 96 | N | ASN |
| V-BETA [D1] | CDR3 | 113V | - | 97 | T | THR |
| V-BETA [D1] | CDR3 | 114V | - | 98 | G | GLY |
| V-BETA [D1] | CDR3 | 115V | - | 99 | E | GLU |
| V-BETA [D1] | CDR3 | 116V | - | 100 | L | LEU |
| V-BETA [D1] | CDR3 | 117V | - | 101 | F | PHE |
| V-BETA [D1] | FR4 | 118V | - | 102 | F | PHE |
| V-BETA [D1] | FR4 | 119V | - | 103 | G | GLY |
| V-BETA [D1] | FR4 | 120V | - | 104 | E | GLU |
| V-BETA [D1] | FR4 | 121V | - | 105 | G | GLY |
| V-BETA [D1] | FR4 | 122V | - | 106 | S | SER |
| V-BETA [D1] | FR4 | 123V | - | 107 | R | ARG |
| V-BETA [D1] | FR4 | 124V | - | 108 | L | LEU |
| V-BETA [D1] | FR4 | 125V | - | 109 | T | THR |
| V-BETA [D1] | FR4 | 126V | - | 110 | V | VAL |
| V-BETA [D1] | FR4 | 127V | - | 111 | L | LEU |

